# The Sugar Porter gene family of *Piriformospora indica*: Nomenclature, Transcript Profiling and Characterization

**DOI:** 10.1101/2021.03.11.434983

**Authors:** Sumit Raj, Manoj Kumar, Alok Kumar Singh, Meenakshi Dua, Atul Kumar Johri

## Abstract

*Piriformospora indica* is one of the prominent mutualistic root endophyte known to overcome phosphate and nitrogen limitation in a wide variety of plant species, reciprocally takes up carbohydrates for its survival and growth. A total of nineteen potential hexose transporters have been identified from *P. indica* genome, that may contributes to its potential of carbohydrate assimilation from host plant. Phylogenetic analysis assembles it in 10 groups within 3 clusters. To ease the study, systematic nomenclature were provided to 19 putative hexose transporters as “PiST1-PiST19” in accordance to their appearance on the supercontigs genome sequence of *P. indica*. The protein length ranges from 487 to 608 amino acids. Out of 19 putative hexose transporters, 9 have been predicted to contain 12 transmembrane domains (PiST1, PiST2, PiST5, PiST6, PiST9, PiST10, PiST11, PiST12 and PiST17), along with MFS family and Sugar porter subfamily motif. Therefore, transcripts were detected for these 9 genes. During colonization, three *P. indica* genes PiST1, PiST5 and PiST9 have shown induction as compared to axenic culture. Similarly during phosphate starvation, revealed PiST12 to be strongly enhanced. Carbon starvation study in liquid axenic culture resulted in induction of 4 genes, PiST6, PiST9, PiST12 and PiST17. We found co-relation in the expression pattern of PiPT and PiST12 during phosphate starvation. *In silico* analysis revealed the presence of functional conserved fucose permease (FucP) domain, involved in fructose transport. Phylogenetic analysis revealed that PiST12 groups closely with basidiomycetes hexose transporters. Further, functional complementation of Δ*hxt* null mutant revealed, PiST12 is able to complement growth on fructose and galactose but negligible on glucose.

## Introduction

To develop sustainable agricultural system, a major research focus is on symbiotic organism especially Arbuscular mycorrhizal fungus (AMF). The symbiosis between plants and AMF is arguably the world’s most prevalent mutualism. The vast majority of land plants form AM interactions, in which plants supply associated AMF with carbohydrates, essential for fungal survival and growth^1^. In exchange, AMF provide their host plants with mineral nutrients [e.g., phosphorus (P)] and other benefits such as protection against biotic (pathogens and herbivores) and abiotic (e.g., drought) stresses^2^. This partnership is credited with driving the colonization of land by plants, enabling massive global nutrient transfer and critical carbon sequestration^2,3^. In mycorrhizal association, the root and fungus does not function independently, but form a unit with specific metabolic pathways and regulated exchange of metabolites^4,5^. It is generally established that there is reciprocal benefit to the partners, due to the exchange of plant-derived carbohydrates for amino acids and nutrients supplied by the fungus^6,7^. It has been proposed that exchange across the root-fungus interface generally involve the passive diffusion of P_i_ and carbohydrates through the fungal and plant plasma membranes into the interfacial apoplast and then the active absorption of nutrients by both partners driven by H^+^-ATPase(s)^8^. However, how carbohydrates and phosphate are transported through the symbiotic interface is still unknown. Recently, glomeromycotan monosaccharide transporter, GpMST1, by exploiting the unique symbiosis of a glomeromycotan fungus (*Geosiphon pyriformis*) with cyanobacteria and another high-affinity Monosaccharide Transporter 2 (MST2) from *Glomus sp.* with a broad substrate spectrum” that functions at several symbiotic root locations have been characterized^9,10^.

Carbon supply to fungus and P uptake are reciprocally dependent on each other. Several workers have shown decrease in photosynthesis leads to a reduction of the mycorrhizal growth response and of the P uptake by AM systems^11^. Presently, we know very little about the regulation of exchange processes occurring in a mycorrhiza and the mechanisms involved in polarizing the transfers due to lack of pure culture and genetic manipulation. It is thus assumed that sucrose is delivered into the apoplast at the plant–fungus interface and hydrolyzed via a plant-derived acid invertase^12,13^. The resulting hexoses are then taken up by the fungus^1^. *Piriformospora indica* has been involved as model fungus to study plant-fungus interaction, as it can be axenically cultured and genetically manipulated to study plant-fungal interaction due to recently developed stable transformation system^14^. Similar to AMF, it forms arbuscules like coiled structure while growing progressively inter- and intra-cellularly^15,16^. The bidirectional nutrient exchange occurs at the apoplastic space enclosed by periarbuscular membrane of the host plant and fungal cell wall^17–22^. However, the carbohydrate substrate as well as its mechanism of transportation during symbiosis is not deciphered yet.

*Saccharomyces cerevisiae* have revealed more than 20 sugar transporters^23^, only few carbohydrate transporters (two from *A. muscaria*^24,25^ and one from *Tuber borchii*^26^) have been characterized from EM fungi yet. From *Laccaria bicolor* genome, 15 members of sugar porter family have been identified. Out of which, 3 (LbMST1.2, LbMST1.3, and LbMST3.1) are investigated as high-affinity glucose transporters^27^. In AMF, only 2 hexose transporters (GpMST1 from symbiotic fungus *Geosiphon pyriformis*^9^ and MST2 from *Glomus intraradices*^10^) are characterized as high affinity transporters with broad substrate specificity. Recently, we have functionally characterized a high affinity hexose transporter with broad substrate specificity *PiHXT5* from *P. indica*^28^. Therefore, investigation of fungal hexose transporters are in initial stages and a large gap has to be filled.

With the genome sequencing of *P. indica*^29^, revealing 19 putative hexose transporters, their further in depth systematic study become necessity. The availability of a battery of putative hexose transporter genes indicates different transporters respond to different stresses and conditions. Here we are systematically reporting their location on genome, functional domains, nomenclature, exon-intron boundary, transmembrane domain. Further we have investigated the expression of nine members of SP gene family that has predicted 12 transmembrane domains as characteristics of majority of MFS members during colonization in maize and axenic conditions. Further, our expression analysis revealed that PiSTP12 (accession number CCA69469.1) is a phosphate starvation responsive hexose transporter in *P. indica*. Till date, no report is available on phosphate starvation induced hexose transporters in mutualistic endophyte *P. indica*. The characterization of phosphate responsive hexose transporter is critical in fungus to decipher its better phosphate scavenging ability during Pi starvation that requires higher photosynthate transfer, better colonization, extensive hyphal growth and deeper soil penetration. We found corelation in the expression pattern of high affinity phosphate transporter PiPT and PiST12 during phosphate starvation. Further we have isolated and functionally complemented putative hexose transporter PiST12. *In silico* analysis revealed the presence of functional conserved fucose permease (FucP) domain, involved in fructose transport. Phylogenetic analysis revealed that PiST12 groups closely with basidiomycetes hexose transporters. Complementation study has given an insight to its unique substrate specificity. PiST12 complemented Δ*hxt null* mutant EBY.VW4000 revealed maximum growth on fructose, followed by galactose and negligible on glucose.

## Results

### Identification and Phylogenetic relationships of *P. indica* sugar porter genes

Initially, we have identified 19 potential hexose transporters of *P. indica* **(Table 1)**, for which phylogenetic analysis was performed. For 19 putative hexose transporters of *P. indica*, location on supercontigs of genome sequence has been identified. *PiST10, PiST11* and PiST12 are located adjacently PIRI_contig_0060. *PiST10* and *PiST12* have high query coverage (100%), identities (89%), similarity (96%) and minimum E-value (0.00) and same number of predicted exons (4). Similarly, *PiST11* and *PiST12* have high query coverage (100%), identities (86%), similarity (92%) and minimum E-value (0.00) and similar number of predicted exons. This suggests *PiST10, PiST11* and *PiST12* have originated from gene duplication during evolution **(Table 1)**. Similarly, *PiST9* and *PiST6* have high query coverage (100%), identities (80%), similarity (90%) and minimum E-value (0.00). Though *PiST1* and *PiST2* are located adjacently on PIRI_contig_0004 in same direction and have high query coverage (91%) and E-value (0.00) but low identity (65%). Members of sugar porter transcript length ranges from 1,464bp (487aa) to 1,827 bp (608 aa) possessing MFS domain (TIGR00879) and SP domain (pfam00083) **(Table 2)**. Contrary, PiST3 have only 1,065bp (354aa) and may be truncated. Phylogenetic analysis assembles sugar porters into 10 groups and 3 cluster as analyzed by bootstrap values (>60%) **(Fig. 1)**. Single SP protein with Accession number CCA67083.1 was classified as lactose permease is not included in phylogenetic analysis. Few members in each (cluster II and cluster III) turned out to be not only highly homologous regarding their protein sequences but also physically linked on a single contig (**Table S1).**

**Table 1:** Systemic Nomenclature of the Sugar Porter gene family of *P. indica*. **Nomenclature of Sugar Porter gene Family of *P.indica***. The recent identification of hexose transporters from *P. indica* created the necessity for systemic nomenclature for study of 19 hexose transporters in *P. indica* genome to avoid confusion in future by different groups. So, we suggest that, since all the sugar transporters identified from *P. indica* possess Sugar Transporter Superfamily Acc. No. pfam00083 domain, so the letter “ST” provides the summary for the members belonging to this family. Further, as it belongs to *P. indica* genome so “ST” is pre-suffixed with “Pi” to form “PiST”. The numbering of the genes is assigned in accordance to their appearance in the Supercontigs genome sequence. Therefore, the letter PiST1 provides the insight for the gene that it belongs to which organism, protein family and their sequential appearance in contig.

| Sr. No. | Protein | Alias Gene Name | Location on Supercontig | Strand | Accession No. Transcript | Gene esembl | Transcript length (bps) | Translation Length (aa) | Coding Exons |
| --- | --- | --- | --- | --- | --- | --- | --- | --- | --- |
| 1. | PiST1 | PiHXT1 | PIRI_contig_0004:83250-85641 | Reverse | CCA66651.1 | PIIN_00334 | 1,689 | 562 | 14 |
| 2. | PiST2 | PiHXT6 | PIRI_contig_0004:87021-89326 | Reverse | CCA66652.1 | PIIN_00335 | 1,827 | 608 | 10 |
| 3. | PiST3 |  | PIRI_contig_0004:91385-93760 | Reverse | CCA66654.1 | PIIN_00337 | 1,065 | 354 | 7 |
| 4. | PiST4 |  | PIRI_contig_0010:167941-170511 | Forward | CCA67037.1 | PIIN_11722 | 1,464 | 487 | 20 |
| 5. | PiST5 | PiHXT7 | PIRI_contig_0011:96026-98147 | Forward | CCA67083.1 | PIIN_00918 | 1,644 | 547 | 10 |
| 6. | PiST6 | PiHXT9 | PIRI_contig_0023:119026-120800 | Reverse | CCA67897.1 | PIIN_01718 | 1,620 | 539 | 4 |
| 7. | PiST7 |  | PIRI_contig_0025:70629-72954 | Forward | CCA67930.1 | PIIN_01799 | 1,818 | 605 | 10 |
| 8. | PiST8 |  | PIRI_contig_0032:27939-30622 | Reverse | CCA68264.1 | PIIN_02129 | 1,557 | 518 | 22 |
| 9. | PiST9 | PiHXT8 | PIRI_contig_0047:71992-73950 | Reverse | CCA68995.1 | PIIN_02855 | 1,614 | 537 | 6 |
| 10. | PiST10 | PiHXT2 | PIRI_contig_0060:97-1833 | Forward | CCA69467.1 | PIIN_03367 | 1,584 | 527 | 4 |
| 11. | PiST11 | PiHXT4 | PIRI_contig_0060:2971-4710 | Forward | CCA69468.1 | PIIN_03368 | 1,539 | 512 | 5 |
| 12. | PiST12 | PiHXT3 | PIRI_contig_0060:5980-7748 | Reverse | CCA69469.1 | PIIN_03369 | 1,593 | 530 | 4 |
| 13. | PiST13 |  | PIRI_contig_0068:31692-34311 | Reverse | CCA69766.1 | PIIN_03707 | 1,704 | 567 | 12 |
| 14. | PiST14 |  | PIRI_contig_0087:34856-37715 | Reverse | CCA70422.1 | PIIN_04361 | 1,656 | 551 | 24 |
| 15. | PiST15 |  | PIRI_contig_0090:12283-14503 | Reverse | CCA70504.1 | PIIN_04441 | 1,665 | 554 | 8 |
| 16. | PiST16 |  | PIRI_contig_0106:12726-14833 | Reverse | CCA70963.1 | PIIN_04898 | 1,557 | 518 | 11 |
| 17. | PiST17 | PiHXT5 | PIRI_contig_0115:16420-18536 | Forward | CCA71201.1 | PIIN_05137 | 1,578 | 525 | 12 |
| 18. | PiST18 |  | PIRI_contig_0232:8886-11354 | Forward | CCA73450.1 | PIIN_07404 | 1,530 | 509 | 20 |
| 19. | PiST19 |  | PIRI_contig_0288:19943-22204 | Forward | CCA74136.1 | PIIN_08090 | 1,707 | 568 | 11 |

**Table 2:** Major Domain Hits of the Sugar Porter gene family of *P. indica*. **List of Transporter Domains present in each member of Sugar Porter Family of *P. indica***. To predict the substrate specificity of different members of Sugar Porter family, the domains were identified. Presence of Major Facilitator Domain, Sugar Porter Domain indicates their family. Further substrate specific domains like D-Xylose permease, Fucose permease suggested their substrate specificity.

| Table 2: Major Domain Hits of the Sugar Porter gene family of *P. indica* |  |  |  |  |  |  |  |
| --- | --- | --- | --- | --- | --- | --- | --- |
| Sr. No. | Protein | Accession Number | Major Facilitator Superfamily (MFS) Acc. No. TIGR00879 |  | Sugar Transporter Superfamily Acc. No. pfam00083 |  | Other Sugar-related Domains Present |
|  |  |  | Interval | E-value | Interval | E-value |  |
| 1. | PiST1 | CCA66651.1 | 21-472 | 1.63e-64 | 22-476 | 2.95e-76 |  |
| 2. | PiST2 | CCA66652.1 | 58-506 | 4.62e-59 | 62-510 | 1.15e-66 |  |
| 3. | PiST3 | CCA66654.1 | 156-344 | 2.09e-08 | 174-344 | 4.29e-09 | Putative S-adenosyl-L-methionine-methyltransferase |
| 4. | PiST4 | CCA67037.1 | 51-434 | 4.24e-87 | 51-438 | 2.71e-88 |  |
| 5. | PiST5 | CCA67083.1 | 62-514 | 1.50e-78 | 64-518 | 1.78e-82 |  |
| 6. | PiST6 | CCA67897.1 | 24-482 | 1.54e-130 | 26-486 | 2.24e-132 |  |
| 7. | PiST7 | CCA67930.1 | 152-548 | 1.21e-82 | 153-552 | 1.37e-89 |  |
| 8. | PiST8 | CCA68264.1 | 29-465 | 1.10e-88 | 35-469 | 3.15e-89 |  |
| 9. | PiST9 | CCA68995.1 | 13-481 | 2.71e-127 | 30-485 | 1.28e-130 |  |
| 10. | PiST10 | CCA69467.1 | 29-481 | 1.60e-69 | 39-485 | 1.05e-79 | Fucose permease FucP (COG0738) |
| 11. | PiST11 | CCA69468.1 | 49-466 | 7.28e-65 | 44-470 | 4.56e-78 | Fucose permease FucP (COG0738) |
| 12. | PiST12 | CCA69469.1 | 77-481 | 1.66e-70 | 44-485 | 1.09e-79 | Fucose permease FucP (COG0738) |
| 13. | PiST13 | CCA69766.1 | 133-558 | 2.38e-73 | 113-558 | 7.37e-76 | Glutaredoxin-like domain (DUF836) (pfam05768) |
| 14. | PiST14 | CCA70422.1 | 32-506 | 3.03e-104 | 36-510 | 4.46e-114 |  |
| 15. | PiST15 | CCA70504.1 | 6-470 | 1.55e-112 | 27-474 | 8.83e-116 |  |
| 16. | PiST16 | CCA70963.1 | 53-489 | 1.54e-84 | 57-493 | 1.67e-94 |  |
| 17. | PiST17 | CCA71201.1 | 28-473 | 2.91e-58 | 40-477 | 5.68e-69 |  |
| 18. | PiST18 | CCA73450.1 | 4-469 | 4.52e-80 | 9-473 | 7.52e-84 |  |
| 19. | PiST19 | CCA74136.1 | 37-476 | 3.40e-60 | 37-480 | 2.16e-64 |  |

**Figure 1.**
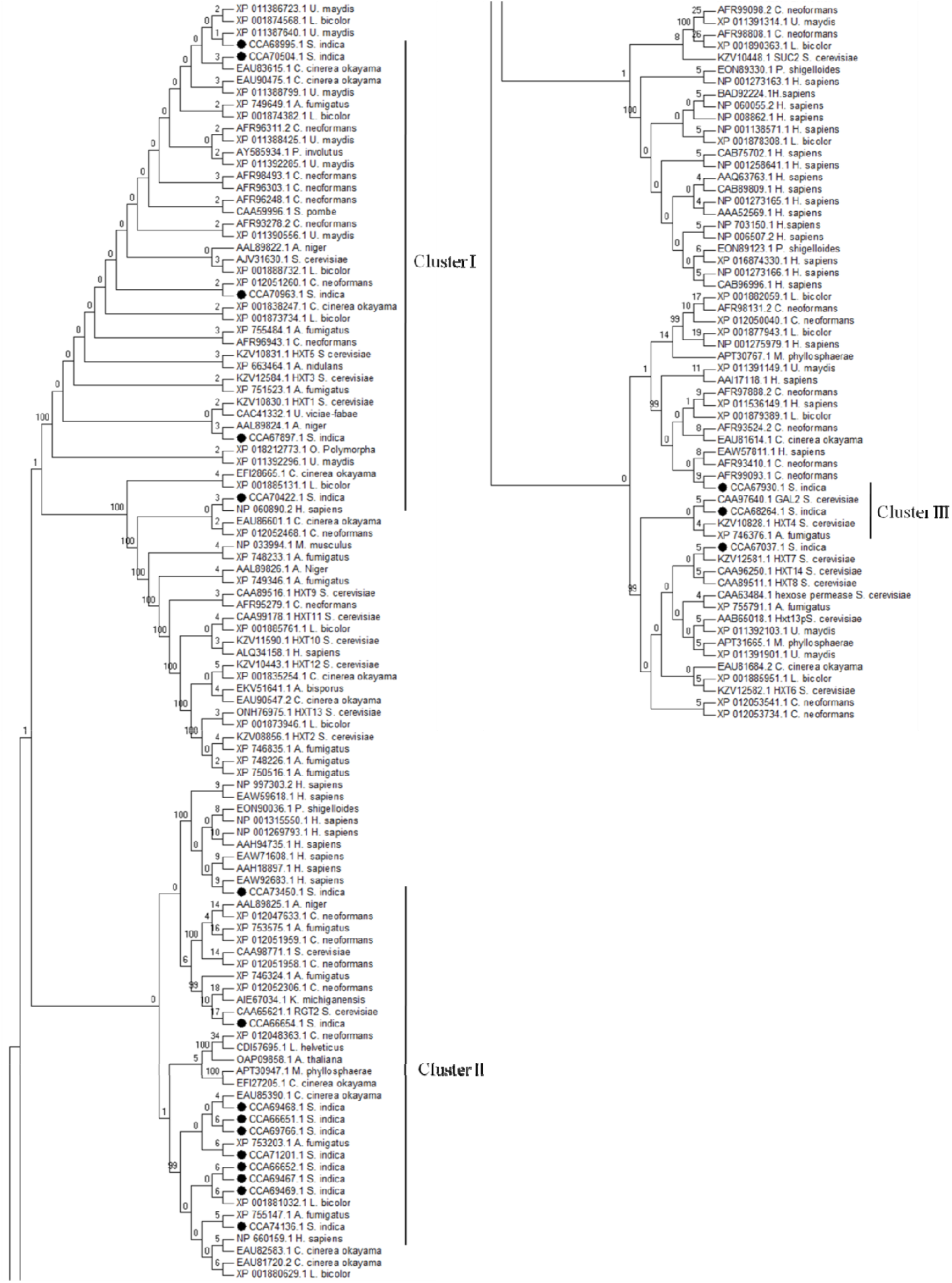
Phylogenetic relationships of the deduced protein sequences of the *Piriformospora indica* sugar porter gene family with known fungal hexose transporters. Putative hexose transporters deduced from the genome sequence of range of fungus, plants and animal kingdom including *Homo sapiens* which is enlisted in **Table S1**. Phylogenetic tree was constructed using MEGA6. The Phylogenetic relationships were obtained by maximum-likelihood method, using the JTT model of amino acid substitution. Number above branches indicates bootstrap values from 1000 replicates. Sugar porter gene family of *P. indica* was grouped into three clusters. The tree was unrooted. The phylogenetic tree was split into two parts for better visibility. Magnified view is also provided **(Figure S9 i-iii)**.

### Systemic Nomenclature of the Sugar Porter gene family of *P. indica*

The identification of sugar porters from *P. indica* leads to requirement of systemic nomenclature for study of 19 sugar porters in *P. indica* genome to avoid confusion in future by different groups **(Table 1)**. All identified Sugar Porter family members belongs to typical Major Facilitator Superfamily (MFS) (Acc. No. TIGR00879). MFS proteins typical length is 400–600 amino acids having duplicated TM topology **(Table 2)**. So, we suggest that, since sugar transporters identified from *P. indica* possess Sugar Transporter Superfamily Acc. No. pfam00083 domain, so the letter “ST” provides the summary for the members belonging to this family. Further, as it belongs to *P. indica* genome so “ST” is pre-suffixed with “Pi” to form “PiST”. The numbering of the genes is assigned in accordance to their appearance in the Supercontigs genome sequence. Therefore, the letter PiST1 provides the insight for the gene, which it belongs to which organism, protein family and their sequential appearance in contig. Here we also identified the direction of gene transcript on forward/reverse strand as well as number of exons that ranges from 4 to 24 **(Table 1)**. Additional monosaccharide transporter domain, Fucose permease FucP (COG0738) that is involved in Fucose and similar sugar derived from degradation of plant cell wall (fructose, galactose) were also identified **(Table 2).**

### Transmembrane Helices Prediction

To identify putative sugar porters, transmembrane (TM) domains, TM helix segments were predicted for all 19 Sugar Porter members. Finally nine sequences were identified with 12 transmembrane helices, with a long intracellular loop between helix 6 and 7 and cytosolic N and C-terminal **(Fig. 2).** These are the typical characteristics of MFS family.

**Figure 2.**
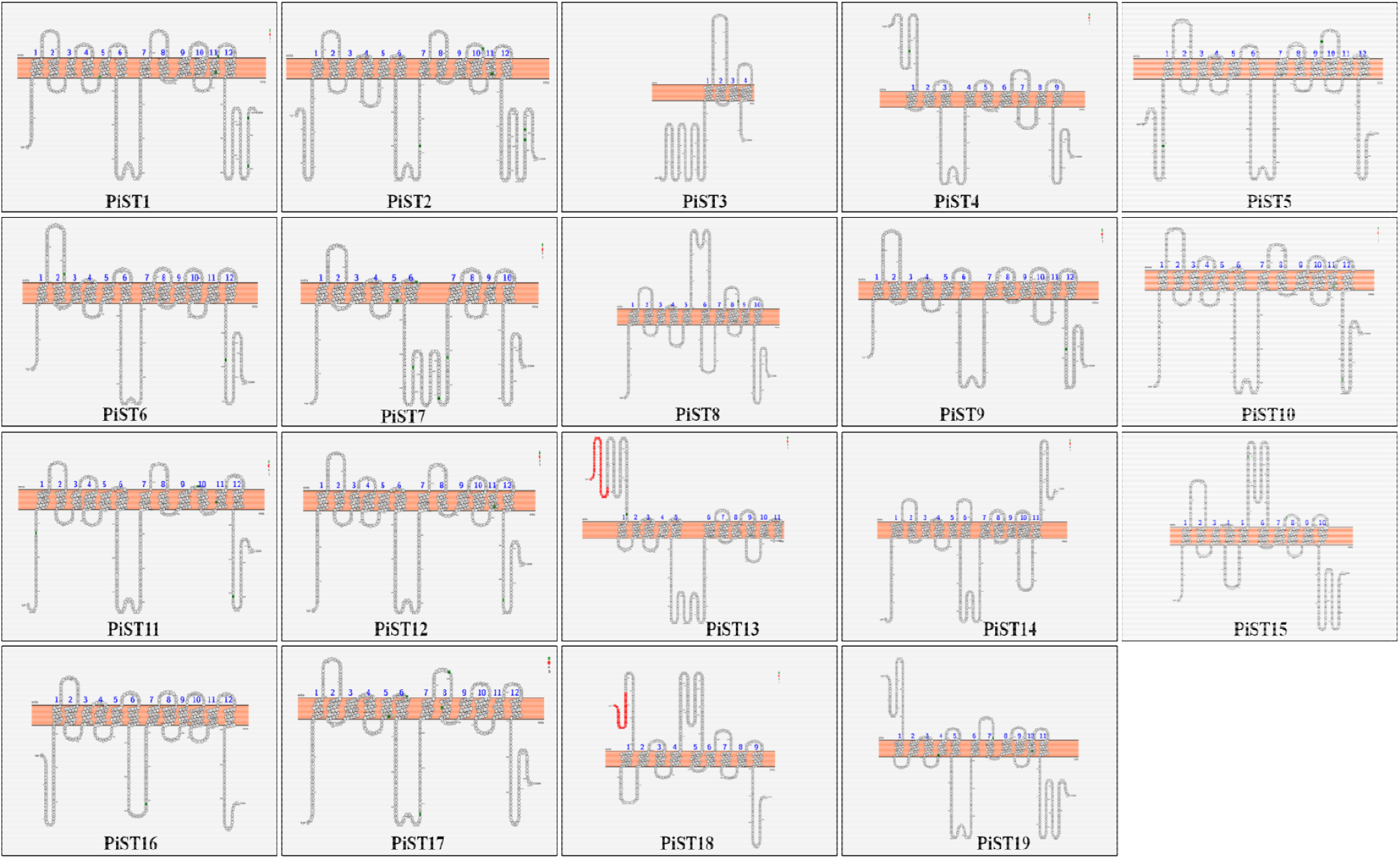
Prediction of Transmembrane domain of Sugar Porter family of *P. indica*: To predict the transmembrane domain and N- and C- face of sugar porter proteins, TM-Pred, HMM-TOP, SOSUI and PROTTER were applied. All 19 PiST were analyzed by this method and their topologies were obtained.

### *P. indica* hexose transporters are induced during mutualistic stage

Six genes (*PiST1, PiST5, PiST6, PiST12, PiST9 and PiST17*) showed enhanced transcript abundance upon colonization in maize plant when compared with the *P. indica* grown in axenic culture **(Fig. 3)**. One of them, *PiST17* showed more than 40 fold induction on colonization. *PiHXT10* has non-significant change in gene expression upon colonization. However, *PiST12* during colonization, revealed an enhanced expression level of putative hexose transporter genes compared with axenic culture.

**Figure 3.**
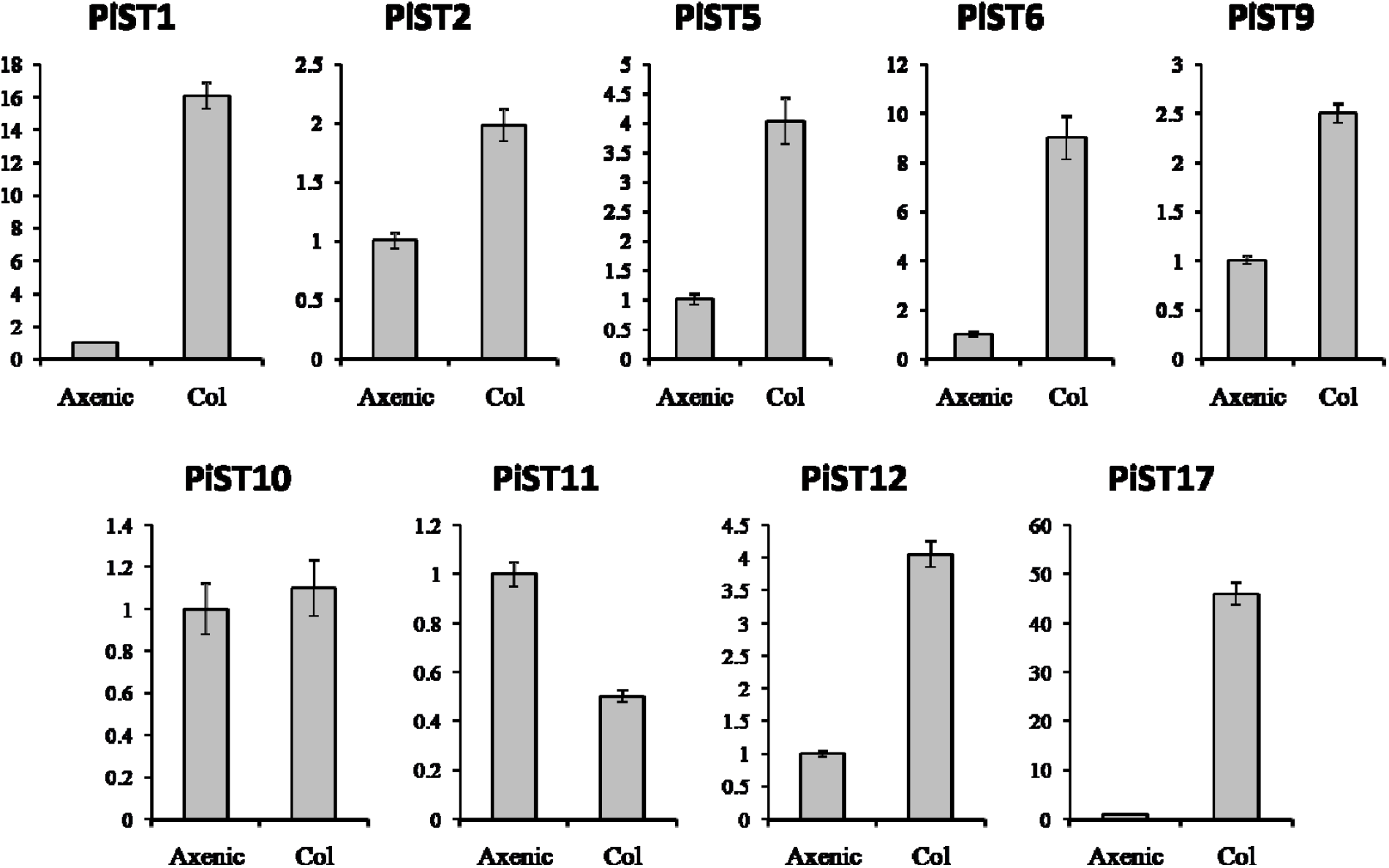
Expression analysis of hexose transporters for quantifying relative expression level in axenically grown and *P. indica* colonized with maize plant (Colonized). For the determination of transcript abundance of hexose transporters (*PiST1, PiST2, PiST5, PiST6, PiST9, PiST10, PiST11, PiST12, PiST17*), cDNA synthesis was performed from RNA obtained from axenic culture of *P. indica* and Maize root inoculated *P. indica*. Quantitative real-time analysis was performed with specific primers. Data were analyzed using comparative Ct method. *PiST17* is up regulated more than 40 folds among all analyzed *PiSTs*.

### Phosphate starvation induces the expression of putative sugar porter *PiST12*

Fungal carbohydrate and phosphate nutrition are interconnected and thus affect each other at the regulatory level. To look at the impact of phosphate nutrition on sugar porter’s transcript abundance, low (10uM) and high P (10mM) concentration were applied. Five out of 9 genes exhibited negligible or little change in their transcript abundances, while 4 genes get induced under Low P conditions. *PiST12* in particular induced more than 50 fold during phosphate starvation **(Fig. 4)**. *P. indica* induces higher fold enrichment in Biomass of Host Plant during phosphate deficiency as compared to P sufficient conditions. This putative fungal hexose transporter may be correlated with stronger carbohydrate sink strength at plant-fungal apoplastic interface, responsible for prominent rhizospheric structural development during phosphate starvation. Therefore, *PiST12* was selected for further study.

**Figure 4.**
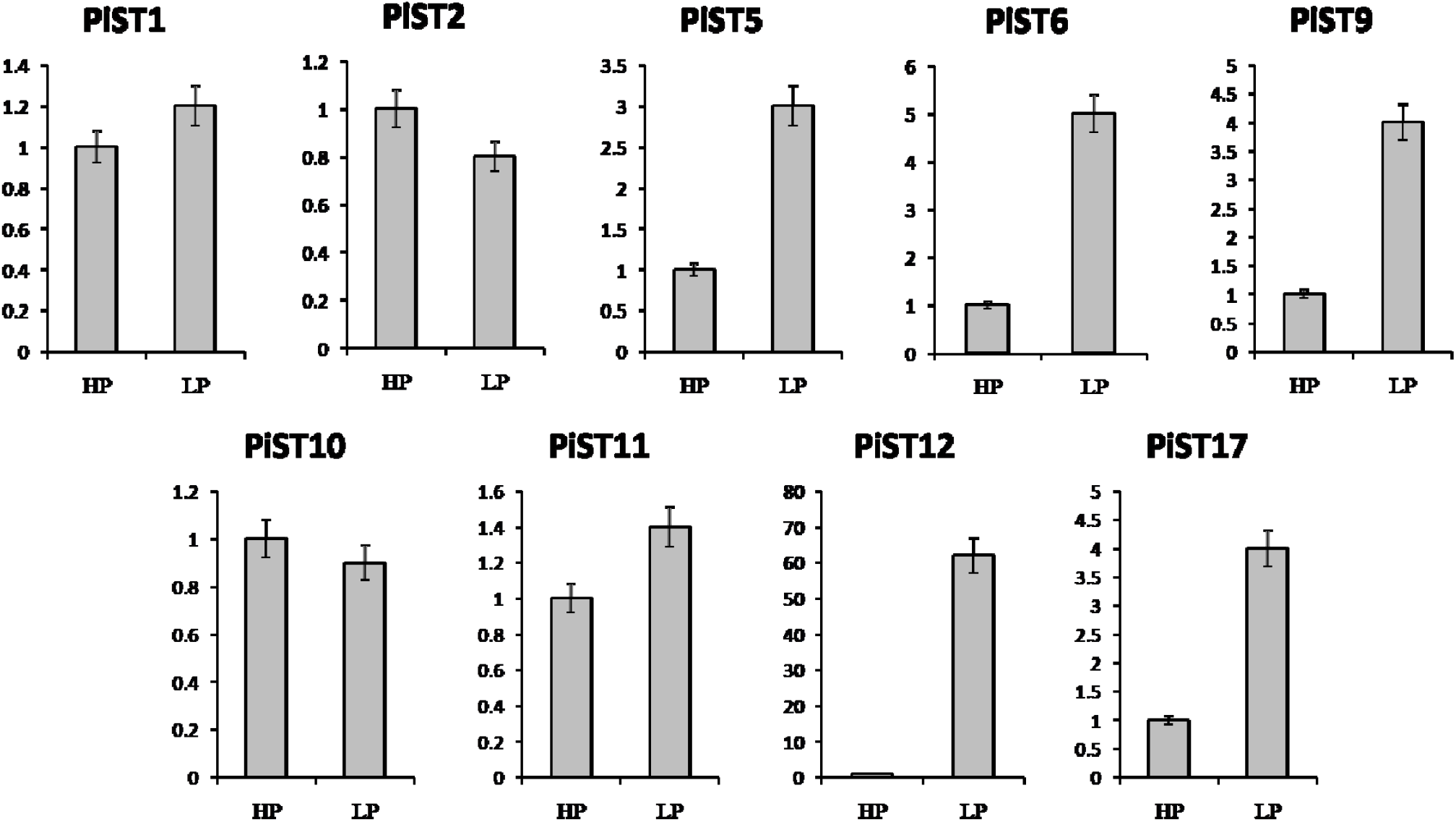
Expression analysis of PiSTs for quantifying relative expression level in *P. indica* colonized in maize plant at low P (10μM) and high P (10mM) for 15 days. For the determination of relative expression of hexose transporters (*PiST1, PiST2, PiST5, PiST6, PiST9, PiST10, PiST11, PiST12, PiST17*), RNA was isolated from *P. indica* colonized maize plant after 15 days of growth. cDNA was prepared and subjected to real time analysis. Quantitative realtime analysis was performed with specific primers. Data were analyzed using comparative Ct method. *PiST12* is up regulated more than 40 folds among all analyzed *PiSTs*.

### Induction of *PiST12* under Glucose starvation

*PiST6, PiST9, PiST12* and *PiST17* were found to be up regulated at 1mM or less glucose concentration. *PiST12* is found to be up regulated more than 40 fold, when no glucose is supplied in MN medium. Similarly, *PiST9* and *PiST6* were found to be up-regulated >20 fold, below 1 mM and *PiST12* is found 5.5-fold up regulated as compared to 111 mM glucose, respectively **(Fig. 5)**.

**Figure 5.**
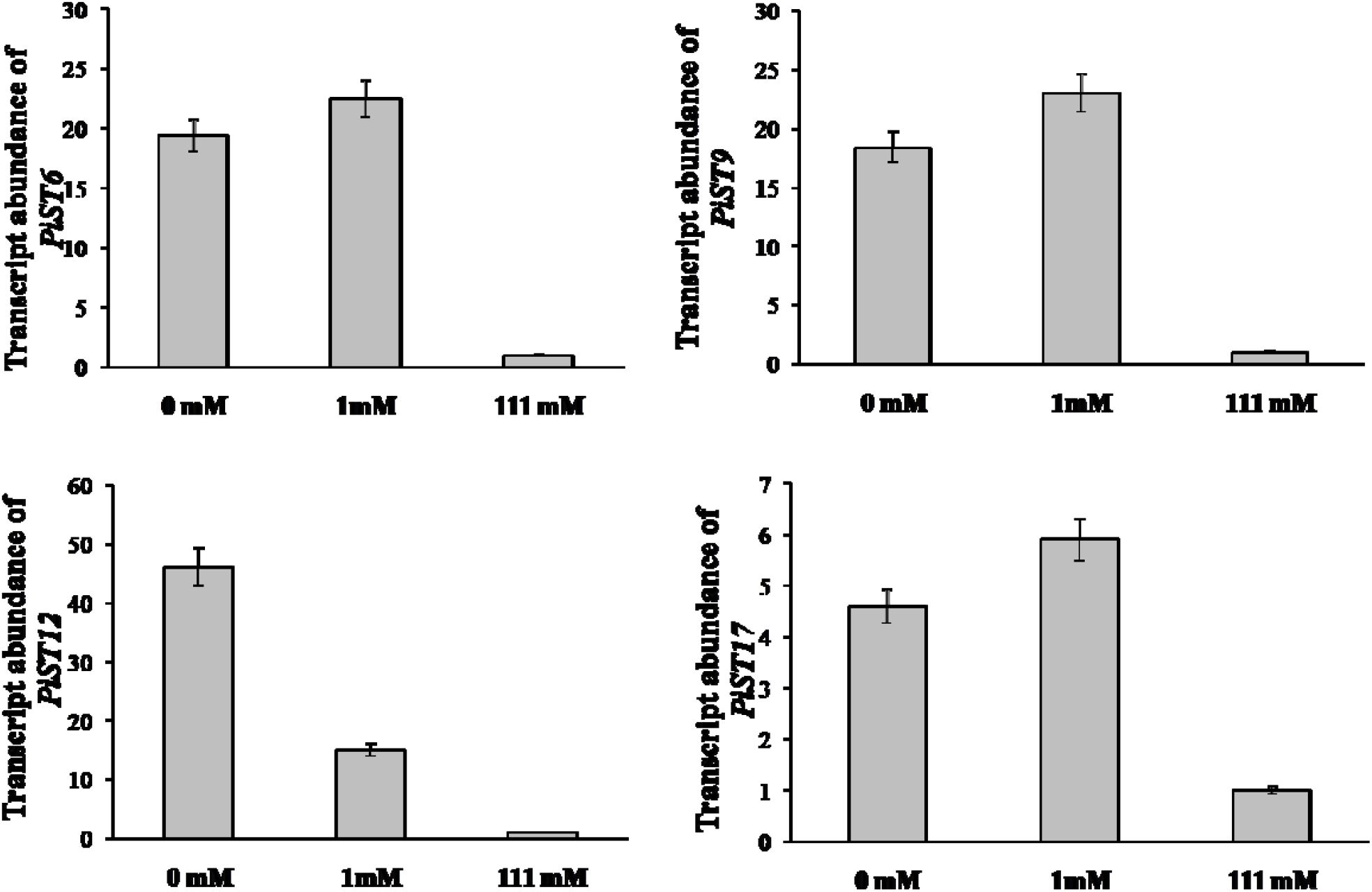
Expression analysis of PiSTs for quantifying relative expression level in axenically grown *P. indica* at different glucose concentration. For the determination of relative expression of *PiST6, PiST9, PiST12* and *PiST17, P. indica* is cultured axenically in modified Aspergillus minimal medium (AMM) and then washed 3 times with MN medium and then allowed to grow on MN medium for 5 more days containing different glucose concentration (0, 1mM and 111 mM). For the determination of transcript abundance of *PiSTs*, cDNA synthesis was performed from RNA obtained from axenic culture of *P. indica*. Quantitative real-time analysis was performed with specific primers. Data were analyzed using comparative Ct method. *PiHXT5* is up regulated more than 40 folds among all analyzed *PiSTs*.

### Multiple Sequence Alignment of hexose transporters

Nine sugar porters were aligned using MULTIALIN and CLUSTALW2, to identify conserved MFS and SP domains. Multiple sequence alignment shows presence of many conserved residues in these proteins. They all contain MFS and SP domains. *PiST5* showed non-conserved residues at critical positions. Two conserved GR-[KR] motifs, which are present in MFS transporters and the PETKG sequence, which is highly conserved among sugar transporters **(Fig. S1)**. *P. indica* putative sugar transporters show 17–60% homology with other sugar transporters. Invariant and highly conserved sugar porter family signature motifs [GR]-[P]-[PESPR]-[V-GR]-[LFP] - [PETKG] are found in *PiST1, PiST2, PiST5, PiST6, PiST9, PiST10, PiST11, PiST12, PiST17* to hGLUT1-3 and XylE with few exceptions **(Fig. S1)**. The structural and functional significance of these motifs has been examined previously. We have done comparative study of *P. indica* hexose transporter with *Homo sapiens* GLUT1. Recent crystallization of first eukaryotic glucose transporter GLUT1 from *Homo sapiens*, revealed critical residues involved in H-bonding with substrate at extracellular gate, sugar binding pocket and cytoplasmic release gate **(Fig. S2)** which are also conserved in PiST12. Expression analysis revealed the critical hexose transporters of *P. indica* that are regulated via phosphate signaling and colonization *i.e.*PiST6, PiST9, PiST12 and PiST17. Sequence alignment of PiST6, PiST9, PiST12, PiST17, hGLUT1, hGLUT2 and hGLUT3 by MULTALIN and CLUSTALW2, reveals conserved critical residues involved in glucose transport from extracellular to cytoplasmic side. Most of the residues are conserved in PiST6 and PiST9. Few residues are conserved in PiST12 and PiST17, critical for glucose transportation **(Fig. S2).**

### Domain identification of putative *PiST12* gene ORF

We have found that *PiST12* belongs to PIRI_contig_0060:5980-7748 (CCA69469.1) in *P. indica* genome. The isolated *PiST12* ORF (GenBank accession no. CCA69469.1) is 1593 bp long with ATG as a start and TAA as stop codon. Deduced amino acid sequence of putative *PiST12* protein possesses 530 amino acids and was predicted to have molecular weight of 63.6 KDa. BLASTX analysis of putative *PiST12* cDNA sequence demonstrated up to 75% identity with other known hexose transporter amino acid sequences. **(Table 1)**

Protein BLAST (BLASTp) conserved domains analysis **(Fig. S3)** of CCA69469.1. showed that it is harboring Major Facilitator Superfamily (MFS) domain, Sugar Transporter Superfamily (SP) domain, Fucose domain involved in transport of Fucose and D-xylose transporter **(Table 2)**. Fucose is a hexose deoxy sugar with the chemical formula C_6_H_12_O_5_. It is found on N-linked glycans on plant cell surface and is the fundamental sub-unit of the fucoidan polysaccharide (α-1→3) linked core fucose. It is equivalent to 6-deoxy-L-galactose.

### Cloning of putative PiST12 gene and sequencing

After identification, putative *PiST12* ORF was PCR amplified with PiST12 specific primers from cDNA of low P condition. We have amplified a 1.593 kb fragment and this was cloned in pGEM-T easy vector and transformed in *E. coli*. Positive clones were confirmed by colony PCR, restriction digestion and sequenced. Our restriction digestion analysis confirmed the presence of 1.593 kb fragment in the transformed positive clones. BLASTX analysis of putative PiST12 cDNA sequence demonstrated up to 75% identity with other known HXTs.

The isolated *PiST12* ORF (GenBank accession no. CCA69469.1) is 1593 bp long with ATG as a start and TAA as stop codon **(Fig. S4)**. Deduced amino acid sequence of putative *PiST12* protein possesses 530 amino acids **(Fig. S5)** and was predicted to have molecular weight of 63.6 KDa. The gene posse’s exons interrupted by 3 introns in genome **(Fig. S6)**. BLASTX analysis of putative *PiST12* cDNA sequence demonstrated up to 75% identity with other known hexose transporter amino acid sequences.3D modeling was performed using Swiss Prot Template based modeling using GLUT3 as template **(Fig. S7)**. Further image adjustments were performed using Pymol.

### Homology and Phylogenetic and Molecular Evolutionary Analysis of *PiST12*

BlastX analysis showed 57–75% identity with many putative fungal hexose transporters. It has 23% identity with other hexose transporters such as GiMST2 (ADM21463.1), GpMST1 (CAJ77495.1), TbHXT1 (AAY26391), and ScHXT1 (AAA34700.1) **(Table S2)**. Phylogenetic analysis suggests that *PiST12* cluster together with uncharacterized fungal hexose transporters from *Auricularia delicate* (EJD41559.1), *Trametes versicolor* (EIW61310.1), *Coprinopsis cinerea okayama* (XP001839222.1), *Paxillus involutus* (AAT91304.1), and *Aspergillus flavus* (XP002374068.1). Multiple sequence alignment with known fungal hexose transporters showed high similarity **(Fig. 6)**. These branches include closely related hexose transporters from Basiodiomycota or Ascomycota fungi **(Fig. S7)**. Further, our additional phylogenetic analysis with diverse groups such as insects, mammals and plants homologs showed that *PiST12* stands among hexose transporters from Basidiomycete fungi **(Fig. 7)**.

**Figure 6.**
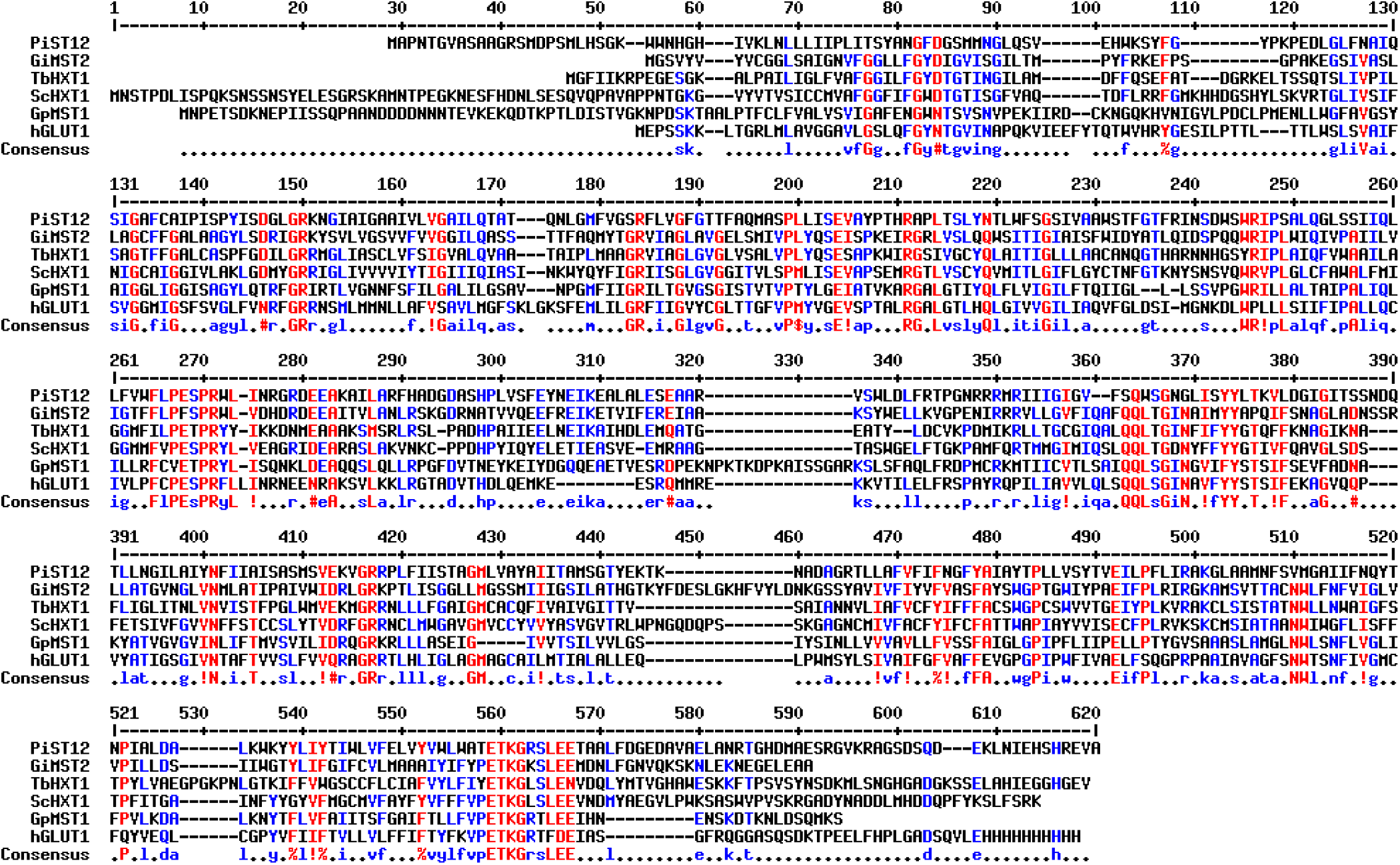
Alignment of the predicted amino acid sequence of PiST12. PiST12 is aligned with *Glomusintraradices* (GiMST2), *Tuber borchii* (TbHXT1), *Saccaharomyces cerevisiae* (ScHXT1) and *Geosiphon pyriformis*(GpMST1) and *Homo sapiens* (hGLUT1) by using MULTIALIN. The degree of sequence conservation at each position amino acids is shown in red, and low consensus amino acids are shown in blue.

**Figure 7.**
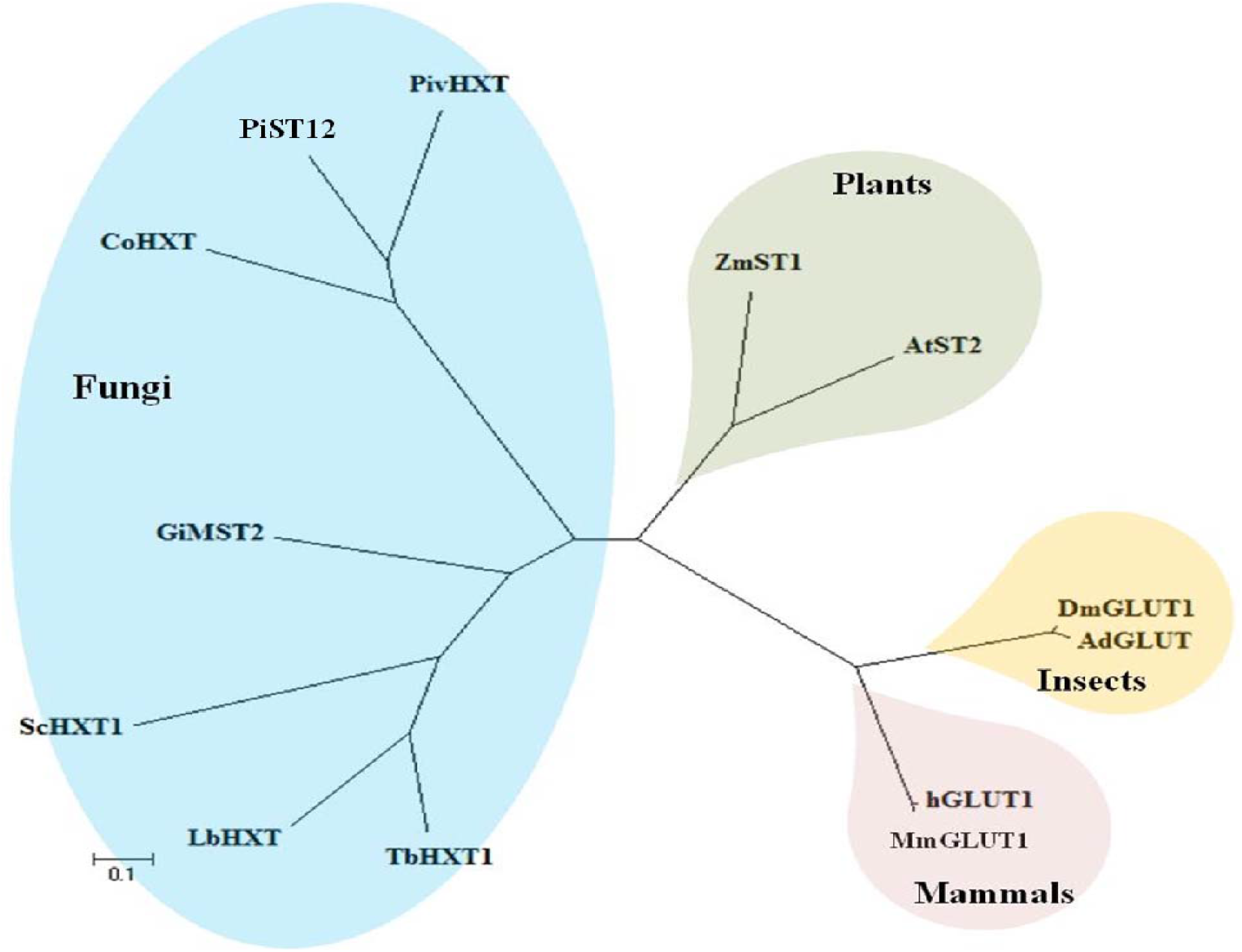
Unrooted phylogenetic relationship of PiST12 with other high affinity hexose transporters from plants and fungi. Protein names with GenBank™ (GB) accession numbers are listed **Table 3**. The evolutionary history was inferred using the Neighbor-Joining method. The tree is drawn to scale, with branch lengths in the same units as those of the evolutionary distances used to infer the phylogenetic tree. Phylogenetic analyses were done by using MEGA6.

### Complementation Assay

In order to functionally characterize *PiST12*, it was cloned in yeast expression vector p112A1NE under the control of ADH1 promoter. This construct p112A1NE-*PiST12* was transformed into *S. cerevisiae* strain EBY.VW4000. Because EBY.VW4000 is devoid of any hexose transporter, its growth was found retarded on glucose, as compare to EBY.VW4000 complemented with *PiST12* or its parental strain harboring only p112A1NE plasmid vector. Growth of EBY.VW4000 transformed with p112A1NE-*PiST12* construct or empty expression vector and untransformed EBY.VW4000 was compared in YNB medium containing all the amino acids except tryptophan. *PiST12able* to complement growth on fructose and galactose, whereas micro colonies were also observed on glucose as compared to WT and p112A1NE vector transformed EBY.VW4000 strains **(Fig. 8)**.

**Figure 8.**
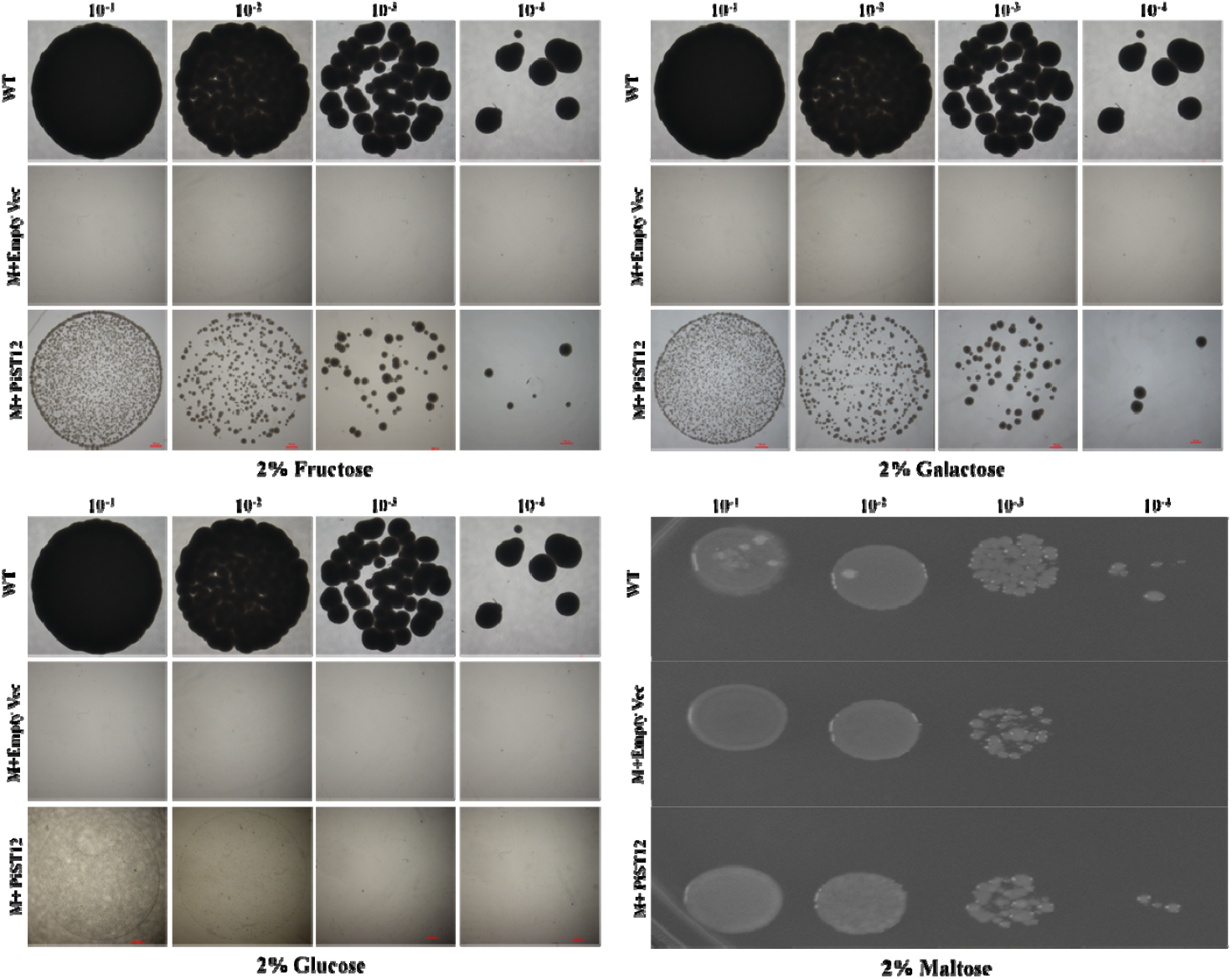
Complementation experiments using the *PiST12* in *S. cerevisiae* Δhxt null mutant strain EBY.VW4000 (A) 2% Fructose: (B) 2% Galactose: (C) 2% Glucose (D) 2% Maltose: The three yeast strains used were: wild type (wt), mutant yeast strain EBY.VW4000 carrying the empty p112A1NE plasmid (Δ*hxt* null + p112A1NE) and mutant yeast strain EBY.VW4000 carrying the p112A1NE plasmid with *PiST12* under the ADH promoter (Δhxt null + p112A1NE-*PiST12*) Freshly streaked cells were suspended in normal saline (0.9% NaCl) to an optical density at 600 nm (OD_600_) of 0.1 and 10-fold serial dilutions were made in 0.9% saline. 10 μ l was spotted on the plates with different carbon source and plates were incubated at 30 °C for 5–7 days to allow comparison between WT and mutant strains. Standard YPD media was used as control.

## Discussion

The translocation of sugar from the host phototrophs to heterotrophic fungus is the basis of symbiosis^30^. The carbohydrate transfer from cortical cells of host plant to fungal tissue occurs in three steps (1) Transfer of sucrose into apoplast (2) Sucrose degradation by host cell wall invertase into simpler monosaccharaides like fructose and glucose (3) Subsequent, uptake of monosacharides by fungal transporters into the intra-radical hyphae^15,31^. The underlying mechanism of carbohydrate translocation across symbiotic interface is still not understood. Monosaccharide transporters are involved in uptake of carbohydrate across plasma membrane^32^. Therefore, for symbiotic establishment as well as survival and maintenance of fungal partner, monosaccharide transporters at plant-fungal interface are crucial.

Presence of 19 putative hexose transporters in genome sequence of *P. indica* **(Table 1)** is similar to model organism *S. cerevisiae* as well as other fungal family like ascomycetous (*A. niger*) and basidiomycetous (*P. chrysosporium, C. cinerea* and *U. maydis*). In contrast, *C. neoformans* which is a basidiomycetous human pathogen posseses 48 predicted putative sugar porter genes. Members of sugar porter family of *P. indica* cluster in three groups in phylogenetic tree **(Fig. 1)**. PiST10, PiST11 and PiST12 have high query coverage (100%), identities (>85%), similarity (>91%) and minimum E-value (0.0), as well as, phylogenetic analysis showed clustering of PiST12 with PiST10 and PiST11, and located adjacently on same contig_0060 **(Fig. 1)**, suggest gene duplication during evolution. Such increase in copy number is related to nutrient stress to demonstrate greater fitness to increase its ability to scavenge more sugar at low substrate concentrations. Similar reports have been documented in *S.cerevisiae* HXT7/6^33^ and *H. sapiens* GLUT 14/3^34^ Similarly, PiST1, PiST2 and PiST3 located adjacently in PIRI_contig_0004 grouped in cluster II having high query coverage (>90%) and minimum E-value (0.0), but low identity (<68%). Cluster II harbors 9 GLUT proteins from *Homo sapiens*, which reflects its common evolution.

In all 19 members of sugar porter family in *P. indica*, MFS and SP domain has been identified **(Table 2)**, which attributes it to “sugar porter” family found in organism from microbes to man, which further belongs to bigger major facilitator superfamily^35^. In addition, all 19 members have conserved XylE domain. XylE, d-xylose–H^+^ transporter of *E. coli* has been established as a homolog of GLUT transporters in *homo sapiens*^36^. Several authors have speculated that uniform mechanism of conformational change forms the basis of translocation in all MFS members, irrespective of their substrate and mode of function: **(i)** for substrate acquisition, face opens on either side of the membrane, **(ii)** occluded, distinguished as hydrophilic substrate bound in protein pocket, **(iii)** substrate release conformation i.e. opening to opposite side and **(iv)** unloaded, re-establishing its original conformation to recycle the process again. Recognized as ‘alternating access’ model of membrane transport^37^, which has been refined by biochemical characteristics of transporter^38–39^. All MFS proteins are symmetrical usually comprising 12 transmembrane helices (TMHs) with two duplicated subdomains separated by cytoplasmic loop (N-terminal TMH1–6 and C-terminal TMH7–12). In addition, the six consecutive transmembrane helices are folded into a pair of ‘3+3’ inverted repeats^40,41^. This supports our transmembrane prediction of 19 sugar porters genes **(Fig. 2)**, out of which, 9 exhibited similar domain conformation and symmetry **(Fig. S1)**. We searched the presence of GR-[KR] motifs, conserved in MFS transporters as well as the PETKG sequence, which represents the signature tag of sugar transporters^42–44^ **(Fig. S2)**. Complementing these observations, transmembrane domain analysis of PiST1 and PiST2, reflects similar TM and N- and C- terminal is cytosolic facing. PiST3 gene seems to be partial. PiST3 clusters with RGT2, surface glucose sensor of *S. cerevisiae*.

In this study, we have analyzed the expression of all 9 putative sugar porters that qualifies to possess the characteristics of MFS, during symbiotic association with maize plant. This enlightens that sugar porter which gets induced during symbiosis and critical for fungal survival. PiHXT5^28^, along with multiple sugar porters PiST1, PiST12, PiST5 and PiST6 showed significant induction during colonization **(Fig. 3)**. Since *P. indica* can be axenically cultured on synthetic media, so we successfully differentiated the expression pattern of sugar porters of *P. indica* in presence and absence of host plant symbiotic signals, not possible in AMF^18^. Therefore, highlighting the effect of mutualism on sugar porter regulation. Our findings support the report on MST2 from *G. intraradices which* have higher transcript abundance during symbiotic stage^10^. Phosphate starvation induced gene regulation is established in model organism *S. cerevisiae* to solubilize P by secreting phosphatase Pho5 in surrounding and P uptake via high affinity phosphate transporter Pho84. Such regulation involves SPX domain containing Pho81 protein that sense the P concentration in environment and regulates the localization of Pho4 through Pho80-85 mediated phosphorylation. Pho4 binds to the consensus sequence on Promoter of P responsive genes. Till date, Pi dependent regulation of hexose transporter in mutualistic partner is not known. Expression analysis revealed the critical hexose transporters of *P. indica* that are regulated via phosphate signaling and colonization *i.e.*PiST6, PiST9, PiST12 and PiST17 **(Fig. 4)**. As reported earlier by **Yadav *et al.*, 2010**^14^ the performance of *P. indica* in promoting maize growth by 1.2 and 1.8 fold in *P. indica* colonized against non-colonized plants under high and low P conditions, respectively. Since, *P. indica* is approximately 1.7 times more efficient in promoting host plant growth under low P conditions. Though this cannot be only attributed to higher expression of *PiPT* in low P condition, as it requires structural support to scavenge nutrients from deep soil that requires prominent structural growth in terms of hyphae. We have observed higher number of fungal hyphae under low P conditions as compared to high P. This further requires higher carbohydrate supply from host to fungal partner for development of hyphal structures. Therefore, we have investigated the expression of several putative hexose transporters across the *P. indica* genome under low P and high P conditions in *P. indica* colonized maize plant, and able to detect > 40 fold higher expression of a hexose transporter PiST12 under low P as compared to high P conditions. We also observed 4 fold higher expressions in axenically grown *P. indica* grown under P-limited conditions **(Fig. 4)**. PiST12 is also induced in response to glucose response **(Fig. 5)**. Our expression data of *PiST12* in response to glucose, phosphate and symbiotic stage strongly suggest that *PiST12* is the key response of *P. indica* to phosphate starvation that creates strong sink strength at plant-fungal interface to uptake carbohydrate and promotes its metabolic and structural growth to scavenge phosphate from larger volume of soil, to promote host plant growth.

Presence of conserved GR-[KR] motifs, characteristic signature sequence of MFS transporters and PETKG sequence of sugar transporters are found in PiST12, with similar substitution **(Fig. 6)**. Phylogenetic analysis revealed PiST12 cluster with fungal hexose transporters **(Fig. 7)**.

Functionally characterized hexose transporter *PiHXT5* (alias PiST17)^28^, which gets regulated in response to mutualism and glucose starvation, groups together with 9 other *P. indica* sugar transporters (PiST18, PiST3,PiST11, PiST1, PiST13, PiST17, including *PiST12*) in cluster II. *PiST17* is characterized as broad substrate monosaccharide transporters ranging from glucose, fructose, xylose, galactose, but failed to complement growth on Δ*hxt* null yeast mutant on glucose^28^. Our study revealed the regulation of *PiST12*, capable of complementing growth in heterogonous expression in *S. cerevisiae. PiST12*, which gets regulated in response to changing P concentration, showed complementation on fructose and galactose **(Fig. 8)** but negligible micro colonies were observed on glucose. High sequence similarity among different members of sugar porter in *P. indica* reflects common evolution and transport mechanism. Both *PiST12* and *PiST17* are induced under carbon starvation and are efficient hexose importers. Although, *S. cerevisiae* hexose transporters and all hexose transporters from EM fungi reported till date has highest affinity for glucose transport, BcFRT1 from *Botrytis cinerea*^45^ is characterized as fructose importer. PiST12, when heterologously expressed in yeast, complemented growth on fructose as sole carbon source.

## Materials and Methods

### Fungal and plant materials

*P. indica* strain was donated by Prof. Ajit Varma, Amity University, Noida, U.P. and maize (*Zea mays*) seeds (HQPM-5 variety) were obtained from maize research institute, Indian Agriculture Research Institute, New Delhi.

*P. indica* was routinely cultured on solidified Aspergillus modified medium (AMM)^46^. Bacterial strain DH-5α were used for cloning pupose that is cultured on LB agar plates containing adequate antibiotic. To study the expression analysis, *P. indica* culture were grown in 250 ml culture flasks with continuous shaking at 110 rpm, and at 30±2°C for 7-9 days in a metabolic shaker (Multitron Incubator Shaker, HT-Infors, Switzerland). Yeast strain was maintained on modified YPD rich medium containing 2% bacterial peptone, 0.5% ammonium phosphate, 1% yeast extract and 2% maltose (w/v). Additionally, strains were also maintained on minimal medium, 0.17% (w/v) yeast nitrogen base without amino acids required as auxotrophic supplements, and either 2% glucose, 2% fructose or 2% galactose (pH 5.5).

### Identification and Phylogenetic Analysis of Sugar Porters Genes in *P. indica*

In our previous report, we have identified 19 putative sugar porters in *P. indica*, by genome-wide screening using BLAST tool available at NCBI (www.ncbi.nlm.nih.gov), out of which only *PiHXT5, PiHXT8* and *PiHXT9* have been documented^28^. Initially, sugar porter proteins were retrieved from the *P. indica* (strain DSM 11827: taxid 1109443) genome^29^, as predicted by the pedant genome database (available at pedant.gsf.de/pedant3htmlview/pedant3view?Method=analysis&Db=p3_t65672_Pir_indic_v2). Further, BLASTx analysis was also performed^28^ to identify missing sequences from GenBank databases using functionally characterized fungal sugar porters (TbHXT1, *Tuber borchii*^26^; *S. cerevisiae* transporters^47^; AmMST1, *Amanita muscaria*^24^; BcFRT1, *Botrytis cinerea*^30^; 15 sugar porters from *Laccaria bicolor^27^;* GpMST1, *Geophison pyriformis*^9^; GiMST2, *Glomus intraradices*^10^; (**Table S1)** as template. In this study, we are reporting identified 19 putative sugar transporters, which were further verified *in silico* and analyzed with different domain prediction softwares (Expasy-Prosite, Pfam, NCBI-CDD) to fit into MFS (TIGR00879) and SP (pfam00083) family. Study includes identifying location on supercontigs, presence on forward/reverse strand, transcript and translation length, exon/intron boundaries, presence/absence of MFS, sugar porter and domains relevant to sugar transport, interval at which domain is present and genes resulted from duplication.

### Phylogenetic Tree Analysis of Sugar Porter Gene Family

Multiple sequence alignment tools like CLUSTALW2 and MULTALIN were applied to align sugar porter sequences^48^. Phylogenetic analysis were performed applying maximumlikelihood algorithm^49^ with the MEGA version 6^50^, using the JTT model of amino acid substitution^51^ according to which substitution rates of 25 discrete units were applied during heuristic search and tree was finally optimized with JTT-Gamma model. Inferred clades were supported with 1000 replicates of Bootstrapping steps^52^. Evolutionary distances were computed using Poisson correction method^53^ for the number of amino acid substitutions per site. Model of gamma distribution were used to determine the rate variation among sites (shape parameter = 1). A total of 166 protein sequences **(Table S1)** for the phylogenetic study were used and 296 positions were analyzed. Protein sequences used for phylogenetic analysis were retrieved from the genomes of *Aspergillus niger*^54^, *S. cerevisiae*^23^, *Phanerochaete chrysosporium*^55^, *Coprinopsis cinerea* (http://www.broad.mit.edu/annotation/genome/coprinus_cinereus/Home.html), *Ustilago maydis*^56^, and *Cryptococcus neoformans*^57^ putative sugar porters **(Table S1)**.

### Systemic Nomenclature of the Sugar Porter Gene Family of *P. indica*

All 19 sugar porter gene locus were traced on super contig of *P. indica* genome. The nomenclature was given in ascending order of gene locus on sequenced contigs of *P. indica* genome. The presence of Sugar Porter domain were taken into account and named as “SP”. Similar method was used for nomenclature of human GLUT proteins^58^.

### Transmembrane Helices Prediction

To identify and predict the trans-membrane (TM) domains, *PROTTER*^59^ was used to draw the TM domains of 19 potential members of sugar porter family. Further, the border of TM helices were verified by various online programs; e.g. SOSUI (http://harrier.nagahama-i-bio.ac.jp/sosui/cgibin/adv_sosui.cgi)^60^, HMMTOP server 2.0^61^, TMHMM server 2.0 (http://www.cbs.dtu.dk/services/TMHMM/)^62^, TMpred (http://www.ch.embnet.org/software/TMPRED_form.html) and TOPPRED^63^ software. The 12 TM conserved domain were observed based on MFS architecture.

### Multiple Sequence Alignment of Nine Sugar Porters

Nine Sugar Porters obtain from optimum transmembrane helices prediction, were used for multiple sequence alignment. The functional sites and their motifs in putative of 9 hexose transporter proteins were determined using PROSITE Data Bank (http://www.expasy.ch/prosite/). Homology modeling was done by using CLUSTALW2 software, (http://www.ebi.ac.uk_clustalw) and MULTALIN^48^. Alignments of multiple sequences were done with ClustalW2 with a gap penalty of 10 for insertion and 5 for extension^64,65^. *H. sapiens* (GLUT1, GLUT2, GLUT3) and *E. coli* (XylE)^40,66–67^ were also used to search conserved motifs in *P. indica* sugar porters in multiple sequence alignment. Based on carbohydrate deficiency induced expression, four hexose transporter proteins (PiST6, PiST9, PiST12 and PiST17) were selected and aligned with human GLUT1-3. The critical residues identified in GLUT1 crystal structure^40^, responsible for H-bonding in extracellular gate, sugar binding pockets and cytoplasmic exit gate were marked in alignment. Multiple sequence alignment was performed by MULTALIN and CLUSTALW2 with 80% cut off for high conservation.

### Expression analysis of Sugar Porter genes during colonization

The Primers were designed using Primer 3 **(Table S2).** Expression of Sugar porter members of *P. indica*, in colonized state with maize in comparison to its axenic state was studied. For expression analysis, *P. indica* was initially cultured in modified AMM^46^ for 8 days. Further, the culture was filtered and washed 3-4 times with MN media with high phosphate (10 mM KH2PO4) and transferred to MN media^68^ with high P and cultured for 15 days. Colonization experiments were set-up, by dipping the radicals of germinated maize seedlings into macerated liquid culture of *P. indica* and further, incubating it on water agar for 4 days. Then, it was transferred to MN liquid media with high P and grown for 15 days, whereas control plant seedlings were dipped in autoclaved double distilled water. The plantlets were transferred weekly to fresh MN media. Growth conditions provided is 30 ± 2°C, temperature; 16 h light/8 h dark, photoperiod; 60–70%, relative humidity and 1000 Lux, light intensity. After 15 days of growth, maize roots were harvested. The percentage root colonization was obtained by analyzing 10 random stained roots under microscope at 20X (Leica Microscope. Type 020-518.500, Germany). Staining was performed by softening the sample with 10% KOH for 15 min, then with 1 N HCl for 10 min, and finally staining with 0.02% trypan blue^69,70^ overnight at room temperature (or at 60°C for 1 h). De-staining of samples was performed with 50% Lacto-phenol for incubation for 1–2 hrs before proceeding to microscopic analysis^14^. The presence of distributed chlamydospores throughout the root length (analyzed per cm) was used to calculate percent colonization. Following formula was applied to obtain percent colonization^71^.

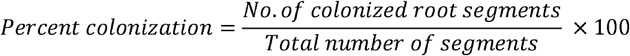

Subsequently, *P. indica* colonized maize roots were harvested and expression analysis were performed. Transcript abundance was compared by quantitative real time PCR for hexose transporters (PiST1, PiST2, PiST5, PiST6, PiST9, PiST10, PiST11, PiST12 and PiST17) in colonized and axenic *P. indica*. All experiments were performed in triplicates and significance (p-value) was calculated.

### Expression analysis of Sugar Porter genes under low P and high P conditions

To analyze the sugar porter transcript abundance in high P and low P, *P. indica* was initially cultured in modified AMM^46^ for 8 days. Further, the culture was distributed into two: for high P (10 mM KH_2_PO_4_) and low P (10 μM KH_2_PO_4_) gene induction. The culture were filtered and washed 3-4 times with MN media with high P or low P MN media^68^. Colonization experiments were set-up, by dipping the radicals of germinated maize seedlings into macerated liquid culture of *P. indica* (in high P and low P) and further, incubating it on water agar for 4 days. Then, it was transferred to MN liquid media with high P and low P and grown for 15 days, whereas control plant seedlings were dipped in autoclaved double distilled water. The plantlets were transferred weekly to fresh MN media. Growth conditions provided is 30 ± 2°C, temperature; 16 h light/8 h dark, photoperiod; 60–70%, relative humidity and 1000 Lux, light intensity. After 15 days of growth, maize roots were harvested. The percentage root colonization was studied. Subsequently, *P. indica* colonized maize roots were harvested and expression analysis were performed. Transcript abundance was compared by quantitative real time PCR for hexose transporters (PiST1, PiST2, PiST5, PiST6, PiST9, PiST10, PiST11, PiST12 and PiST17) in low P and high P conditions. All experiments were performed in triplicates and significance (p-value) was calculated.

### Expression Analysis of Sugar Porter genes under different glucose concentration

The Primers were designed using Primer 3 **(Table S2).** Expression of Sugar porter members in different glucose concentration in axenically cultivated was performed. For expression analysis, *P. indica* was initially cultured in modified AMM^46^ supplemented with 111mM (2%) glucose and high P for 8 days. Further, the culture was filtered and washed 3-4 times with MN media supplemented with 0 percent glucose^68^ with then further cultured for 2 days in different glucose concentration ranging very low to high (0, 1mM and 111mM) at 110 rpm and 30 ± 2°C. Subsequently, *P. indica* culture was harvested and expression analysis was performed. Transcript abundance was compared by quantitative real time PCR for PiST6, PiST9, PiST12 and PiST17,grown in low glucose concentration to optimum glucose concentration (111 mM glucose). All experiments were performed in triplicates and significance (p-value) was calculated.

### Isolation of RNA, cDNA Synthesis and qRT-PCR

Total RNA was isolated from *P. indica* colonized maize plant roots and from axenically grown fungus *P. indica*. Samples were harvested and washed with sterile water and frozen in liquid nitrogen. These tissue samples were then crushed in liquid nitrogen and RNA was isolated using TRIzol reagent^72^ according to the protocol provided by the manufacturer (Invitrogen, USA).

For this purpose approx. 0.2-0.5 g of transformed fungal tissue was crushed in liquid nitrogen without letting it thaw. The powdered material was transferred to a new tube; immediately 1 ml TRIZOL reagent was added and the mixture was homogenized and was incubated for 15 min at RT. Two hundred microliter of chloroform was added per ml of TRIZOL reagent and the whole mixture was vigorously vortexed for 30 sec and incubated at RT for 10 min. After centrifugation (13,000 rpm; 15 min, 4°C) the upper aqueous phase was transferred into a fresh tube and to this 0.7 volumes of isopropyl alcohol was added and the mixture was incubated for 1 h at 4°C and centrifuged (12,000 rpm; 10 min; 4°C). The supernatant was discarded and the pellet obtained was washed twice with 75% ethanol and centrifugation was done at 10,000 rpm for 5 min at 4°C. Finally, the RNA pellet obtained was dried for 10 min and dissolved in adequate volume of DEPC treated water. cDNA synthesis was performed using 2 μg total RNA. The RNA was subjected to DNase I treatment at 37°C, 30 min. Further, it was subjected to 65°C, 10 min for heat inactivation. The cDNA was synthesized using Reverse Transcriptase (250 U, Thermo Scientific) according to the manufacturer’s instruction. cDNA synthesis was performed at 42 °C for 30 min. 10 times diluted cDNA was used for gene amplification and in real time PCR reaction. For the expression of hexose transporters in different conditions, gene specific primer pairs **(Table S2)** were used. Real time were performed using SYBR Green I in ABI 7500 Real-Time PCR System (Applied Biosystems) according to manufacturer’s instructions. The reaction was set up using the program: 95°C for 10 m; 40 PCR cycles of 95°C for 15 s; 60°C for 1 m and 72°C for 25s, and the resulting fluorescence was monitored. To determine the constitutive expression of *PiTef* gene, the reaction was set up using the program; 95°C for 10 min, 40 PCR cycles of 95°C for 15 s, 57°C for 1 min, and 72 °C for 20 s. Single PCR product was confirmed by heat dissociation curve for each gene. The melting temperatures used were 81.1 °C for the amplified PCR products of *PiTef* gene, respectively. The fold change in expression of mRNA was calculated using comparative Ct method.

### Growth Promotion of maize plant and expression of *PiST12* under low P condition

The efficiency of *P. indica* in promoting growth of maize plant differed in high P and low P. As reported by **Yadav et al., 2010**^14^ that *P. indica* promoted 1.8 fold and 1.2 fold growth of maize plant in low P and high P. Since the penetration of fungal hyphae and corresponding biomass was also induced in low P conditions, so this also affects the sink strength of rhizosphere at plant-fungal interface. Further, sink strength is indicated by spatial induced expression of hexose transporters. To know the phosphate starvation dependent expression of sugar porters, we have designed this experiment. So, we have analyzed the growth promotion of Maize plant under low P condition and corresponding expression of *PiPT*^14^ and *PiST12* in order to understand the corelation between phosphate starvation, efficient growth of host maize plant promotion by *P. indica* and phosphate dependent gene regulation. For expression analysis of *PiPT* and *PiST12, P. indica* was initially cultured in modified AMM^46^ for 8 days. Further, the culture was distributed into two, in high P (10 mM KH_2_PO_4_) and low P (10 μM KH_2_PO_4_) gene induction. The culture were filtered and washed 3-4 times with MN media with high P or low P MN media^54^. Colonization experiments were set-up, by dipping the radicals of germinated maize seedlings into macerated liquid culture of *P. indica* (in high P and low P) and further, incubating it on water agar for 4 days. Further it is transferred to pots filled with sand and soil in the ratio of 3:1 (from Jawaharlal Nehru University garden and acid-washed soil). The *P. indica* colonized and non-colonized maize plants were allowed to grow for 2 weeks under low P (10 μM KH_2_PO_4_) and high P (10 mM KH_2_PO_4_) conditions. The plants were nourished weekly with half-strength modified Hoagland solution containing 10 μM and 10 mM KH_2_PO_4_ ^14^ with media composition: 2 mM MgSO_4_, 1 μM NaMoO_4_, 5 mM KNO_3_, 5 mM Ca(NO_3_)_2_, 4 μM ZnSO_4_, 50 μM H_3_BO_3_, 10 μMMgCl_2_, 1 μM CaSO_4_, and 40 mM sucrose. Growth conditions provided is 30 ± 2°C, temperature; 16 h light/8 h dark, photoperiod; 60–70%, relative humidity and 1000 Lux, light intensity. After 2 weeks of growth, maize roots were harvested. The percentage root colonization studied. Subsequently, *P. indica* colonized maize roots were harvested and expression analysis were performed. Transcript abundance was compared by quantitative real time PCR for PiPT and PiST12 in low P and high P. All experiments were performed in triplicates and significance (p-value) was calculated.

### Isolation of Coding DNA Sequence (CDS) of *PiST12*, a putative hexose transporter

*CDS* encoding *PiST12* open reading frame (ORF), was specifically amplified using gene specific primers **(Table S3)**. The PCR reaction was set up using *P. indica* cDNA (*P. indica* axenically cultured in low P 10 μM) as template. PCR mix (50 μl) contains 10 mM Tris-HCl (pH 8.3); 1.5 mM MgCl_2_; 50 mM KCl; 200 μM of dNTPs; 0.01% (w/v) gelatin; 3 U *Pfu*-polymerase 3 μM of each primer; and 80-120 ng of cDNA as template. The PCR reaction mix containing *Pfu polymerase* (Thermo Scientific) was subjected to PCR program: 94°C for 2 min; 94°C for 30 s, 58°C for 30 s, 72°C for 3 min (35 cycles); and 72°C for 5 min. PCR amplicon obtained was ligated to pGEM-T Easy vector (Promega). Competent cells *E. coli DH5-α* were prepared for CaCl_2_ mediated transformation. *E. coli DH5-α* cell was transformed with pGEMT-*PiST12* construct. Transformants were selected by blue/white screening and further subjected to colony PCR for confirmation using gene specific primers **(Table S3)**. Further, the positive clones were confirmed by restriction digestion using *EcoR*I. Lastly, the clone was confirmed by sequencing 10 different clones, to eliminate any incorporated mutation while amplification.

### Bioinformatics Analysis

Phylogenetic analysis was performed using *PiST12* putative sugar porter sequence with sugar transporters sequences from other fungi, plants and human. Phylogenetic analysis was performed applying maximum-likelihood algorithm^49^ with the MEGA version 6^50^. Inferred clades were supported with 500 replicates of Bootstrapping steps^37^. Evolutionary distances were computed using Poisson correction method^53^ for the number of amino acid substitutions per site. Model of gamma distribution were used to determine the rate variation among sites (shape parameter = 1). The sequences used for phylogenetic tree was enlisted in **Table 3.** Conserved domain analysis was performed from PFAM, NCBI on putative hexose transporter PiST12 (CCA69469.1) in P*. indica* genome. Presence of identified domains, Sugar Porter (SP) superfamily domain and Major Facilitator Superfamily were assigned in amino acid intervals **(Table 2).**Homology 3D modeling was performed using SWISS-MODEL (https://swissmodel.expasy.org/Swiss-Prot) using GLUT3 as template. Figure was obtained using Pymol.

**Table 3:** Homology of PiST12 with hexose transporters of different organism

| Sr. No. | Organism | Hexose Transporter | GenBank Accession No. | Identity with *PiST12*(%) | E-value | Query |
| --- | --- | --- | --- | --- | --- | --- |
| 1. | *Rhizoctonia solani* | Hexose transporter | EUC537746.1 | 71 | 0.0 | 99 |
| 2. | *Auricularia delicate* (fungus) | Putative hexose transporter | XP_007342912.1 | 62 | 0.0 | 93 |
| 3. | *Rhizoctonia solani* 123E (fungus) | MFS sugar transporter | KEP49025.1 | 64 | 0.0 | 89 |
| 4. | *Gloeophyllum trabeum* (fungus) | General substrate transporter | XP007862033.1 | 53 | 0.0 | 97 |
| 5. | *Trametes versicolor* (mushroom) | Putative hexose transporter | EIW61310.1 | 51 | 0.0 | 99 |
| 6. | *Coprinopsis cinerea okayama* (mushroom) | Putative hexose transporter | XP001839222.1 | 55 | 0.0 | 95 |
| 7. | *Dichomitus squalens* (fungus) | Putative hexose transporter | EJF62162.1 | 54 | 0.0 | 99 |
| 8. | *Coniophora puteana* (fungus) | Putative hexose transporter | EIW76243.1 | 49 | 0.0 | 97 |
| 9. | *Colletotrichum graminicola* (fungus) | Putative sugar transporter | ENH85362.1 | 49 | 0.0 | 89 |
| 10. | *Glomus intraradices* (AMF) | GiMST2 | ADM21463.1 | 27 | 3e-43 | 93 |
| 11. | *Geosiphon pyriformis* (AMF) | GpMST1 | CAJ77495.1 | 25 | 7e-23 | 79 |
| 12. | *Tuber borchii* (fungus) | TbHXT1 | AAY26391 | 24 | 3e-24 | 86 |
| 13. | *Saccharomyces cerevisiae* (yeast) | ScHXT1 | AAA34700.1 | 24 | 3e-29 | 80 |
| 14. | *Homo sapiens* (human) | hGLUT1 | GI636666609 | 25 | 4e-19 | 94 |
| 15. | *Zea mays* (Maize plant) | ZmMST1 | NP001105681.1 | 25 | 5e-25 | 83 |

### Yeast Complementation and Functional Analysis of Putative *PiST12*

The yeast Δ*hxt null* mutant EBY.VW4000^47^ was obtained for complementation, to characterize putative PiST12 protein as hexose transporter. In EBY.VW4000, all hexose transporters of yeast *S. cerevisiae* are deleted, unable to grow on any hexose (Glucose, Fructose, Galactose, etc.) as carbon source except maltose. For cloning the coding region of putative *PiST12gene* in yeast expression vector p112A1NE^14^, the putative *PiST12* coding region was digested out from pGEM-T easy vector by using *EcoR*I and *Hind*III and the fragment was resolved on 1% agarose/EtBr gel. The fragment was eluted from the gel using the Minelute™ gel extraction kit and subcloned in *EcoR*I and *Hind*III digested p112A1NE vector. The ligated p112A1NE-*PiST12* construct was transformed in to DH-5α *E. coli* cells. Presence of insert was confirmed by colony PCR and digestion and plasmid was isolated from 10 ml culture inoculated with single positive colony. This plasmid was used for yeast transformation.

### Yeast Transformation

Mutant Δ*hxt null* yeast strain EBY.VW4000^11^ was used for transformation. Yeast transformation was performed by lithium acetate method^73^. A single colony was inoculated into 20 ml YPD and incubated at 30°C at 220 rpm until OD_650_reaches 1-1.5. One ml of this freshly grown culture was transferred to fresh 100 ml YPD medium to produce an initial OD_600_ of 0.5. After reaching the OD_600_ of 0.5, cells were pelleted down by centrifugation at 4000 rpm for 5 min at RT. Supernatant was discarded and cell pellet was resuspended in 50 ml H_2_O. Cells were again centrifuged (same conditions as above), supernatant was discarded and pellet was resuspended in 0.5 ml sterile LiAc solution (1ml 10 X LiAc (1 M LiAc pH 7.5), 1ml 10 X TE (0.1 M Tris-HCl pH 7.5, 10 mM EDTA), and 8 ml H_2_O). 100μl of cell suspension was dispended in 1.5 ml micro centrifuge tubes, and 0.1μg of each type of plasmid was added together with 100 μg of salmon sperm DNA followed by addition of 0.6 ml sterile PEG/LiAc solution (8 ml 50 % PEG, 1ml 10 X TE, 1ml 10 X LiAc) to each tube and vortexed. Micro centrifuge tubes were incubated at 30°C for 30 min at 250 rpm. After incubation, 70 μl of 100% DMSO was added (10 % final conc.) and was mixed gently. Further, heat shock treatment was given for 15 min at 42°C, followed by chilling on ice for 5 min and later was centrifuged for 1 min at 13,000 rpm. After centrifugation supernatant was discarded and cell pellet obtained was resuspended in 0.5 mL of TE buffer. Finally 100 μl of this mixture was plated on Petri plates containing the appropriate selection medium (YNB-trp-)^23^ containing 1% maltose as sole carbon source. Plates were incubated for 3-4 days until colonies appear.

### Growth complementation analysis on different carbon sources

WT BY4741 and Δ*hxt* null EBY.VW4000 yeast *S. cerevisiae* strains were maintained in YPD medium. Putative *PiST12*-transformed yeast cells and empty p112A1NE vector-transformed Δ*hxt null* EBY.VW4000 yeast were grown in YNB-trp^-^ containing 1% maltose as sole carbon source at 30°C 220 rpm. To study the growth on different carbon source, Δ*hxt null* EBY.VW4000+ p112A1NE and Δ*hxt null* EBY.VW4000+ *p112A1NE-PiST12were* serially diluted starting with 0.1 O.D. _600_(10^-1^, 10^-2^, 10^-3^, 10^-4^) and each strain was spotted in a row on YNB 2% deficient in trp agar plates supplemented with different carbon source as selection reagents i.e. 2%Fructose, 2%Galactose, 2%Glucose and 2% Maltose. The plates were incubated at 30°C for 4–7 days to observe growth comparison between the wild type, complemented and the mutant strains.

## Reference

1. Parniske, M. Arbuscular mycorrhiza: the mother of plant root endosymbioses. Nat. Rev. Microbiol. 6, 763–775 (2008).

2. Smith, S. E., and Read, D. J. Mycorrhizal symbiosis, Academic press (2010).

3. Bonfante, P. & Genre, A. Mechanisms underlying beneficial plant–fungus interactions in mycorrhizal symbiosis. Nat. Commun. 1, 1–11 (2010).

4. Martin, F., Boiffin, V. & Pfeffer, P. E. Carbohydrate and Amino Acid Metabolism in the *Eucalyptus globulus-Pisolithus tinctorius* Ectomycorrhiza during Glucose Utilization. Plant Physiol. 118, 627–635 (1998).

5. Nehls, U., Mikolajewski, S., Magel, E. & Hampp, R. Carbohydrate metabolism in ectomycorrhizas: gene expression, monosaccharide transport and metabolic control. New Phytol. 150, 533–541 (2001).

6. Plett, J. M. & Martin, F. Blurred boundaries: lifestyle lessons from ectomycorrhizal fungal genomes. Trends Genet. 27, 14–22 (2011).

7. Bonfante, P. & Anca, I.-A. Plants, Mycorrhizal Fungi, and Bacteria: A Network of Interactions. Annu. Rev. Microbiol. 63, 363–383 (2009).

8. Smith, S. E., Dickson, S., & Smith, F. A. Nutrient transfer in arbuscular mycorrhizas: how are fungal and plant processes integrated?. Funct. Plant Biol., 28(7), 685–696 (2001).

9. Schüßler, A., Martin, H., Cohen, D., Fitz, M. & Wipf, D. Characterization of a carbohydrate transporter from symbiotic glomeromycotan fungi. Nature 444, 933–936 (2006).

10. Helber, N. et al. A Versatile Monosaccharide Transporter That Operates in the Arbuscular Mycorrhizal Fungus Glomus sp Is Crucial for the Symbiotic Relationship with Plants. The Plant Cell 23, 3812–3823 (2011).

11. Bücking, H. & Shachar-Hill, Y. Phosphate uptake, transport and transfer by the arbuscular mycorrhizal fungus *Glomus intraradices* is stimulated by increased carbohydrate availability. New Phytol 165, 899–912 (2005).

12. Fitter, A. H. What is the link between carbon and phosphorus fluxes in arbuscular mycorrhizas? A null hypothesis for symbiotic function. New Phytol. 172, 3–6 (2006).

13. Nehls, U. Mastering ectomycorrhizal symbiosis: the impact of carbohydrates. J. Exp. Bot. 59, 1097–1108 (2008).

14. Yadav, V. et al. A Phosphate Transporter from the Root Endophytic Fungus *Piriformospora indica* Plays a Role in Phosphate Transport to the Host Plant. J. Biol. Chem. 285, 26532–26544 (2010).

15. Gianinazzi-Pearson, V. Plant Cell Responses to Arbuscular Mycorrhizal Fungi: Getting to the Roots of the Symbiosis. The Plant Cell 8, 1871 (1996).

16. Schäfer, P., & Kogel, K. H. (2009). The sebacinoid fungus Piriformospora indica: an orchid mycorrhiza which may increase host plant reproduction and fitness. In Plant relationships Springer, Berlin, Heidelberg. 99–112 (2009).

17. Parniske, M. Molecular genetics of the arbuscular mycorrhizal symbiosis. Curr Opin Plant Biol.. 7, 414–421 (2004).

18. Harrison, M. J. Signaling In The Arbuscular Mycorrhizal Symbiosis. Annu. Rev. Microbiol. 59, 19–42 (2005).

19. Felle, H. H., Waller, F., Molitor, A. & Kogel, K.-H. The Mycorrhiza Fungus Piriformospora indica Induces Fast Root-Surface pH Signaling and Primes Systemic Alkalinization of the Leaf Apoplast Upon Powdery Mildew Infection. Mol. Plant-Microbe Interact. 22, 1179–1185 (2009).

20. Shachar-Hill, Y. et al. Partitioning of Intermediary Carbon Metabolism in Vesicular-Arbuscular Mycorrhizal Leek. Plant Physiol. 108, 7–15 (1995).

21. Karandashov, V. & Bucher, M. Symbiotic phosphate transport in arbuscular mycorrhizas. Trends Plant Sci. 10, 22–29 (2005).

22. Liu, C., Muchhal, U. S., Uthappa, M., Kononowicz, A. K. & Raghothama, K. G. Tomato Phosphate Transporter Genes Are Differentially Regulated in Plant Tissues by Phosphorus. Plant Physiol. 116, 91–99 (1998).

23. Boles, E. The molecular genetics of hexose transport in yeasts. FEMS Microbiol. Rev. 21, 85–111 (1997).

24. Nehls, U., Wiese, J., Guttenberger, M. & Hampp, R. Carbon Allocation in Ectomycorrhizas: Identification and Expression Analysis of an *Amanita muscaria* Monosaccharide Transporter. Mol. Plant-Microbe Interact. 11, 167–176 (1998).

25. Wiese, J., Kleber, R., Hampp, R. & Nehls, U. Functional Characterization of the *Amanita muscaria* Monosaccharide Transporter, AmMst1. Plant Biol. 2, 278–282 (2000).

26. Polidori, E. et al. Hexose uptake in the plant symbiotic ascomycete *Tuber borchii Vittadini:* biochemical features and expression pattern of the transporter TBHXT1. Fungal Genet. Biol. 44, 187–198 (2007).

27. López, M. F. et al. The sugar porter gene family of Laccaria bicolor: function in ectomycorrhizal symbiosis and soil-growing hyphae. New Phytol. 180, 365–378 (2008).

28. Rani, M. et al. Functional Characterization of a Hexose Transporter from Root Endophyte *Piriformospora indica*. Front Microbiol. 7, (2016).

29. Zuccaro, A. et al. Endophytic Life Strategies Decoded by Genome and Transcriptome Analyses of the Mutualistic Root Symbiont *Piriformospora indica*. PLoS Pathog. 7, (2011).

30. Ho, I., & Trappe, J. Translocation of 14C from Festuca plants to their endomycorrhizal fungi. Nature 244, 30–31 (1973).

31. Hodge, A., Helgason, T. & Fitter, A. Nutritional ecology of arbuscular mycorrhizal fungi. Fungal Ecol. 3, 267–273 (2010).

32. Lalonde, S., Wipf, D., & Frommer, W. B. Transport mechanisms for organic forms of carbon and nitrogen between source and sink. Annu. Rev. Plant Biol. 55, 341–372 (2004).

33. Brown, C. J., Todd, K. M., & Rosenzweig, R. F. Multiple duplications of yeast hexose transport genes in response to selection in a glucose-limited environment. Mol. biol. evol., 15(8), 931–942. (1998).

34. Wu, X., & Freeze, H. H. GLUT14, a duplicon of GLUT3, is specifically expressed in testis as alternative splice forms. Genomics, 80(6), 553–557 (2002).

35. Reddy, V. S., Shlykov, M. A., Castillo, R., Sun, E. I. & Saier, M. H. The major facilitator superfamily (MFS) revisited. FEBS J. 279, 2022–2035 (2012).

36. Maiden, M. C. J., Davis, E. O., Baldwin, S. A., Moore, D. C. M. & Henderson, P. J. F. Mammalian and bacterial sugar transport proteins are homologous. Nature 325, 641–643 (1987).

37. Jardetzky, O. Simple Allosteric Model for Membrane Pumps. Nature 211, 969–970 (1966).

38. Yan, N. Structural advances for the major facilitator superfamily (MFS) transporters. Trends in Biochem. Sci. 38, 151–159 (2013).

39. Radestock, S. & Forrest, L. R. The Alternating-Access Mechanism of MFS Transporters Arises from Inverted-Topology Repeats. J. Mol. Biol. 407, 698–715 (2011).

40. Shi, Y. Common Folds and Transport Mechanisms of Secondary Active Transporters. Ann. Rev. Biophy. 42, 51–72 (2013).

41. Henderson, P. J. F. & Baldwin, S. A. This is about the in and the out. Nat. Strut. Mol. Biol. 20, 654–655 (2013).

42. Deng, D. et al. Crystal structure of the human glucose transporter GLUT1. (2014). doi:10.2210/pdb4pyp/pdb

43. Pao, S. S., Paulsen, I. T., & Saier, M. H. Major facilitator superfamily. Microbiol. Mol. Biol. Rev. 62, 1–34 (1998).

44. Sun, C. et al. *Piriformospora indica* confers drought tolerance in Chinese cabbage leaves by stimulating antioxidant enzymes, the expression of drought-related genes and the plastid-localized CAS protein. J. Plant Physiol. 167, 1009–1017 (2010).

45. Doehlemann, G., Molitor, F. & Hahn, M. Molecular and functional characterization of a fructose specific transporter from the gray mold fungus Botrytis cinerea. Fungal Genet. Biol. 42, 601–610 (2005).

46. Hill, T. W. & Kafer, E. Improved protocols for Aspergillus minimal medium: trace element and minimal medium salt stock solutions. Fungal Genet. Rep. 48, 20–21 (2001).

47. Wieczorke, R. et al. Concurrent knock-out of at least 20 transporter genes is required to block uptake of hexoses in *Saccharomyces cerevisiae*. FEBS Lett. 464, 123–128 (1999).

48. Corpet, F. Multiple sequence alignment with hierarchical clustering. Nucleic Acids Res. 16, 10881–10890 (1988).

49. Felsenstein, J. Evolutionary trees from DNA sequences: A maximum likelihood approach. J. Mol. Evol. 17, 368–376 (1981).

50. Tamura, K., Stecher, G., Peterson, D., Filipski, A. & Kumar, S. MEGA6: Molecular Evolutionary Genetics Analysis Version 6.0. Mol. Biol. Evol. 30, 2725–2729 (2013).

51. Jones, D. T., Taylor, W. R. & Thornton, J. M. The rapid generation of mutation data matrices from protein sequences. Bioinformatics 8, 275–282 (1992).

52. Felsenstein, J. Confidence Limits on Phylogenies: An Approach Using the Bootstrap. Evolution 39, 783 (1985).

53. Zuckerkandl, E. & Pauling, L. Evolutionary Divergence and Convergence in Proteins. Evolv. Genes and Protein. 97–166 (1965).

54. Pel, H.J. et al., Genome sequencing and analysis of the versatile cell factory *Aspergillus niger CBS 513.88*. Nature biotech., 25(2), 22 (2007).

55. Martinez, D. et al. Erratum: Genome sequence of the lignocellulose degrading fungus *Phanerochaete chrysosporium* strain RP78. Nature Biotech. 22, 899–899 (2004).

56. Kämper, J. et al. Insights from the genome of the biotrophic fungal plant pathogen *Ustilago maydis*. Nature, 444(7115), 97 (2006).

57. Loftus, B.J. et al. The genome of the basidiomycetous yeast and human pathogen *Cryptococcus neoformans*. Science (2005).

58. Joost, H.-G. et al. Nomenclature of the GLUT/SLC2A family of sugar/polyol transport facilitators. Amer. J. Physiol-Endocrinol. Met. 282, (2002).

59. Omasits, U., Ahrens, C. H., Müller, S. & Wollscheid, B. Protter: interactive protein feature visualization and integration with experimental proteomic data. Bioinformatics 30, 884–886 (2013).

60. Hirokawa, T., Boon-Chieng, S. & Mitaku, S. SOSUI: classification and secondary structure prediction system for membrane proteins. Bioinformatics 14, 378–379 (1998).

61. Tusnady, G. E. & Simon, I. The HMMTOP transmembrane topology prediction server. Bioinformatics 17, 849–850 (2001).

62. Kahsay, R. Y., Gao, G. & Liao, L. An improved hidden Markov model for transmembrane protein detection and topology prediction and its applications to complete genomes. Bioinformatics 21, 1853–1858 (2005).

63. Claros, M. G. & Heijne, G. V. TopPred II: an improved software for membrane protein structure predictions. Bioinformatics 10, 685–686 (1994).

64. Clustal W (improving the sensitivity of progressive multiple sequence alignment through sequence weighting, position-specific gap penalties and weight matrix choice). Encyclo. Gen., Geno., Proteomics and Inform. 376–377 (2008). doi: 10.1007/978-1-4020-6754-9_3188

65. Henikoff, S. & Henikoff, J. G. Amino acid substitution matrices from protein blocks. Proc. Natl. Acad. Sci. USA 89, 10915–10919 (1992).

66. Iancu, C. V., Zamoon, J., Woo, S. B., Aleshin, A. & Choe, J.-Y. Crystal structure of a glucose/H symporter and its mechanism of action. Proc. Natl. Acad. Sci. USA 110, 17862–17867 (2013).

67. Wisedchaisri, G., Park, M.-S., Iadanza, M. G., Zheng, H. & Gonen, T. Proton-coupled sugar transport in the prototypical major facilitator superfamily protein XylE. Nature Commun. 5, (2014).

68. Becard, G. & Fortin, J. A. Early events of vesicular-arbuscular mycorrhiza formation on Ri T-DNA transformed roots. New Phytol. 108, 211–218 (1988).

69. Phillips, J. & Hayman, D. Improved procedures for clearing roots and staining parasitic and vesicular-arbuscular mycorrhizal fungi for rapid assessment of infection. Trans. Br. Mycol. Soc. 55, (1970).

70. Kumar, M., Yadav, V., Tuteja, N. & Johri, A. K. Antioxidant enzyme activities in maize plants colonized with *Piriformospora indica*. Microbiol. 155, 780–790 (2009).

71. Mcgonigle, T. P., Miller, M. H., Evans, D. G., Fairchild, G. L. & Swan, J. A. A new method which gives an objective measure of colonization of roots by vesicular-arbuscular mycorrhizal fungi. New Phytol. 115, 495–501 (1990).

72. Valach, M. RNA extraction using the home-made Trizol substitute v1. protocols.io (2016). doi:10.17504/protocols.io.eiebcbe

73. Gietz, R. D. & Schiestl, R. H. Applications of high efficiency lithium acetate transformation of intact yeast cells using single-stranded nucleic acids as carrier. Yeast 7, 253–263 (1991).

