## Supplementary Data for "The Sugar Porter gene family of *Piriformospora indica*: Nomenclature, Transcript Profiling and Characterization"

#### Supplementary Figures

**Figure S1. Multiple sequence alignment of *Piriformospora indica* hexose transporters *PiHXT1-9*.**Invariant and significantly conserved protein sequences are in blue and green, respectively. The conserved sugar porter family signature motifs are underscored with red lines.

**Figure S2. Multiple sequence alignment of hexose transporter of *P. indica* with hGLUT1-3.**Invariant and highly conserved amino acids are red and blue, respectively. The critical residues for H-bonding at extracellular gate glucose transport and cytoplasmic gate as reported in hGLUT1 crystal structure (Deng *et al.*, 2014) were shown in yellow box.

**Figure S3. Conserved domain on putative hexose transporter PiHXT3** (CCA69469.1) in *P. indica* genome: Complete Sugar Porter (SP) Superfamily domain and Major Facilitator Superfamily (MFS) domain is present. Moreover, sugar transporter PFAM motif (PF0083) was also present in PiHXT3 polypeptide at position 44–485 and MFS Superfamily domain (cd6174) is present at position 47–470 **(Table S4)**.

**Figure S4. Putative *PiST12* ORF sequence :** A. PiST12 ORF sequence (1593 bp) showing start and stop codon.

**Figure S5. Deduced putative PiHXT3 protein:** Deduced amino acid sequence (530 AA) of putative *PiHXT3* protein. The prediction was done by BioEdit tool (Hall *et al.,* 1999).

**Figure S6. Schematic Genomic orientation, ORF and 3 D model of PiST12 protein: (i)** It was observed that putative *PiST12* has 1769 bp at genomic sequence, 1593 bp ORF sequence and 530 amino acids long protein. *PiST12* has 4 exons (denoted by E1-E4) and 3 introns (denoted by I1-I3). **(ii)3D Model of PiST12 Putative hexose transporter.** 3D modeling was performed using Swiss Prot Template based modeling using GLUT 3 as template. Further image adjustment were performed using Pymol (A) Side view (B) view after rotating 180ºC (C) Extracellular view, (D) cytoplasmic view.

**Figure S7. Phylogeny of protein sequence of PiHXT3**. Fungal monosaccharide transporter protein sequences with GenBank Accession Number CCA69469.1. The phylogenetic tree was constructed using Mega 6.06 software applying the Neighbor-joining method for the construction of the phylogeny. Bootstrap test were performed using 500 replicates. The branch lengths are proportional to the phylogenetic distance. *P. indica PiST12* cluster together with fungal hexose transporters from basidiomycota fungi.

**Figure S8.Analysis of development of three weeks old maize plants colonized with WT *P. indica*, under Low P (10µM).** (A) Morphology of maize plants (i) non-colonized (i) colonized with WT *P. indica* grown under low P (10µM) at 15 days.(B) Growth comparison of the root length of the maize plants (i) non-colonized (i) colonized with WT *P. indica* grown under low P (10µM) at 15 days (C) *P. indica* promotes more growth percentage in low P as compared to high P. *Data is statistically significant ($P<0.005$) as compared to WT and over-expressed P. *indica* colonizing plant under P-limited condition. Each column represents the mean of five observations ±Standard Error. (D) Quantitative and Semi quantitative expression analysis of *PiST12* and *PiPT* in WT *P. indica* inoculated under maize plant, grown under (i) low P (10µM) and (ii) high P (10mM) at 12 dpi.

**Figure S9. Phylogenetic relationships of the deduced protein sequences of the *Piriformospora indica* sugar porter gene family with known fungal hexose transporters.**(Magnified view) Putative hexose transporters deduced from the genome sequence of range of fungus, plants and animal kingdom including *homo sapiens* which is enlisted in **Table S1**. Phylogenetic tree was constructed using MEGA6. The Phylogenetic relationships were obtained by maximum-likelihood method, using the JTT model of amino acid substitution. Number above branches indicates bootstrap values from 1000 replicates. Sugar porter gene family of *P. indica* was grouped into three clusters. The tree was unrooted **(Figure S9 i-iii)**. N-glycosylation sites were also shown.

**Figure S10. Prediction of Transmembrane domain of Sugar Porter family of *P. indica* (Magnified view):** To predict the transmembrane domain and N- and C- face of sugar porter proteins, TM-Pred, HMM-TOP, SOSUI and PROTTER were applied. All 19 PiST were analyzed by this method and their topologies were obtained. **(i)** PiST1 **(ii)** PiST2 **(iii)** PiST3 **(iv)** PiST4 **(v)** PiST5 **(vi)** PiST6 **(vii)** PiST7 **(viii)** PiST8 **(ix)** PiST9 **(x)** PiST10 **(xi)** PiST11 **(xii)** PiST12 **(xiii)** PiST13 **(xiv)** PiST14**(xv)** PiST15 **(xvi)** PiST16 **(xvii)** PiST17 **(xviii)** PiST18 **(xix)** PiST19

**Supplementary Tables**

**Table S1:** Details of Sequence used in Phylogenetic Analysis of Sugar Porter Family of *P. indica.*

**Table S2:** Primers for expression analysis of Hexose Transporters from *P. indica.*

**Table S3:** Primers used in cloning of *PiST12* and its sequencing.

**Table S4:** List of Domains present on PiST12 ORF.

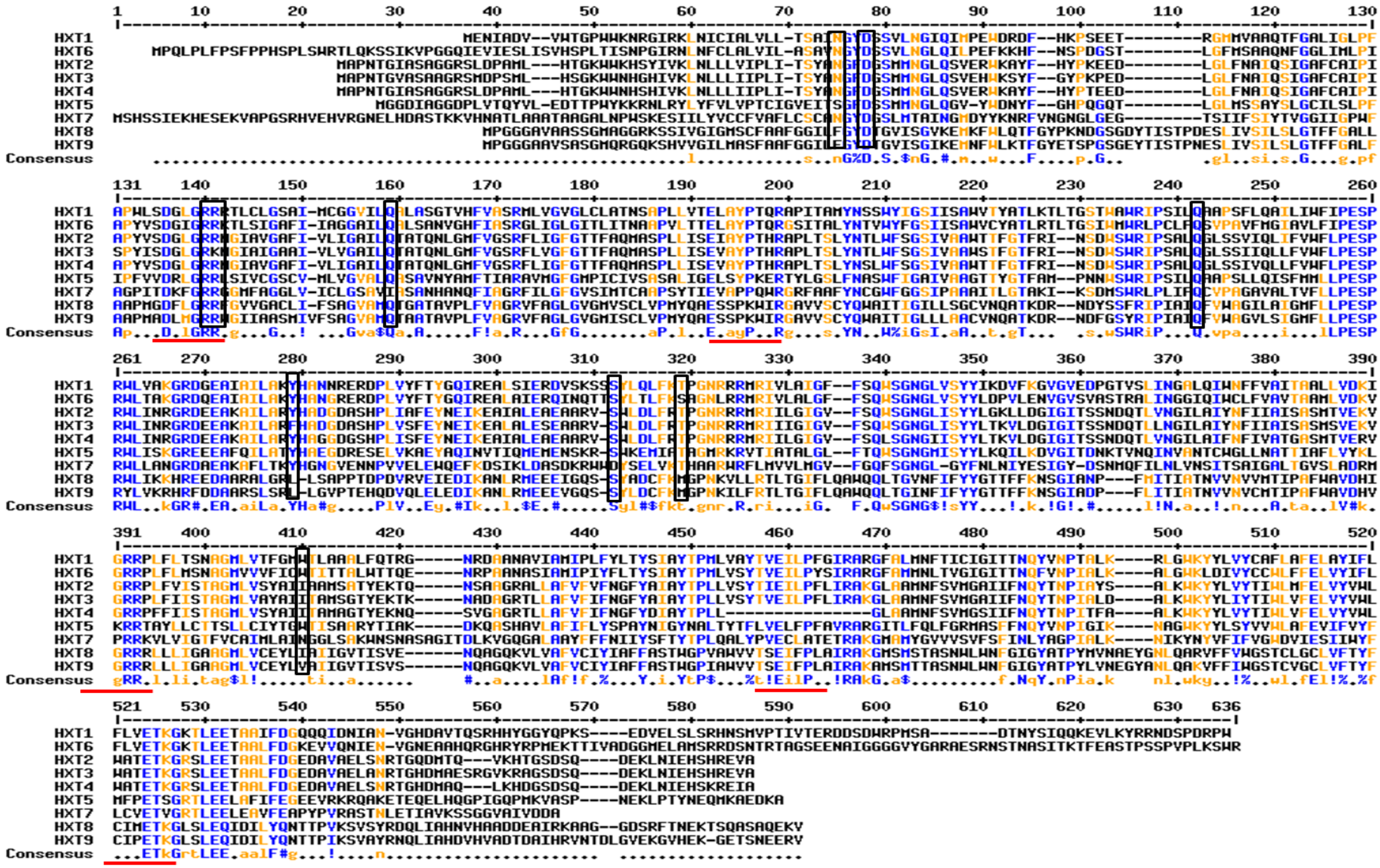

**Figure S1**

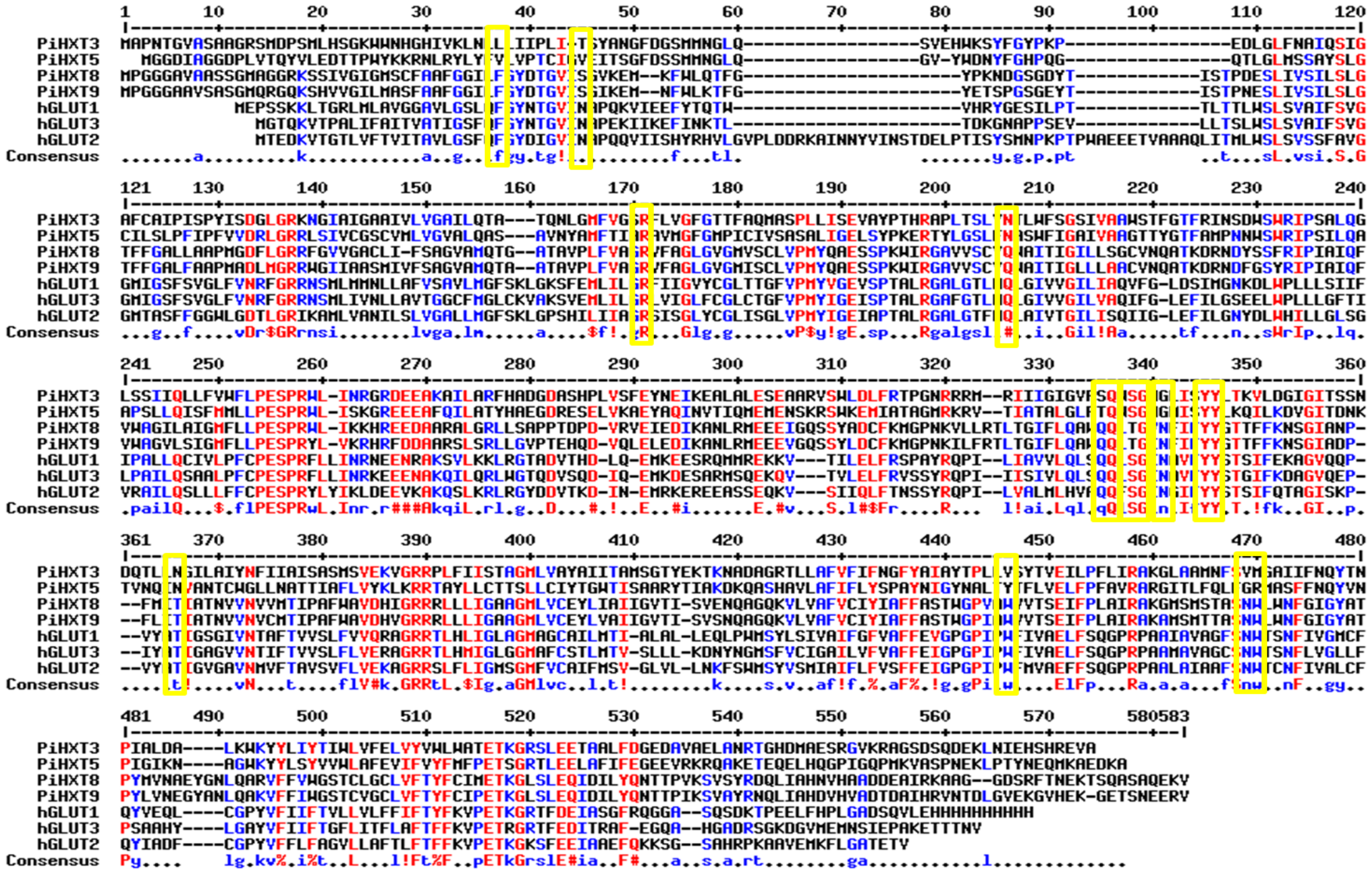
PiHXT3
PiHXT5
PiHXT8
PiHXT9
hGLUT1
hGLUT3
hGLUT2
Consensus

**Figure S2**

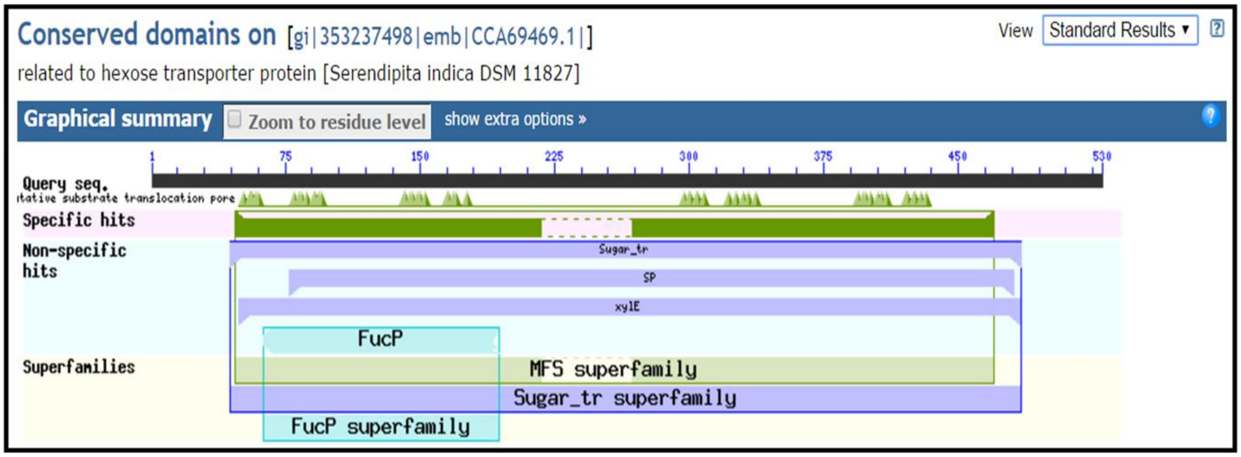
Conserved domains on [gi|353237498|emb|CCA69469.1|]
View Standard Results
related to hexose transporter protein [Serendipita indica DSM 11827]
Graphical summary
Zoom to residue level
show extra options »
1
75
150
225
300
375
450
530
Query seq.
Specific hits
Non-specific hits
Sugar_tr
SP
xylE
FucP
Superfamilies
MFS superfamily
Sugar_tr superfamily
FucP superfamily

**Figure S3**

**ATG**GCTCCCAACACTGGTGTCGCCTCGGCTGCTGGCCGTTCGATGGATCCCTCCATGCTCCACTCTGGAA
AATGGTGGAACCACGGCCATATTGTCAAGCTCAACTTGCTGCTCATCATCCCCCTCATCACCTCGTATGC
CAACGGTTTCGACGGTTCGATGATGAACGGTCTCCAGTCGGTCGAGCACTGGAAGTCTTACTTTGGCTAT
CCAAAGCCCGAGGATCTCGGTCTCTTCAACGCCATCCAGTCCATTGGTGCCTTCTGCGCCATTCCCATCT
CGCCATACATCTCTGATGGTCTTGGTCGCAAGAACGGTATTGCCATTGGTGCCGCCATTGTCCTCGTCGG
TGCTATCCTCCAAACGGCTACCCAGAACCTTGGCATGTTCGTCGGTTCCCGTTTCCTCGTCGGTTTCGGA
ACCACCTTTGCCCAGATGGCTTCTCCTCTTCTCATCTCCGAGGTTGCCTACCCAACTCACCGTGCTCCCC
TCACTTCGCTCTACAACACCCTCTGGTTCTCTGGCTCCATCGTTGCCGCCTGGTCGACTTTCGGTACCTT
CCGCATCAACAGTGACTGGTCCTGGCGTATTCCCTCGGCTCTCCAGGGTCTCTCGTCGATCATCCAGCTC
CTCTTCGTCTGGTTCCTTCCCGAGTCTCCTCGTTGGTTGATCAATCGCGGTCGCGATGAAGAGGCCAAGG
CCATCCTTGCCCGCTTCCACGCCGATGGTGATGCGTCGCACCCACTCGTCTCGTTTGAGTACAACGAAAT
CAAGGAGGCCCTCGCTCTCGAATCCGAGGCTGCTCGCGTCTCTTGGCTCGATCTCTTCCGCACCCCCGGT
AACCGTCGCCGCATGAGGATCATCATCGGTATCGGTGTCTTCTCCCAGTGGTCCGGAAACGGTCTCATCT
CCTACTACCTCACCAAGGTCCTCGACGGTATTGGAATCACGAGCTCTAACGACCAGACTCTTCTTAACGG
TATCCTCGCCATCTACAACTTTATCATTGCCATCTCTGCCTCCATGTCCGTCGAGAAGGTTGGACGTCGC
CCACTCTTCATCATCTCGACCGCTGGTATGCTTGTCGCCTACGCCATCATCACTGCCATGTCCGGAACGT
ACGAGAAGACCAAGAACGCTGATGCTGGACGTACCCTCCTCGCATTCGTCTTCATCTTCAACGGTTTCTA
CGCCATCGCCTATACTCCTCTCCTCGTCTCGTACACTGTCGAAATTCTTCCTTTCCTCATTCGTGCCAAG
GGTCTCGCTGCGATGAACTTCTCCGTCATGGGTGCCATCATCTTCAACCAGTACACCAACCCCATCGCCC
TCGACGCTCTCAAGTGGAAGTACTACCTCATCTACACGATCTGGCTCGTGTTCGAGTTGGTCTACGTCTG
GCTCTGGGCCACCGAGACCAAGGGTCGTTCGCTCGAGGAGACTGCAGCCCTGTTCGATGGTGAGGATGCC
GTCGCAGAGCTTGCCAACCGCACTGGCCACGATATGGCCGAGTCTCGCGGTGTCAAGCGCGCTGGCTCGG
ATTCGCAGGACGAGAAGCTCAACATTGAGCACTCTCACCGCGAGGTTGCC**TGA**

**Figure S4**

```
1     ATGGCTCCCAACACTGGTGTCGCCTCGGCTGCTGGCCGTTCGATGGATCCCTCCATGCTC 60
1      M  A  P  N  T  G  V  A  S  A  A  G  R  S  M  D  P  S  M  L  20

61    CACTCTGGAAAATGGTGGAACCACGGCCATATTGTCAAGCTCAACTTGCTGCTCATCATC 120
21     H  S  G  K  W  W  N  H  G  H  I  V  K  L  N  L  L  L  I  I  40

121   CCCCTCATCACCTCGTATGCCAACGGTTTCGACGGTTCGATGATGAACGGTCTCCAGTCG 180
41     P  L  I  T  S  Y  A  N  G  F  D  G  S  M  M  N  G  L  Q  S  60

181   GTCGAGCACTGGAAGTCTTACTTTGGCTATCCAAAGCCCGAGGATCTCGGTCTCTTCAAC 240
61     V  E  H  W  K  S  Y  F  G  Y  P  K  P  E  D  L  G  L  F  N  80

241   GCCATCCAGTCCATTGGTGCCTTCTGCGCCATTCCCATCTCGCCATACATCTCTGATGGT 300
81     A  I  Q  S  I  G  A  F  C  A  I  P  I  S  P  Y  I  S  D  G  100

301   CTTGGTCGCAAGAACGGTATTGCCATTGGTGCCGCCATTGTCCTCGTCGGTGCTATCCTC 360
101    L  G  R  K  N  G  I  A  I  G  A  A  I  V  L  V  G  A  I  L  120

361   CAAACGGCTACCCAGAACCTTGGCATGTTCGTCGGTTCCCGTTTCCTCGTCGGTTTCGGA 420
121    Q  T  A  T  Q  N  L  G  M  F  V  G  S  R  F  L  V  G  F  G  140

421   ACCACCTTTGCCCAGATGGCTTCTCCTCTTCTCATCTCCGAGGTTGCCTACCCAACTCAC 480
141    T  T  F  A  Q  M  A  S  P  L  L  I  S  E  V  A  Y  P  T  H  160

481   CGTGCTCCCCTCACTTCGCTCTACAACACCCTCTGGTTCTCTGGCTCCATCGTTGCCGCC 540
161    R  A  P  L  T  S  L  Y  N  T  L  W  F  S  G  S  I  V  A  A  180

541   TGGTCGACTTTCGGTACCTTCCGCATCAACAGTGACTGGTCCTGGCGTATTCCCTCGGCT 600
181    W  S  T  F  G  T  F  R  I  N  S  D  W  S  W  R  I  P  S  A  200

601   CTCCAGGGTCTCTCGTCGATCATCCAGCTCCTCTTCGTCTGGTTCCTTCCCGAGTCTCCT 660
201    L  Q  G  L  S  S  I  I  Q  L  L  F  V  W  F  L  P  E  S  P  220

661   CGTTGGTTGATCAATCGCGGTCGCGATGAAGAGGCCAAGGCCATCCTTGCCCGCTTCCAC 720
221    R  W  L  I  N  R  G  R  D  E  E  A  K  A  I  L  A  R  F  H  240

721   GCCGATGGTGATGCGTCGCACCCACTCGTCTCGTTTGAGTACAACGAAATCAAGGAGGCC 780
241    A  D  G  D  A  S  H  P  L  V  S  F  E  Y  N  E  I  K  E  A  260

781   CTCGCTCTCGAATCCGAGGCTGCTCGCGTCTCTTGGCTCGATCTCTTCCGCACCCCCGGT 840
261    L  A  L  E  S  E  A  A  R  V  S  W  L  D  L  F  R  T  P  G  280

841   AACCGTCGCCGCATGAGGATCATCATCGGTATCGGTGTCTTCTCCCAGTGGTCCGGAAAC 900
281    N  R  R  R  M  R  I  I  I  G  I  G  V  F  S  Q  W  S  G  N  300

901   GGTCTCATCTCCTACTACCTCACCAAGGTCCTCGACGGTATTGGAATCACGAGCTCTAAC 960
301    G  L  I  S  Y  Y  L  T  K  V  L  D  G  I  G  I  T  S  S  N  320

961   GACCAGACTCTTCTTAACGGTATCCTCGCCATCTACAACTTTATCATTGCCATCTCTGCC 1020
321    D  Q  T  L  L  N  G  I  L  A  I  Y  N  F  I  I  A  I  S  A  340

1021  TCCATGTCCGTCGAGAAGGTTGGACGTCGCCCACTCTTCATCATCTCGACCGCTGGTATG 1080
341    S  M  S  V  E  K  V  G  R  R  P  L  F  I  I  S  T  A  G  M  360

1081  CTTGTCGCCTACGCCATCATCACTGCCATGTCCGGAACGTACGAGAAGACCAAGAACGCT 1140
361    L  V  A  Y  A  I  I  T  A  M  S  G  T  Y  E  K  T  K  N  A  380
```

```
1141 GATGCTGGACGTACCCTCCTCGCATTCGTCTTCATCTTCAACGGTTTCTACGCCATCGCC 1200
381    D  A  G  R  T  L  L  A  F  V  F  I  F  N  G  F  Y  A  I  A  400

1201 TATACTCCTCTCCTCGTCTCGTACACTGTCGAAATTCTTCCTTTCCTCATTCGTGCCAAG 1260
401    Y  T  P  L  L  V  S  Y  T  V  E  I  L  P  F  L  I  R  A  K  420

1261 GGTCTCGCTGCGATGAACTTCTCCGTCATGGGTGCCATCATCTTCAACCAGTACACCAAC 1320
421    G  L  A  A  M  N  F  S  V  M  G  A  I  I  F  N  Q  Y  T  N  440

1321 CCCATCGCCCTCGACGCTCTCAAGTGGAAGTACTACCTCATCTACACGATCTGGCTCGTG 1380
441    P  I  A  L  D  A  L  K  W  K  Y  Y  L  I  Y  T  I  W  L  V  460

1381 TTCGAGTTGGTCTACGTCTGGCTCTGGGCCACCGAGACCAAGGGTCGTTCGCTCGAGGAG 1440
461    F  E  L  V  Y  V  W  L  W  A  T  E  T  K  G  R  S  L  E  E  480

1441 ACTGCAGCCCTGTTCGATGGTGAGGATGCCGTCGCAGAGCTTGCCAACCGCACTGGCCAC 1500
481    T  A  A  L  F  D  G  E  D  A  V  A  E  L  A  N  R  T  G  H  500

1501 GATATGGCCGAGTCTCGCGGTGTCAAGCGCGCTGGCTCGGATTCGCAGGACGAGAAGCTC 1560
501    D  M  A  E  S  R  G  V  K  R  A  G  S  D  S  Q  D  E  K  L  520

1561 AACATTGAGCACTCTCACCGCGAGGTTGCCTGA 1593
521    N  I  E  H  S  H  R  E  V  A  *  530
```

**Figure S5**

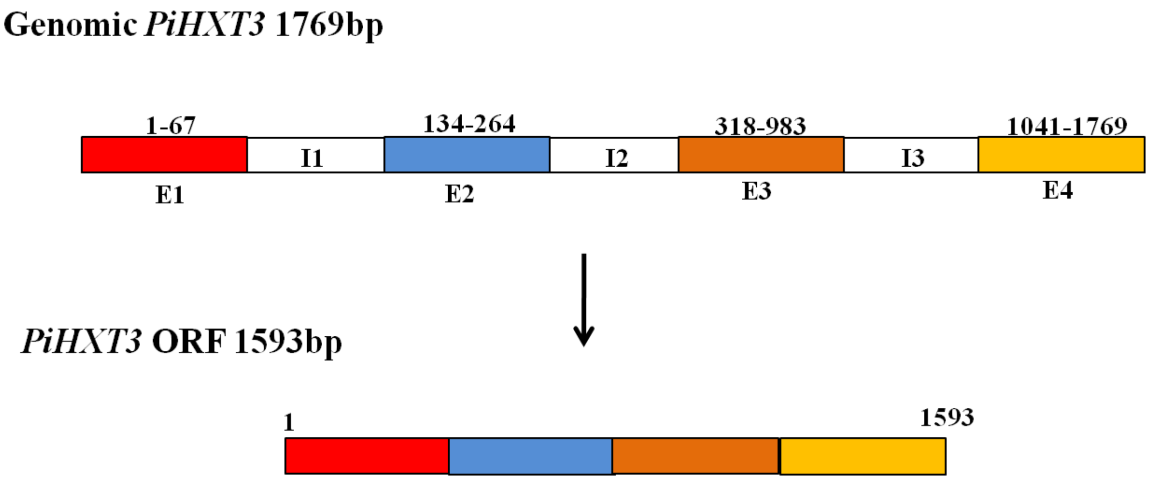


**(i)**

**Figure S6**

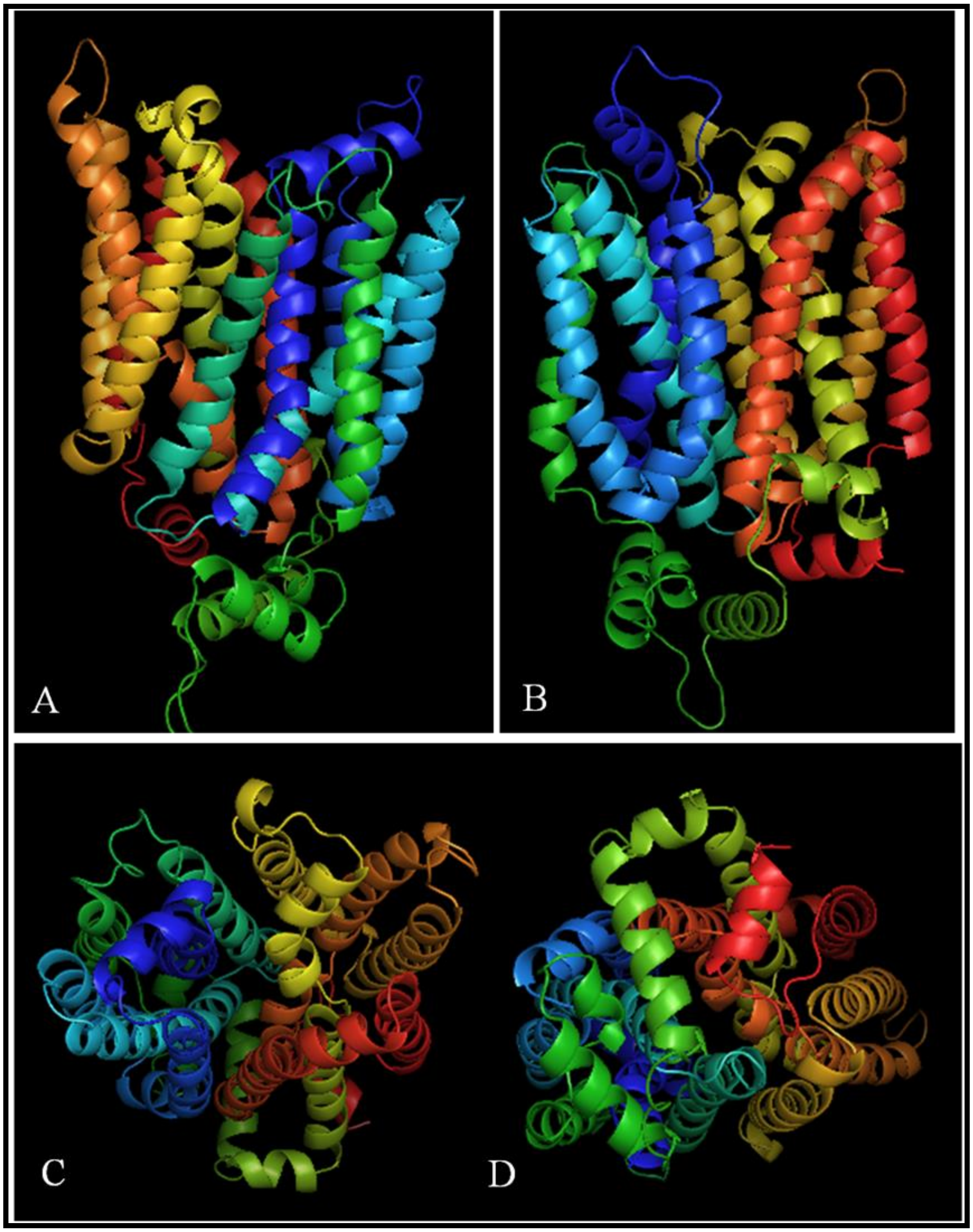


**(ii)**

**Figure S6**

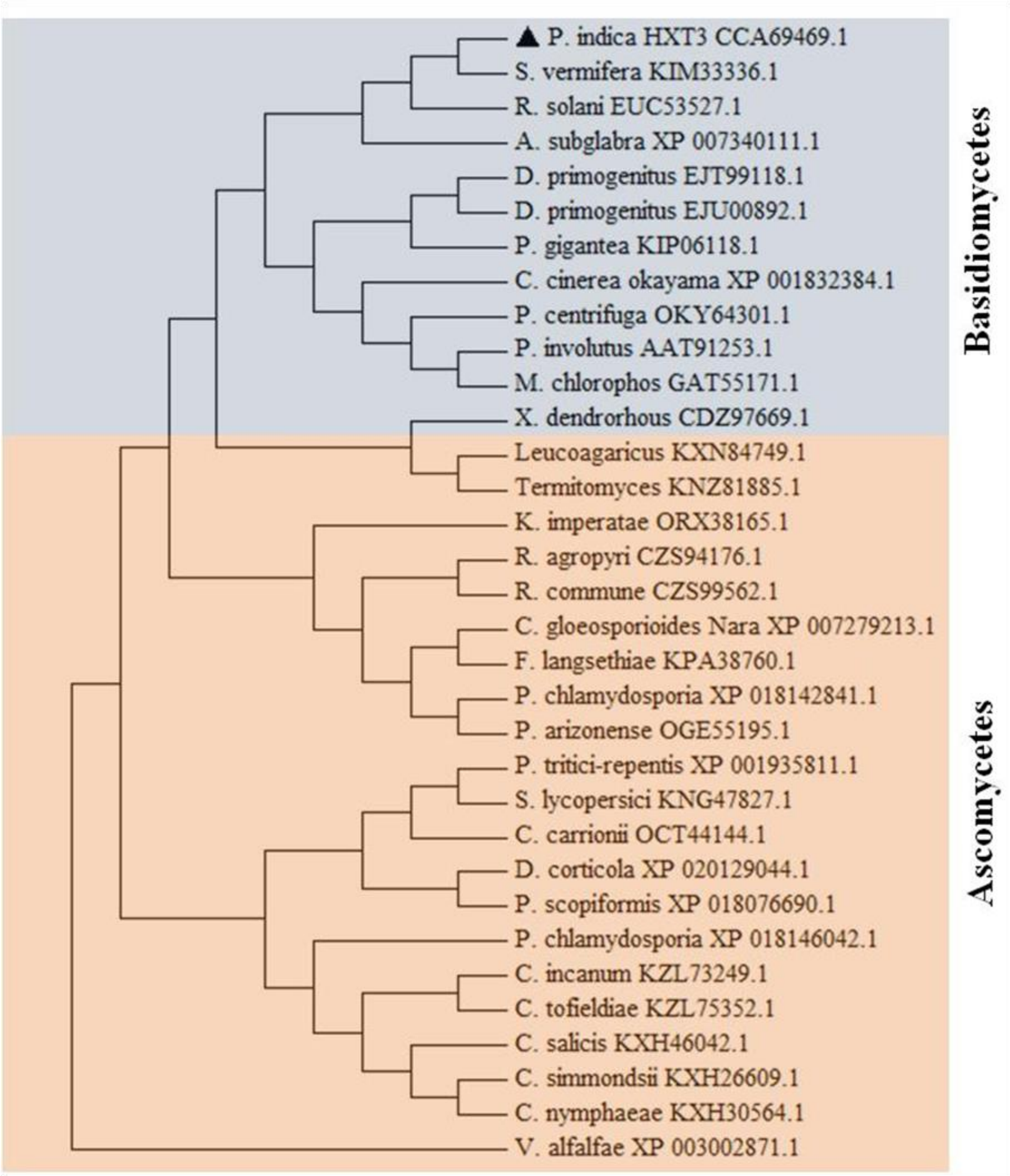
▲ P. indica HXT3 CCA69469.1
S. vermifera KIM33336.1
R. solani EUC53527.1
A. subglabra XP 007340111.1
D. primogenitus EJT99118.1
D. primogenitus EJU00892.1
P. gigantea KIP06118.1
C. cinerea okayama XP 001832384.1
P. centrifuga OKY64301.1
P. involutus AAT91253.1
M. chlorophos GAT55171.1
X. dendrorhous CDZ97669.1
Leucoagaricus KXN84749.1
Termitomyces KNZ81885.1
K. imperatae ORX38165.1
R. agropyri CZS94176.1
R. commune CZS99562.1
C. gloeosporioides Nara XP 007279213.1
F. langsethiae KPA38760.1
P. chlamydosporia XP 018142841.1
P. arizonense OGE55195.1
P. tritici-repentis XP 001935811.1
S. lycopersici KNG47827.1
C. carrionii OCT44144.1
D. corticola XP 020129044.1
P. scopiformis XP 018076690.1
P. chlamydosporia XP 018146042.1
C. incanum KZL73249.1
C. tofieldiae KZL75352.1
C. salicis KXH46042.1
C. simmondsii KXH26609.1
C. nymphaeae KXH30564.1
V. alfalfae XP 003002871.1
Basidiomycetes
Ascomycetes

**Figure S7**

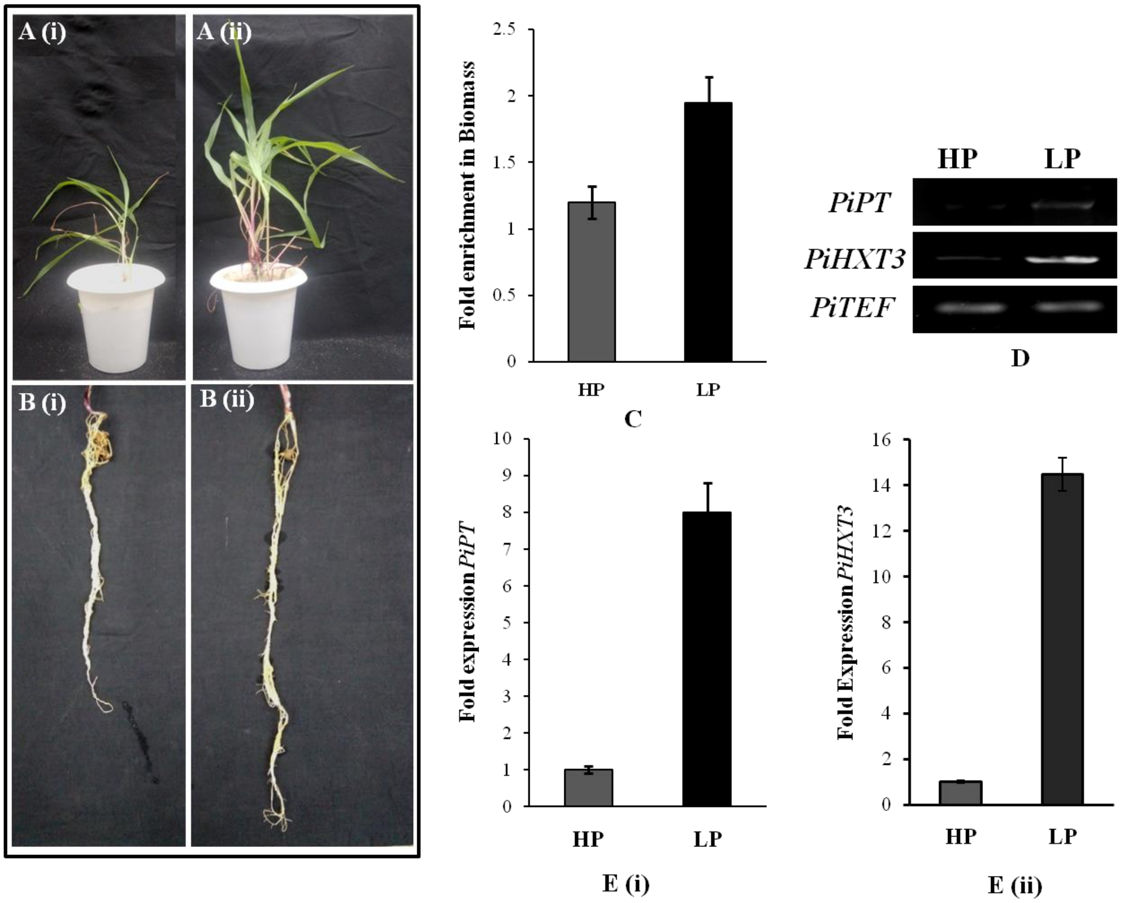
A (i)
A (ii)
B (i)
B (ii)
Fold enrichment in Biomass
2.5
2
1.5
1
0.5
0
HP
LP
C
HP
LP
PiPT
PiHXT3
PiTEF
D
Fold expression PiPT
10
9
8
7
6
5
4
3
2
1
0
HP
LP
E (i)
Fold Expression PiHXT3
16
14
12
10
8
6
4
2
0
HP
LP
E (ii)

**Figure S8**

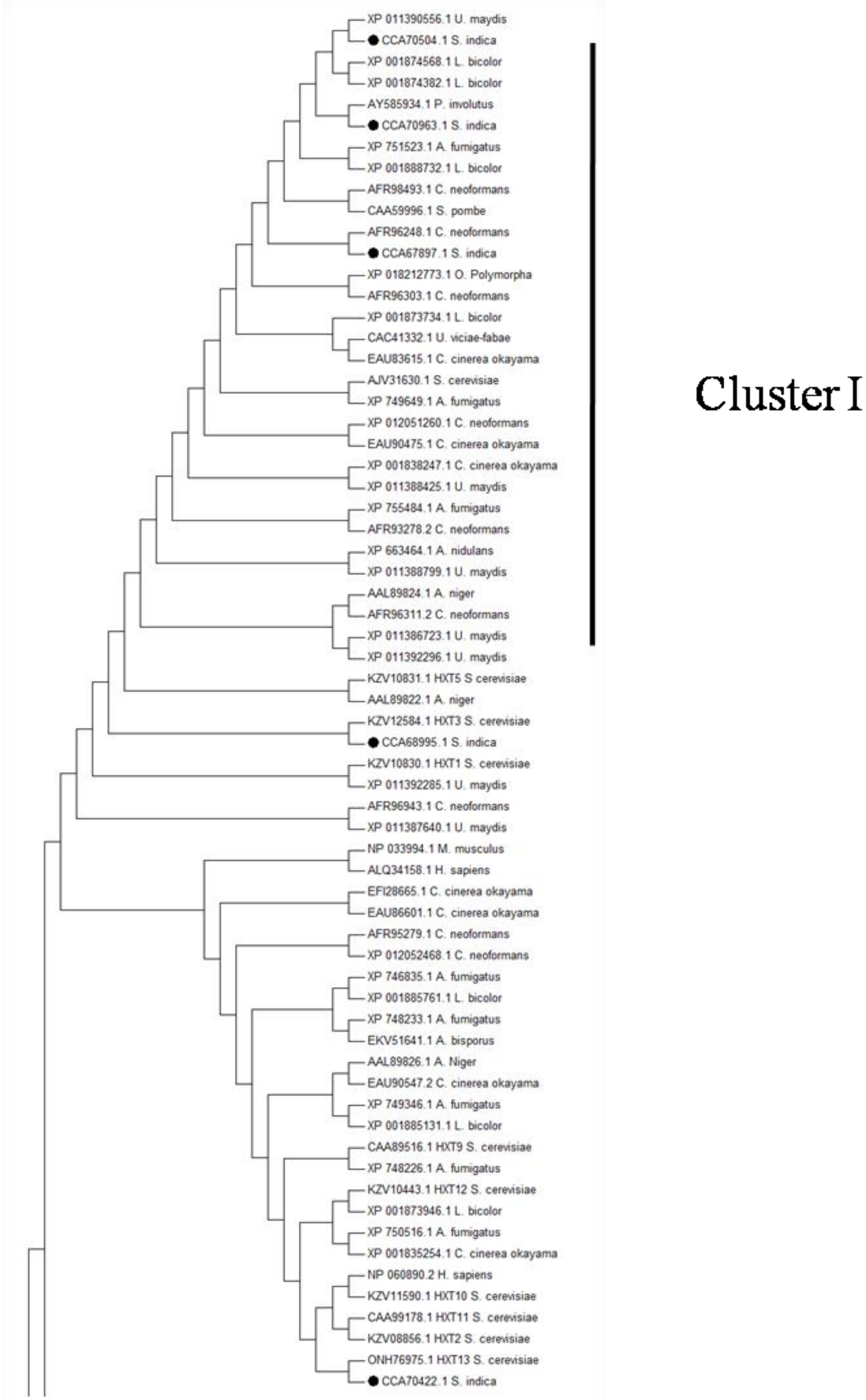
XP 011390556.1 U. maydis
CCA70504.1 S. indica
XP 001874568.1 L. bicolor
XP 001874382.1 L. bicolor
AY585934.1 P. involutus
CCA70963.1 S. indica
XP 751523.1 A. fumigatus
XP 001888732.1 L. bicolor
AFR98493.1 C. neoformans
CAA59996.1 S. pombe
AFR96248.1 C. neoformans
CCA67897.1 S. indica
XP 018212773.1 O. Polymorpha
AFR96303.1 C. neoformans
XP 001873734.1 L. bicolor
CAC41332.1 U. viciae-fabae
EAU83615.1 C. cinerea okayama
AJV31630.1 S. cerevisiae
XP 749649.1 A. fumigatus
XP 012051260.1 C. neoformans
EAU90475.1 C. cinerea okayama
XP 001838247.1 C. cinerea okayama
XP 011388425.1 U. maydis
XP 755484.1 A. fumigatus
AFR93278.2 C. neoformans
XP 663464.1 A. nidulans
XP 011388799.1 U. maydis
AAL89824.1 A. niger
AFR96311.2 C. neoformans
XP 011386723.1 U. maydis
XP 011392296.1 U. maydis
KZV10831.1 HXT5 S cerevisiae
AAL89822.1 A. niger
KZV12584.1 HXT3 S. cerevisiae
CCA68995.1 S. indica
KZV10830.1 HXT1 S. cerevisiae
XP 011392285.1 U. maydis
AFR96943.1 C. neoformans
XP 011387640.1 U. maydis
NP 033994.1 M. musculus
ALQ34158.1 H. sapiens
EFI28665.1 C. cinerea okayama
EAU86601.1 C. cinerea okayama
AFR95279.1 C. neoformans
XP 012052468.1 C. neoformans
XP 746835.1 A. fumigatus
XP 001885761.1 L. bicolor
XP 748233.1 A. fumigatus
EKV51641.1 A. bisporus
AAL89826.1 A. Niger
EAU90547.2 C. cinerea okayama
XP 749346.1 A. fumigatus
XP 001885131.1 L. bicolor
CAA89516.1 HXT9 S. cerevisiae
XP 748226.1 A. fumigatus
KZV10443.1 HXT12 S. cerevisiae
XP 001873946.1 L. bicolor
XP 750516.1 A. fumigatus
XP 001835254.1 C. cinerea okayama
NP 060890.2 H. sapiens
KZV11590.1 HXT10 S. cerevisiae
CAA99178.1 HXT11 S. cerevisiae
KZV08856.1 HXT2 S. cerevisiae
ONH76975.1 HXT13 S. cerevisiae
CCA70422.1 S. indica
Cluster I

**Figure S9 (i)**

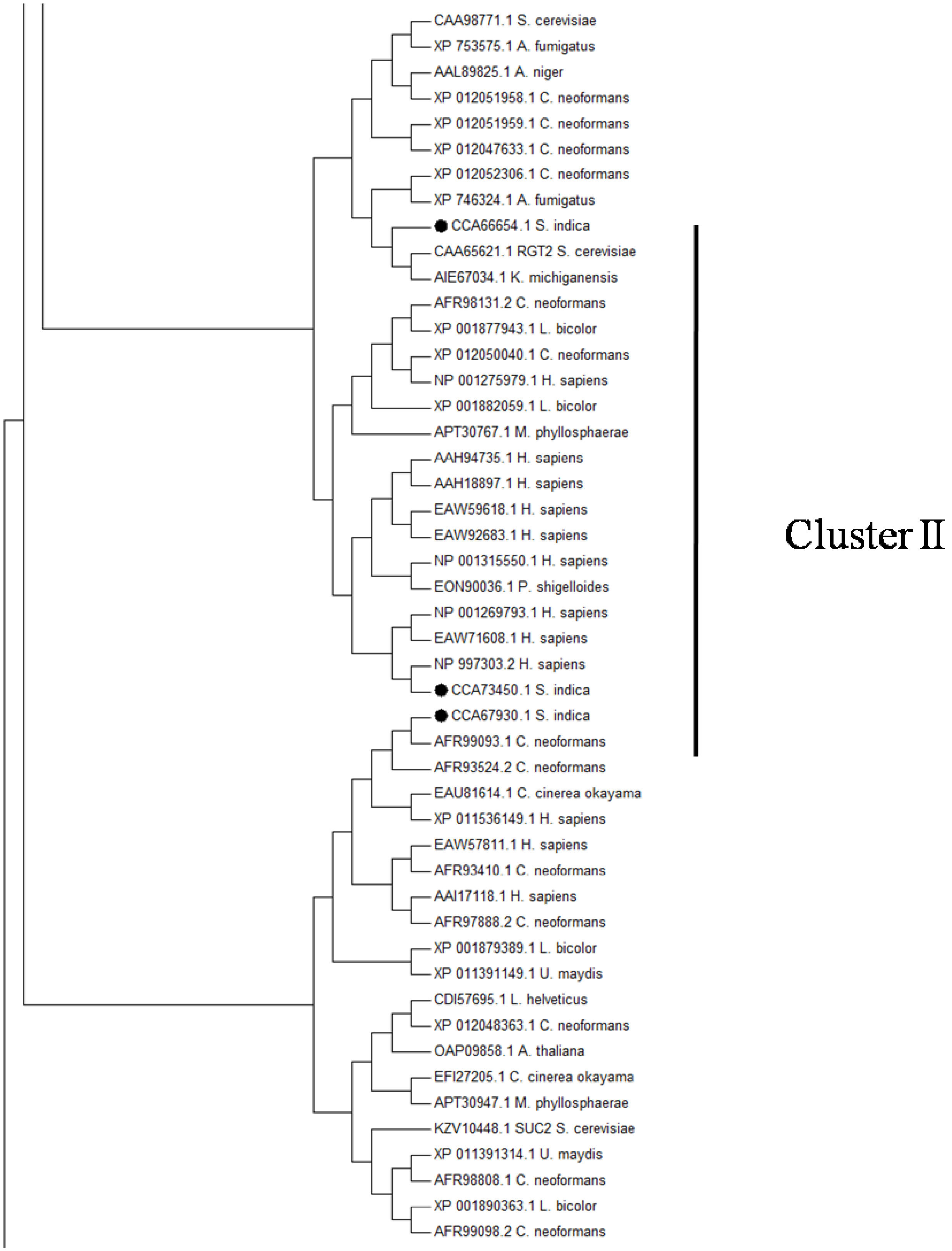
CAA98771.1 S. cerevisiae
XP 753575.1 A. fumigatus
AAL89825.1 A. niger
XP 012051958.1 C. neoformans
XP 012051959.1 C. neoformans
XP 012047633.1 C. neoformans
XP 012052306.1 C. neoformans
XP 746324.1 A. fumigatus
CCA66654.1 S. indica
CAA65621.1 RGT2 S. cerevisiae
AIE67034.1 K. michiganensis
AFR98131.2 C. neoformans
XP 001877943.1 L. bicolor
XP 012050040.1 C. neoformans
NP 001275979.1 H. sapiens
XP 001882059.1 L. bicolor
APT30767.1 M. phyllosphaerae
AAH94735.1 H. sapiens
AAH18897.1 H. sapiens
EAW59618.1 H. sapiens
EAW92683.1 H. sapiens
NP 001315550.1 H. sapiens
EON90036.1 P. shigelloides
NP 001269793.1 H. sapiens
EAW71608.1 H. sapiens
NP 997303.2 H. sapiens
CCA73450.1 S. indica
CCA67930.1 S. indica
AFR99093.1 C. neoformans
AFR93524.2 C. neoformans
EAU81614.1 C. cinerea okayama
XP 011536149.1 H. sapiens
EAW57811.1 H. sapiens
AFR93410.1 C. neoformans
AAI17118.1 H. sapiens
AFR97888.2 C. neoformans
XP 001879389.1 L. bicolor
XP 011391149.1 U. maydis
CDI57695.1 L. helveticus
XP 012048363.1 C. neoformans
OAP09858.1 A. thaliana
EFI27205.1 C. cinerea okayama
APT30947.1 M. phyllosphaerae
KZV10448.1 SUC2 S. cerevisiae
XP 011391314.1 U. maydis
AFR98808.1 C. neoformans
XP 001890363.1 L. bicolor
AFR99098.2 C. neoformans
Cluster II

**Figure S9(ii)**

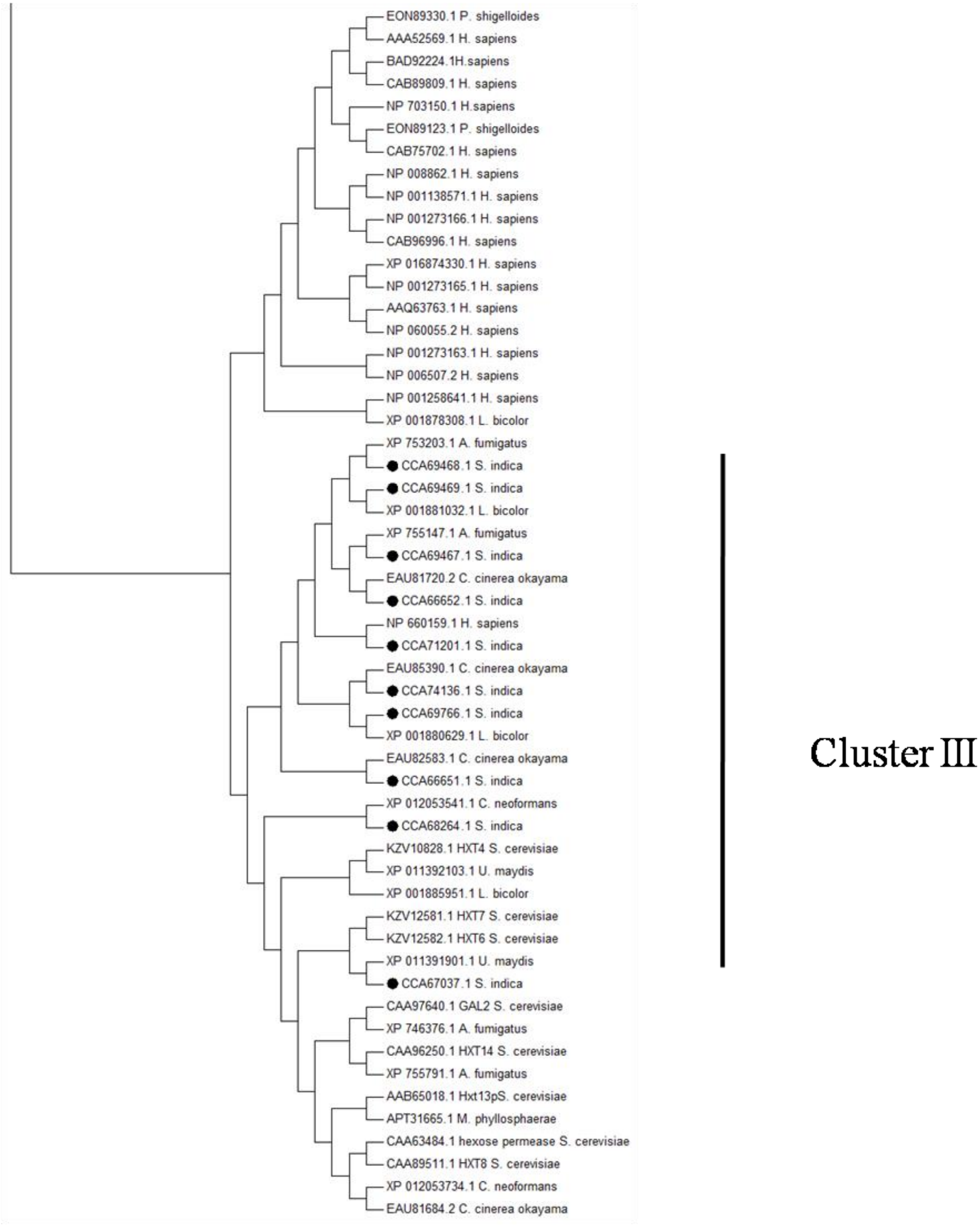


**Figure S9 (iii)**

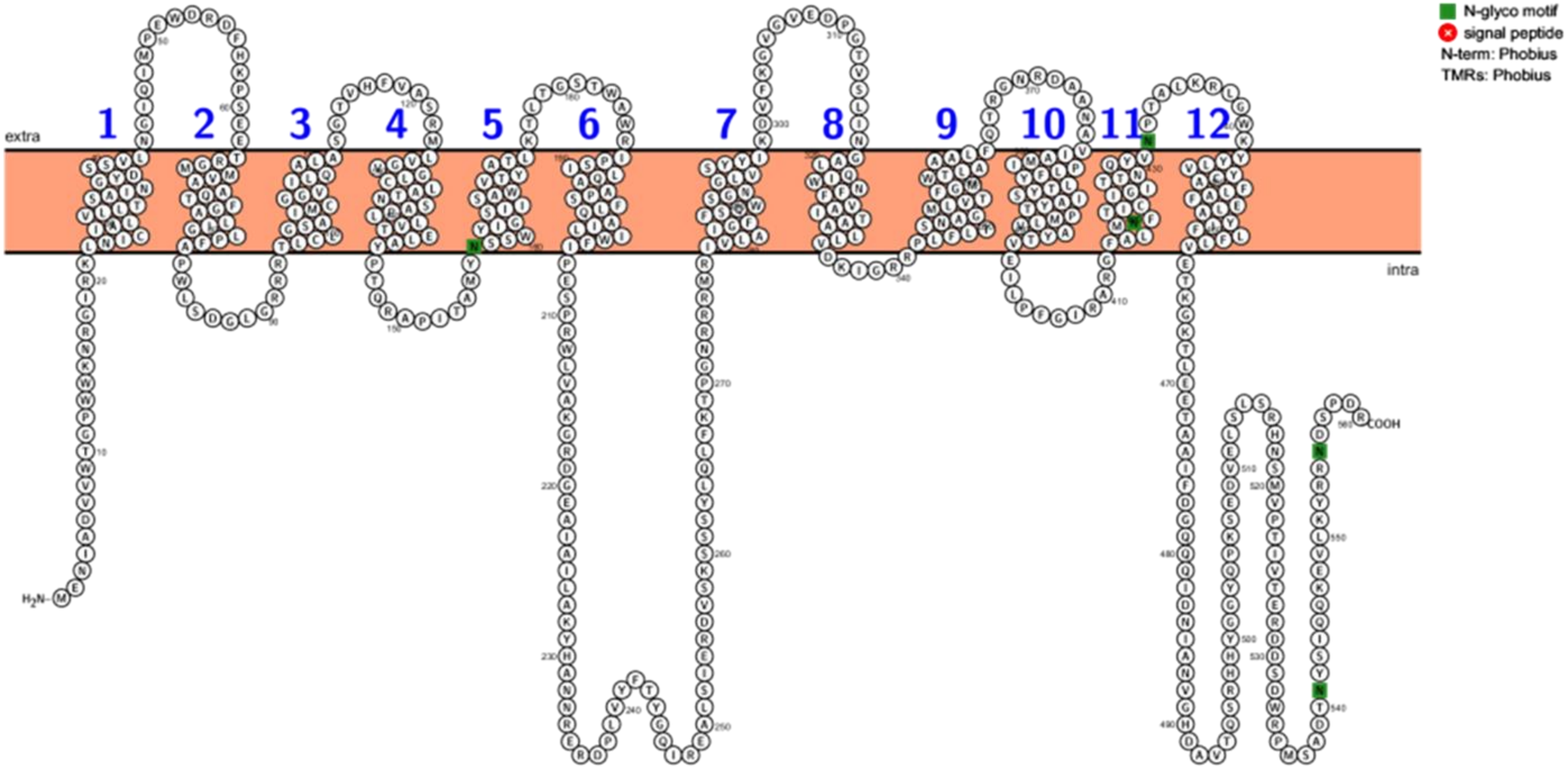


**PiST1**

**Figure S10 (i)**

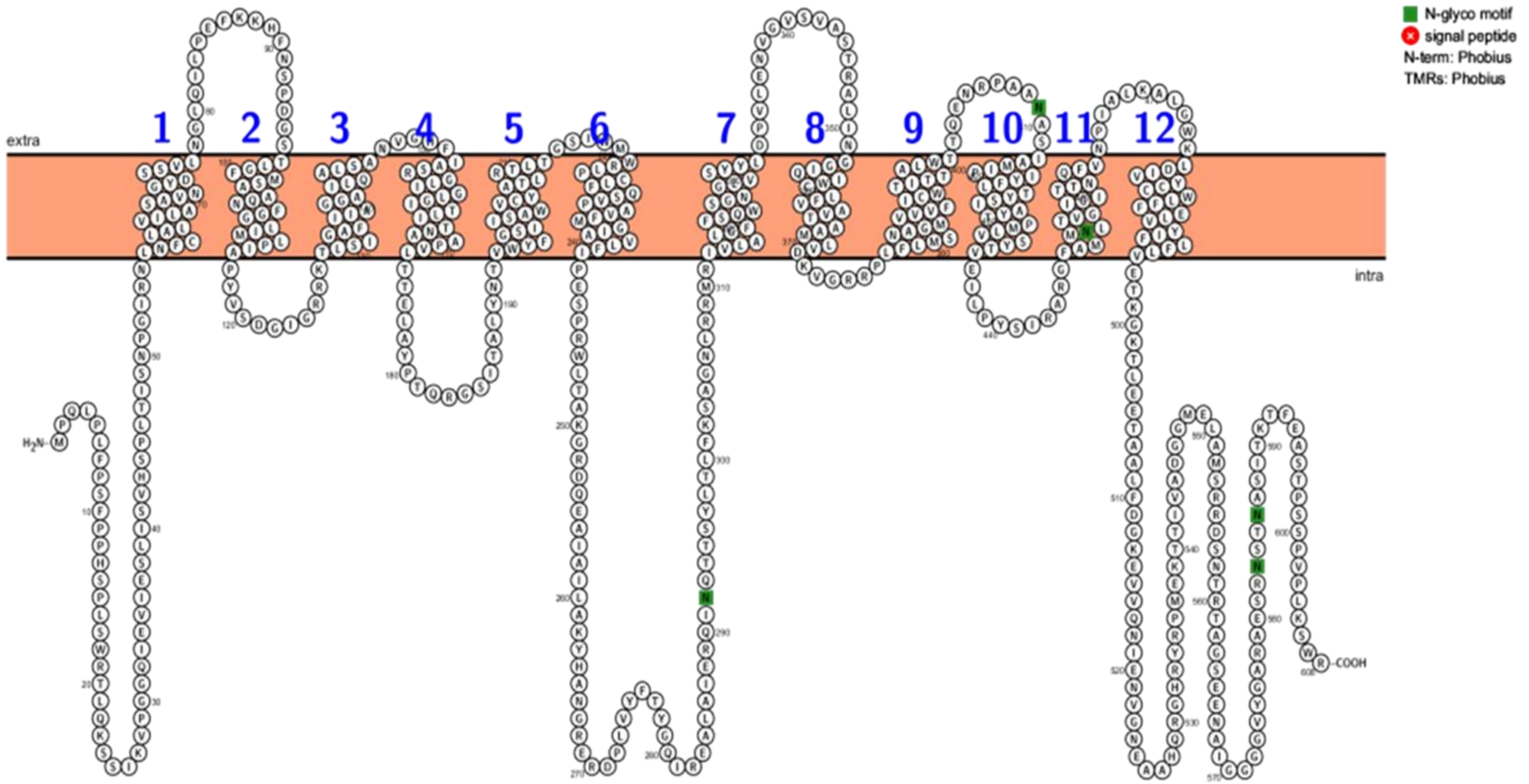


### PiST2

**Figure S10 (ii)**

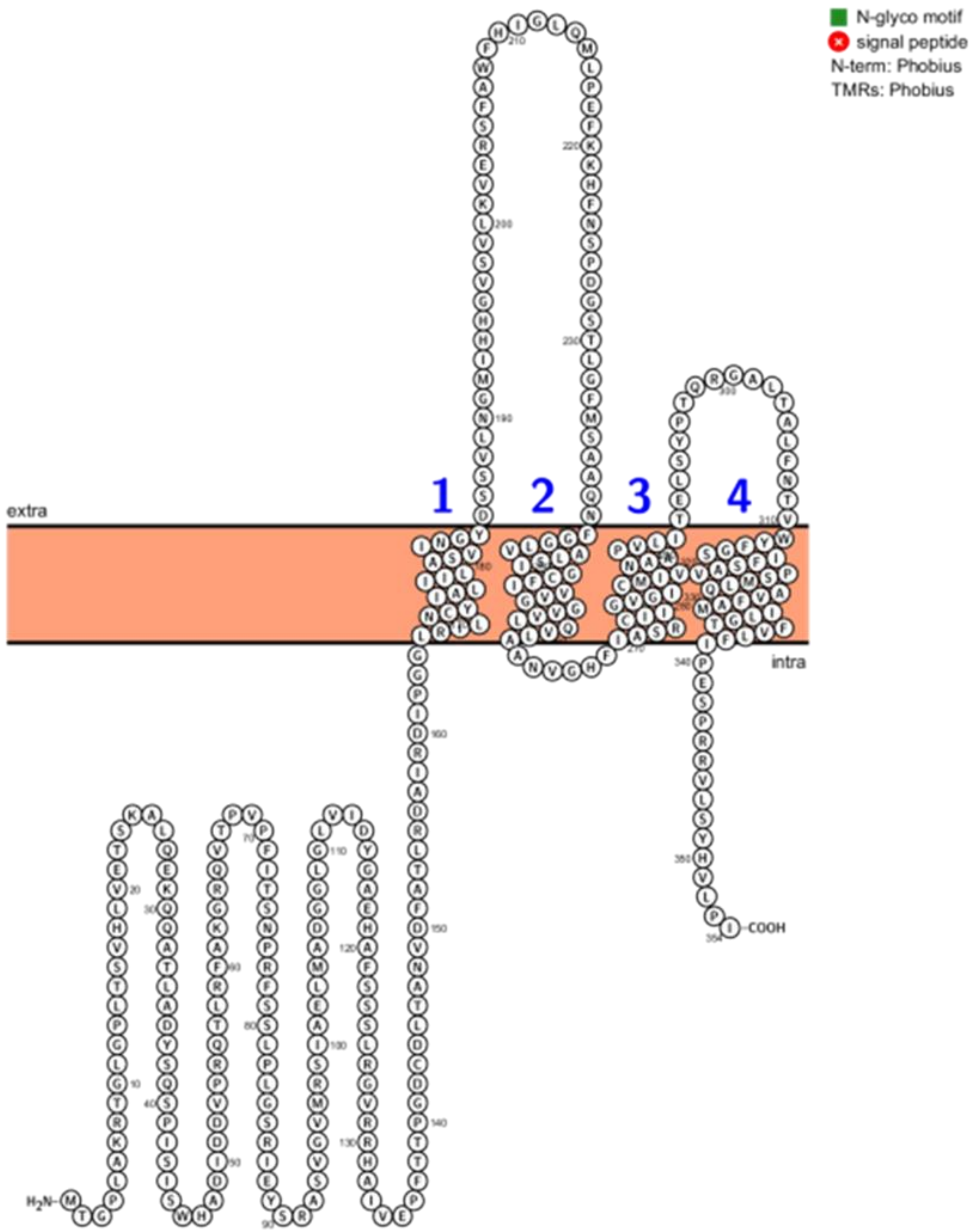


**PiST3**

**Figure S10 (iii)**

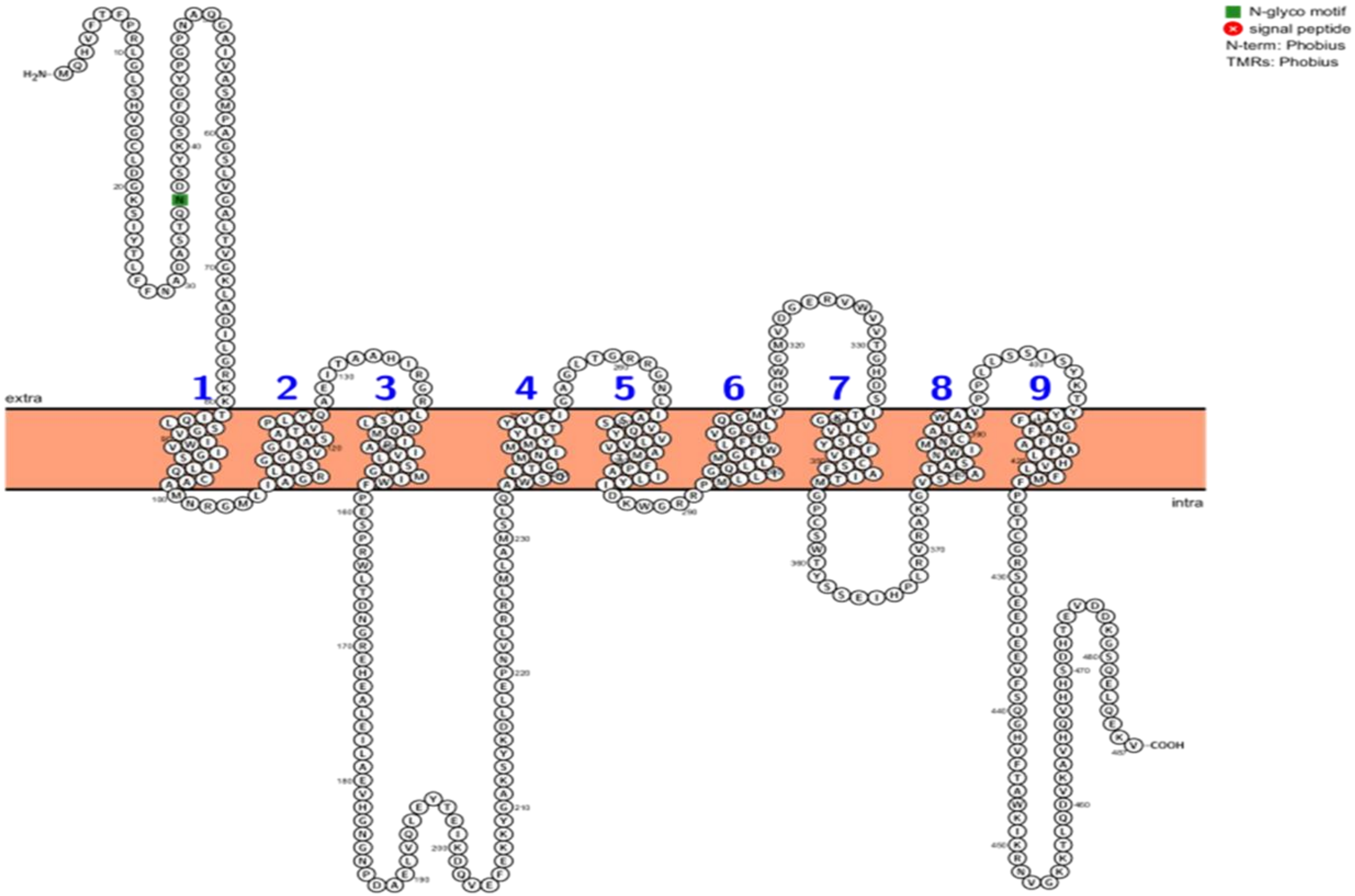


**PiST4**

**Figure S10 (iv)**

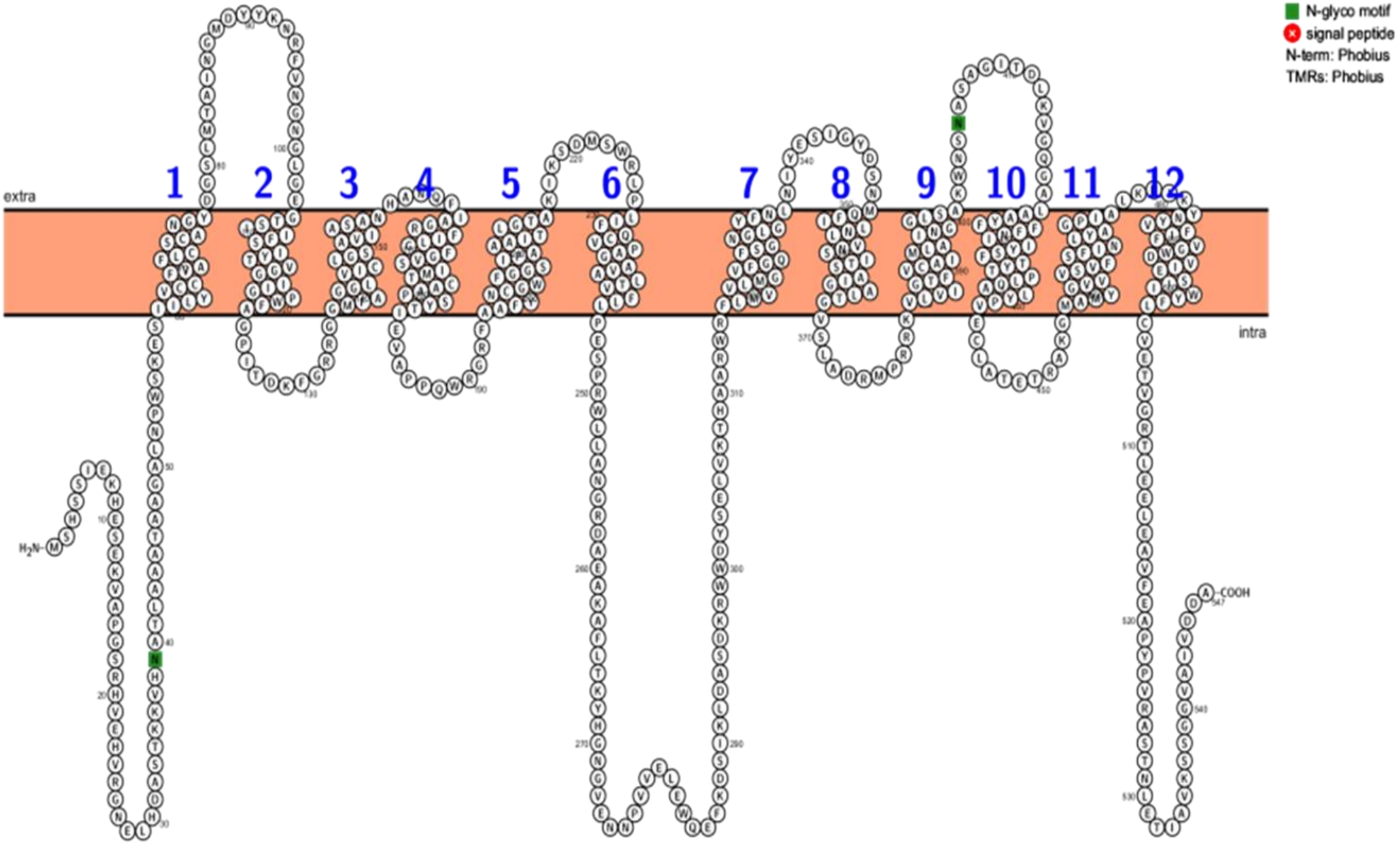


**PiST5**

**Figure S10 (v)**

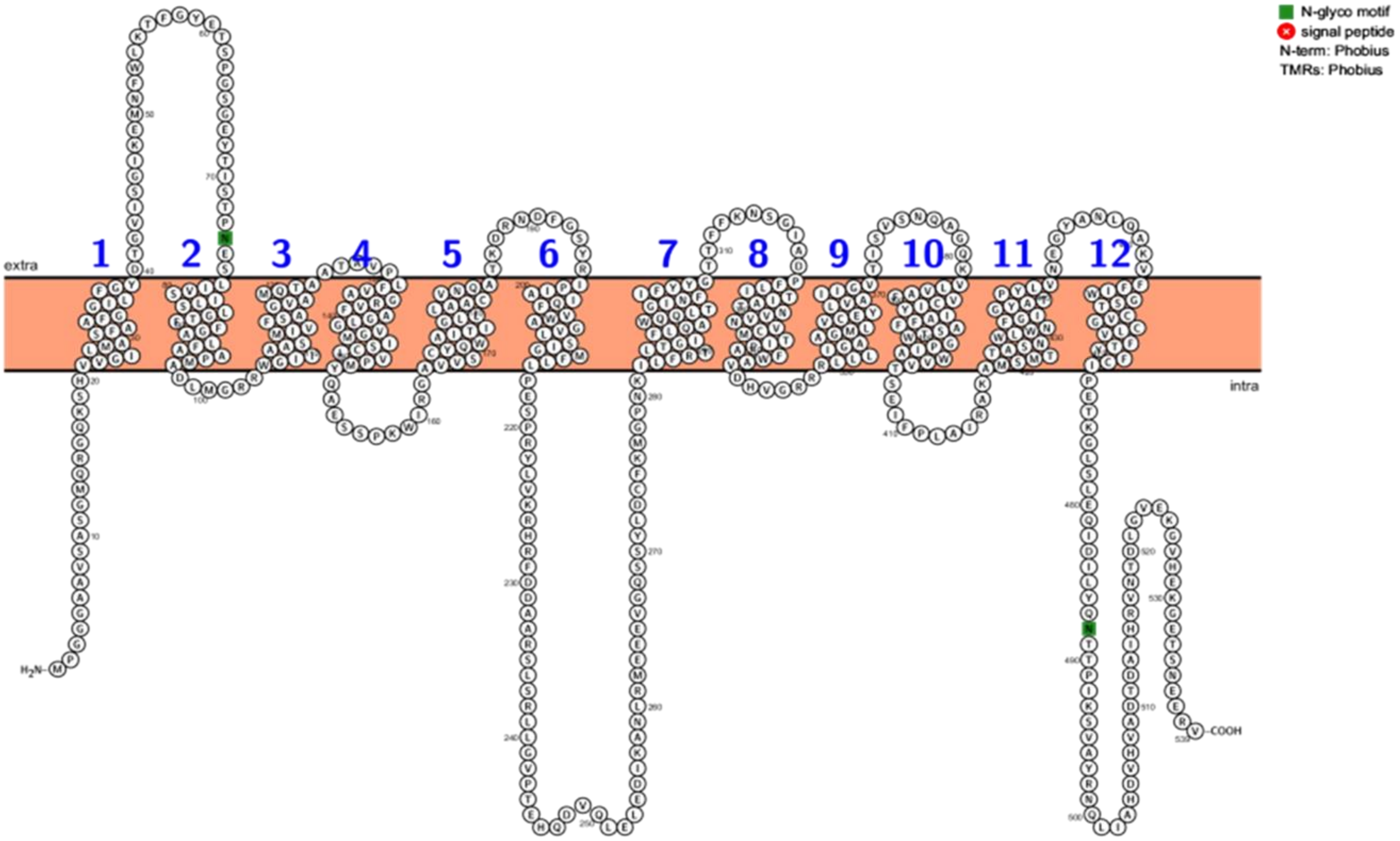


**PiST6**

**Figure S10 (vi)**

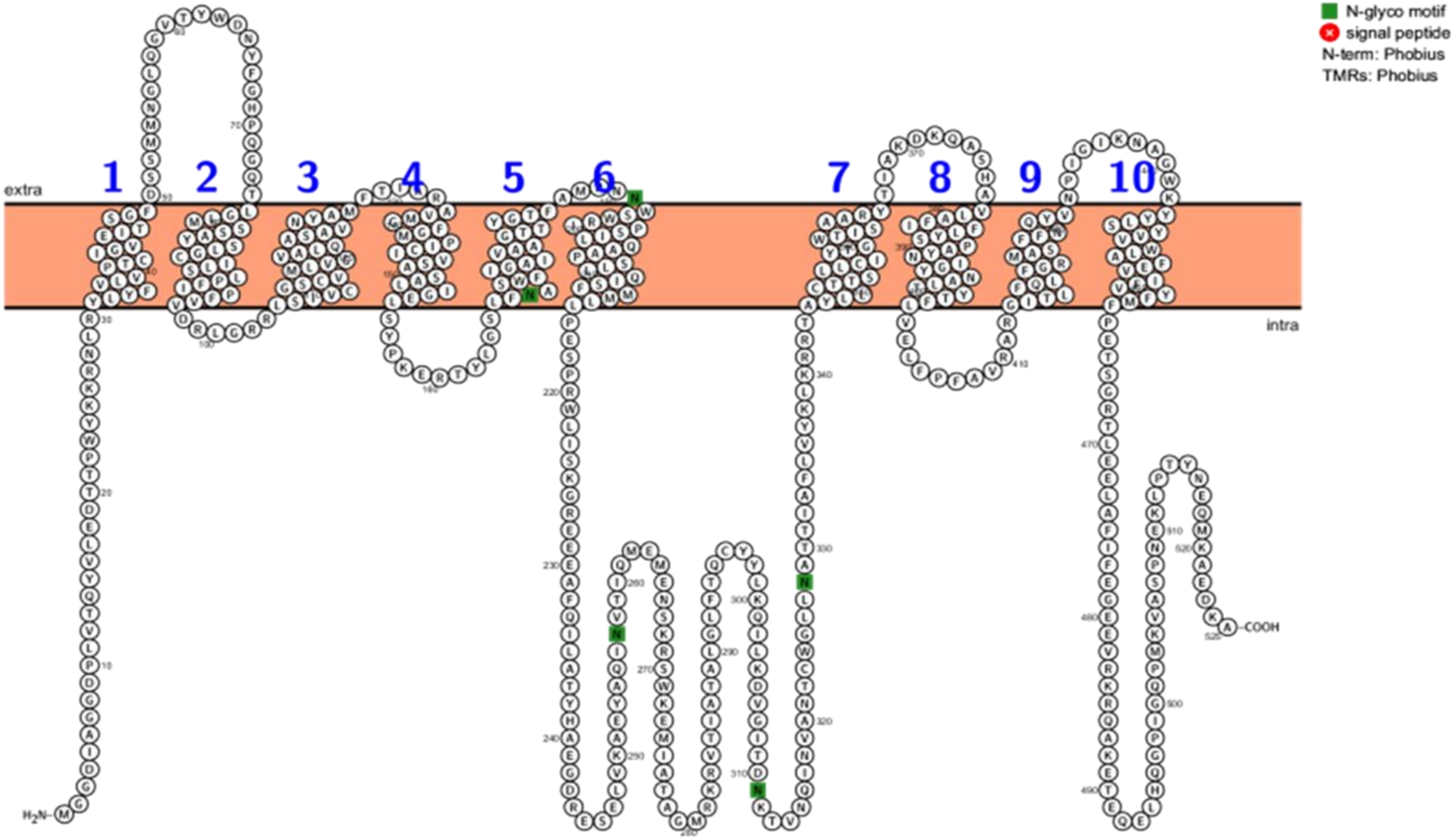


### PiST7

**Figure S10 (vii)**

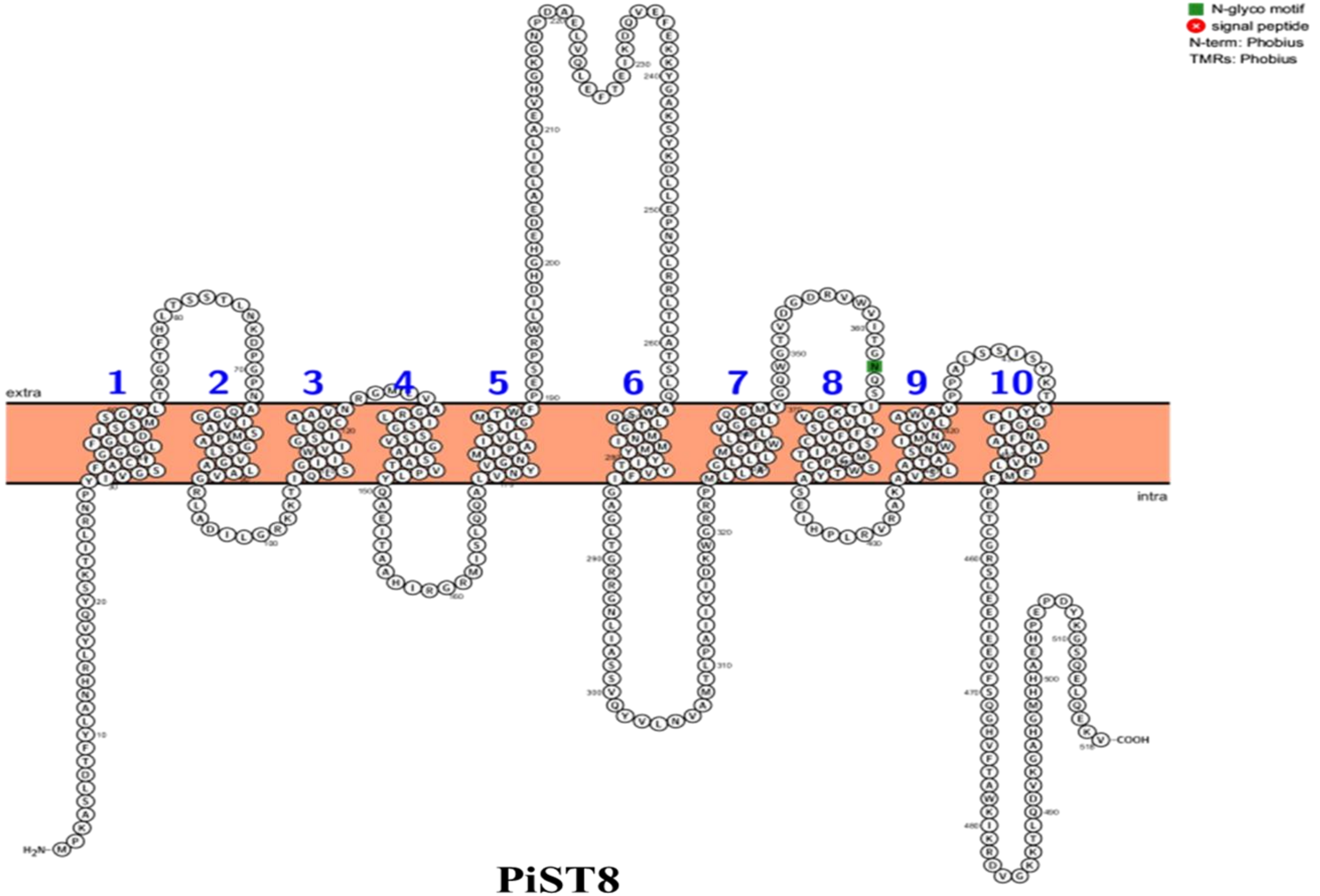


### PiST8

**Figure S10 (viii)**

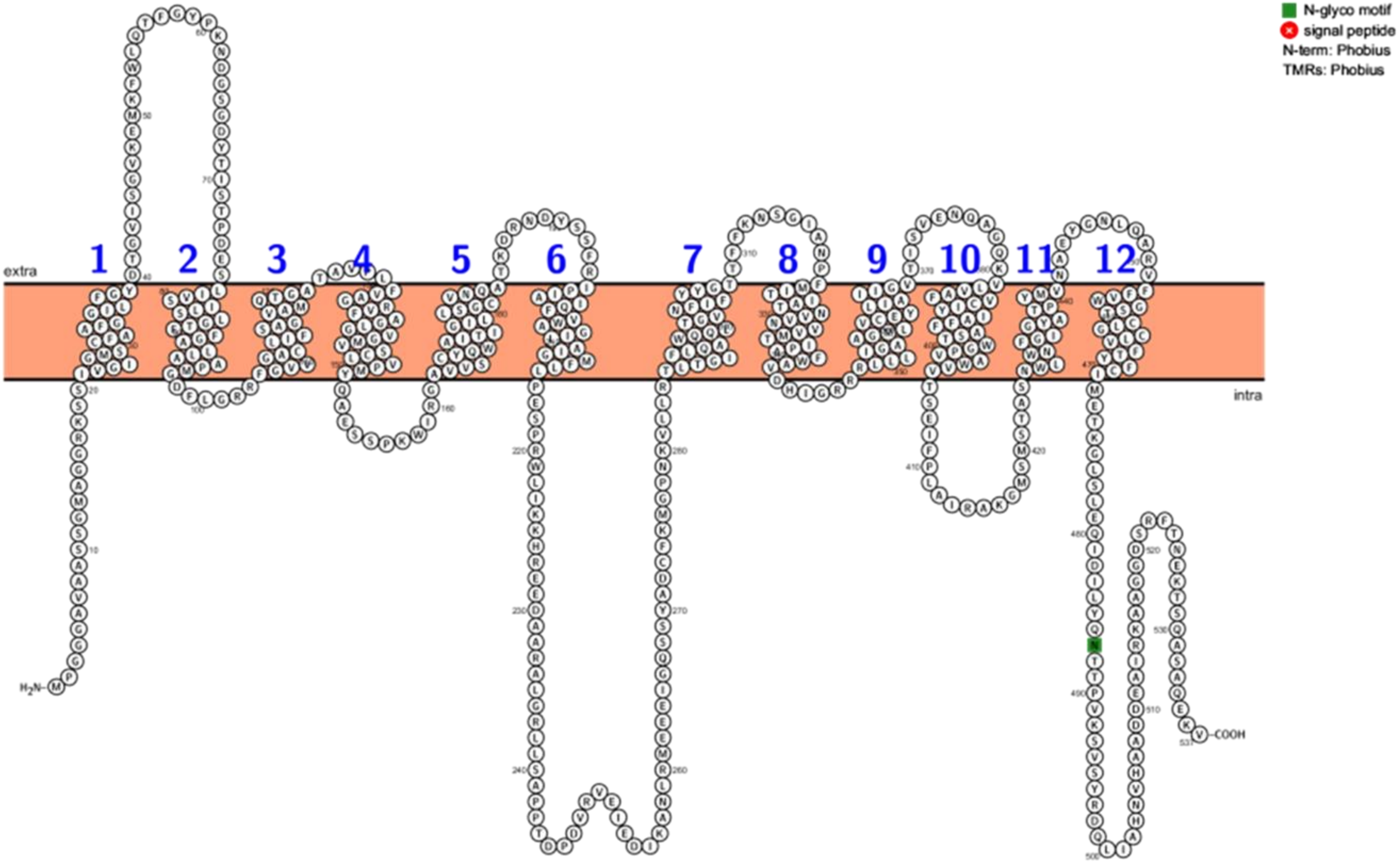


**PiST9**

**Figure S10 (ix)**

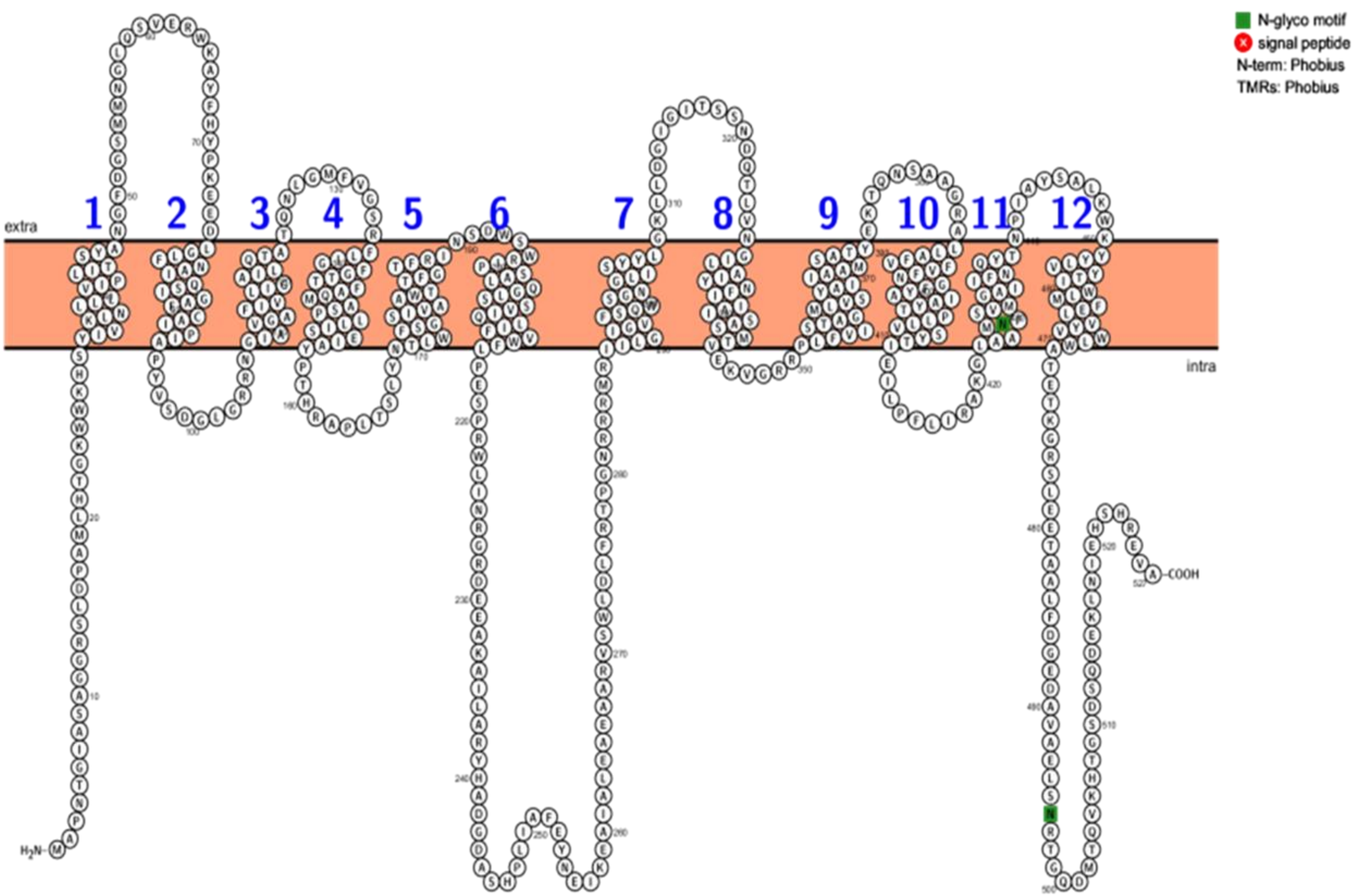


**PiST10**

**Figure S10 (x)**

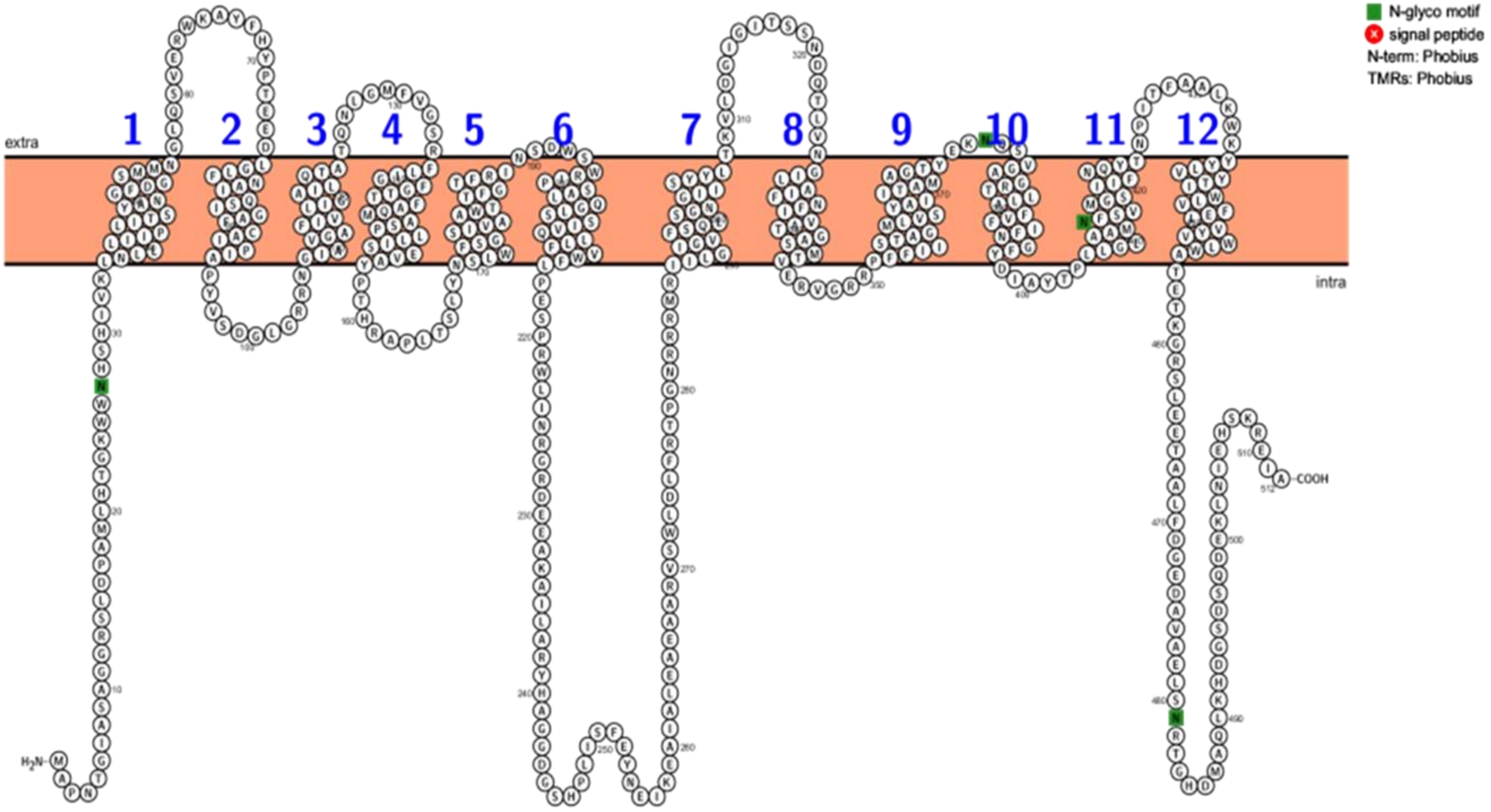


**PiST11**

**Figure S10 (xi)**

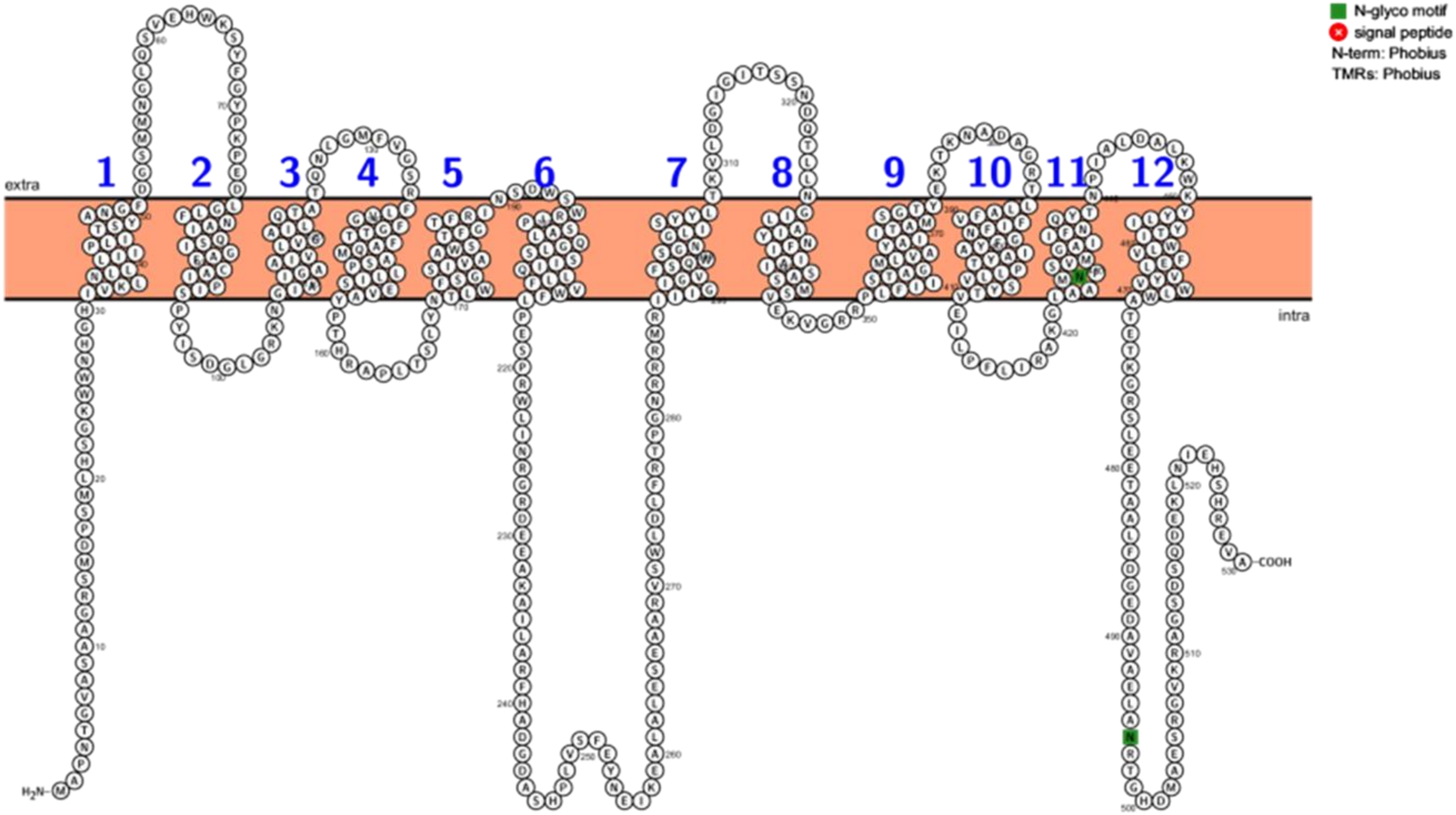


**PiST12**

Figure S10 (xii)

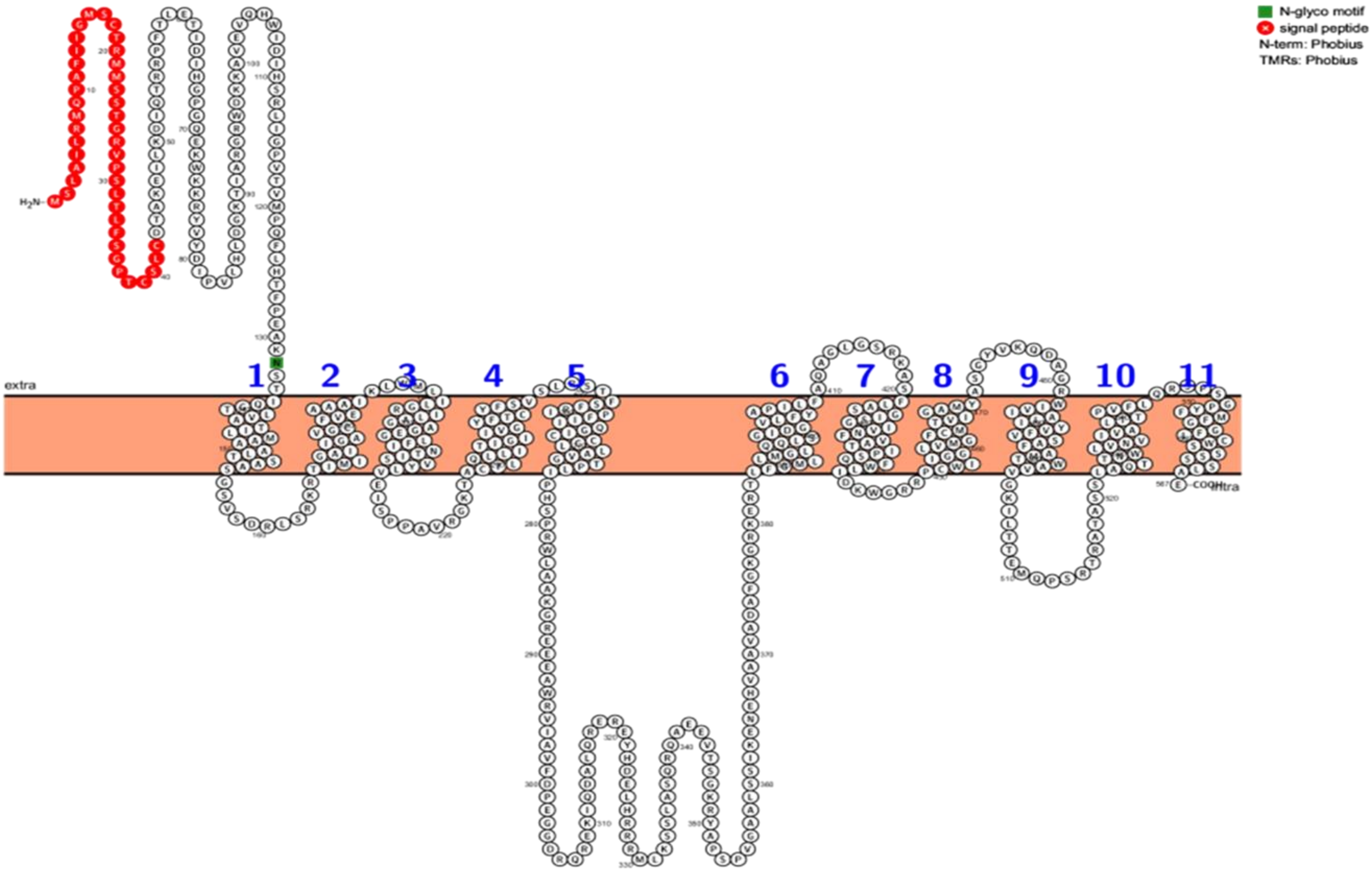


**PiST13**

**Figure S10 (xiii)**

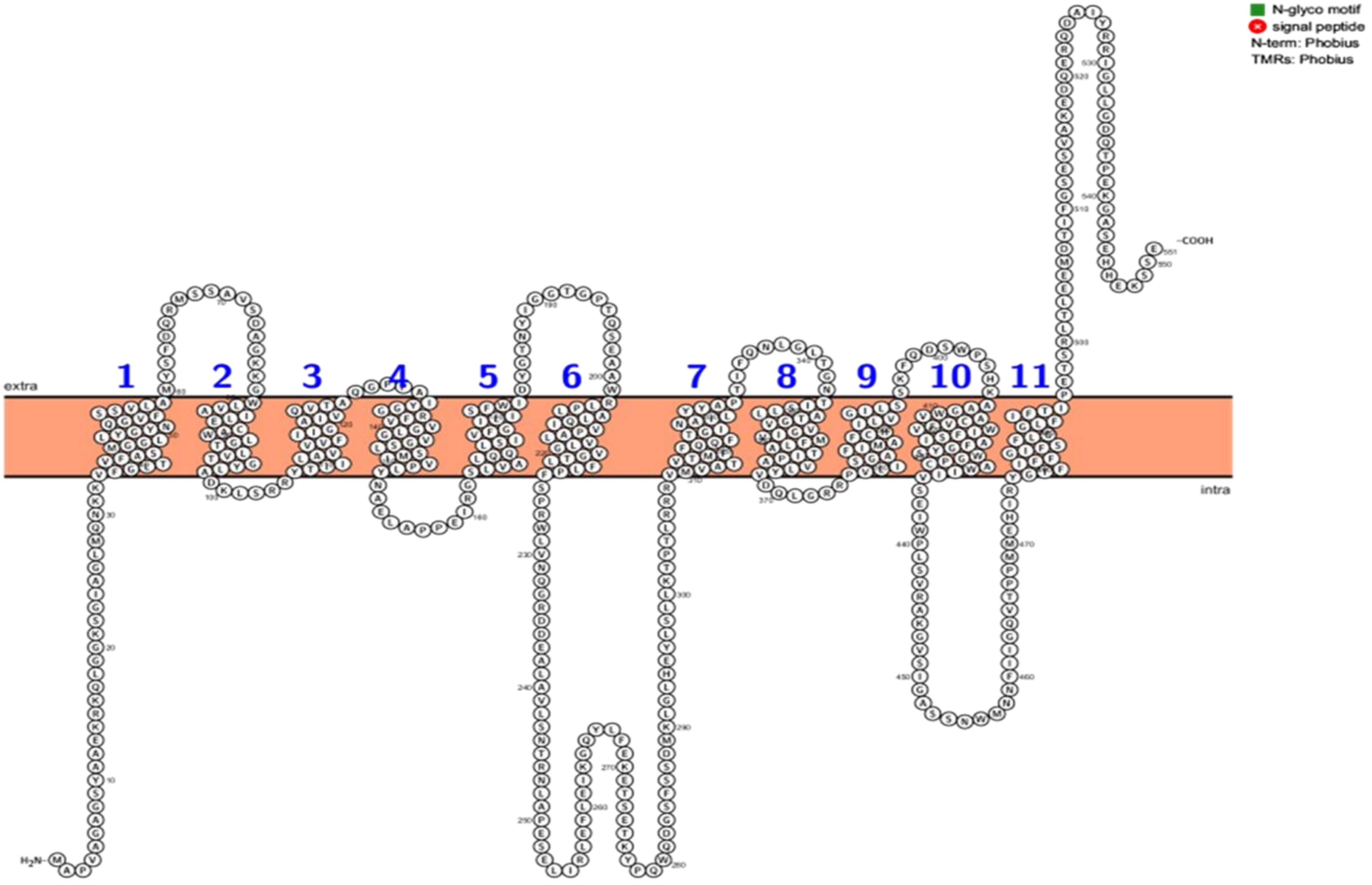


**PiST14**



**Figure S10 (xiv)**

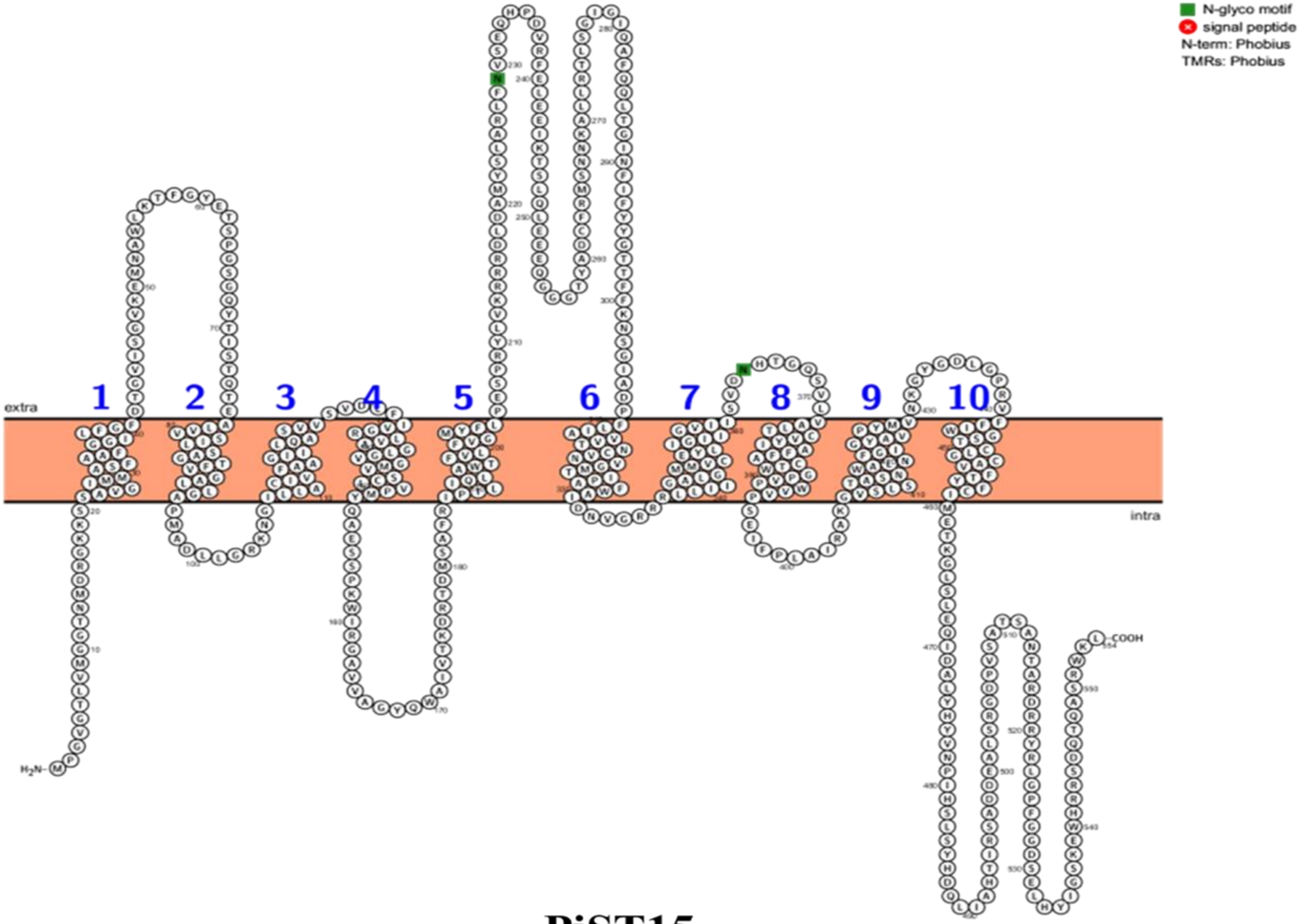


**PiST15**

**Figure S10 (xv)**

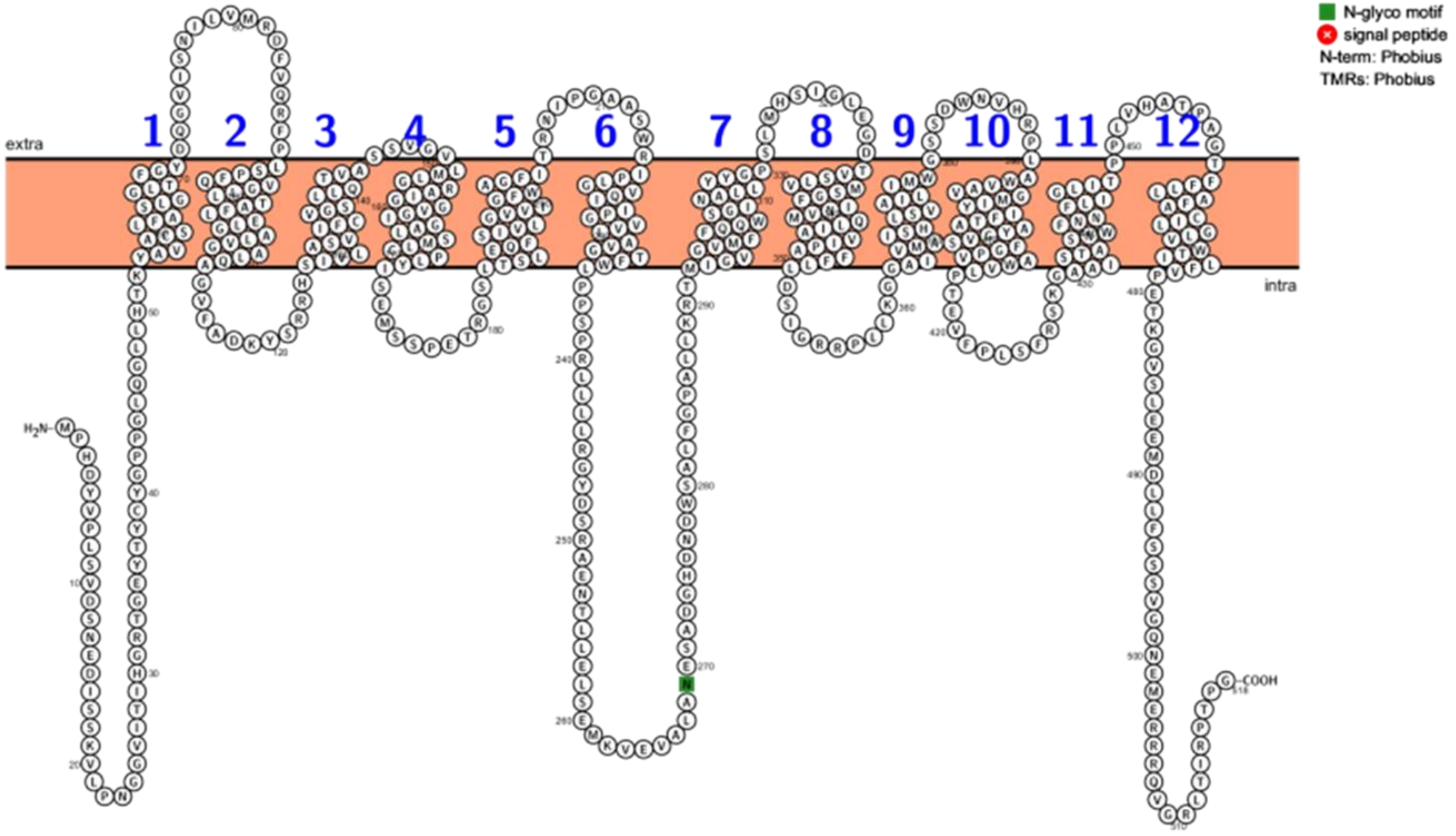


### PiST16

**Figure S10 (xvi)**

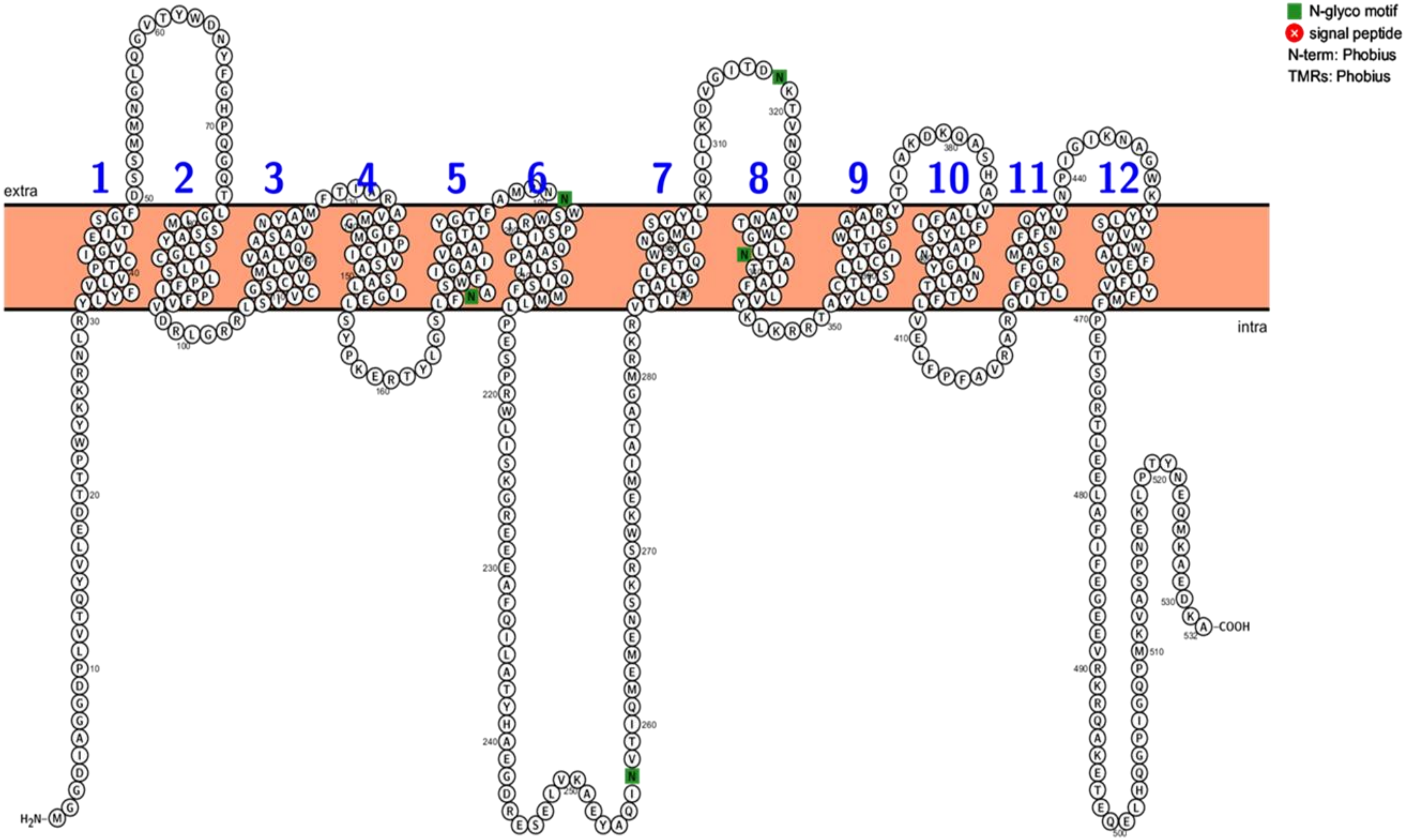


### PiST17

**Figure S10 (xvii)**

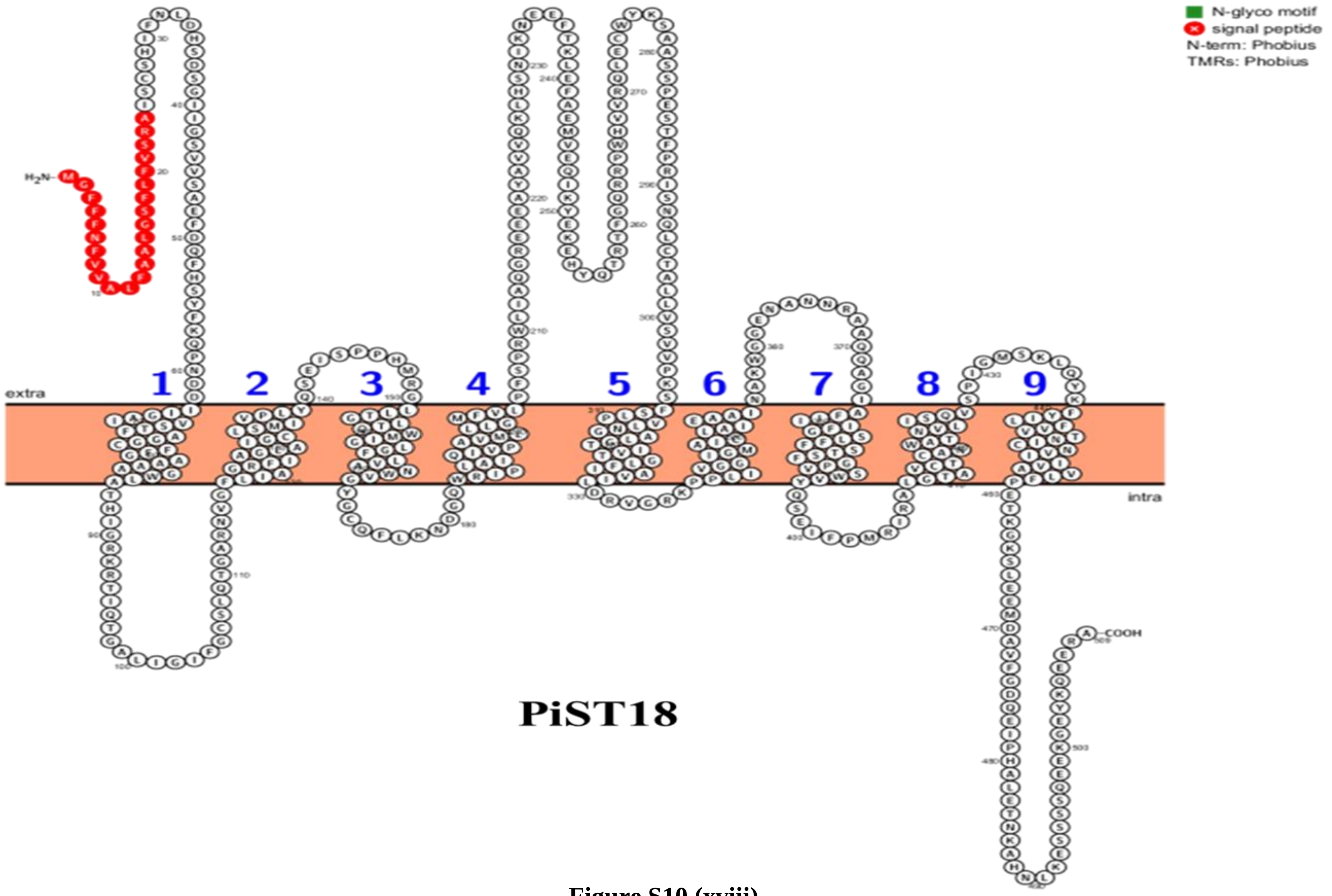


Figure S10 (xviii)

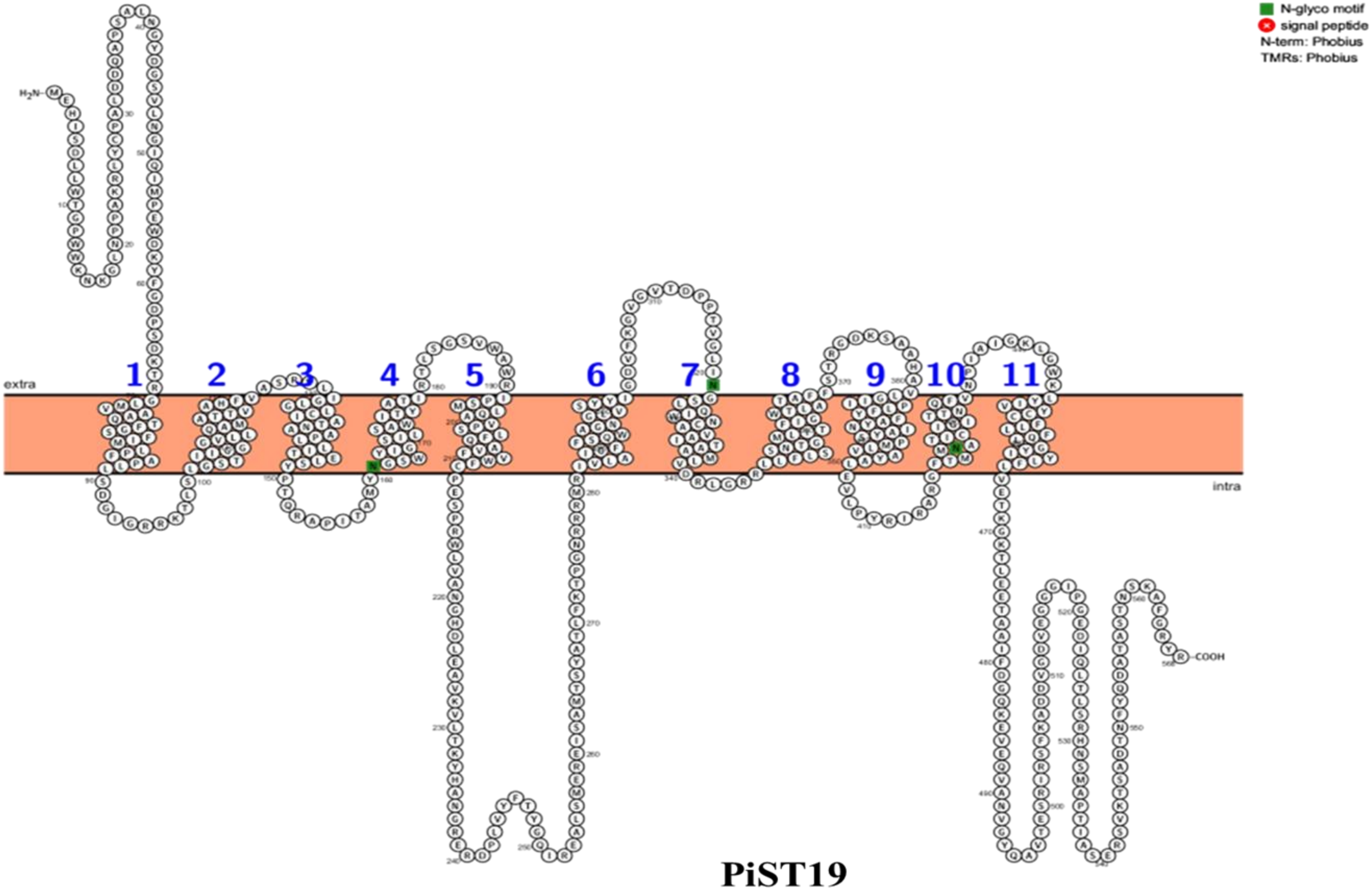


**PiST19**

**Figure S10 (xix)**

| **Table. S1: Details of Sequence used in Phylogenetic Analysis of Sugar Porter Family of *P. indica*** | | | |
|---|---|---|---|
| **Sr. No.** | **Accession No.** | **Organism** | **Protein Length** |
| **1** | CAA63484.1 | *Hexose permease S. cerevisiae* | 567 |
| **2** | AAB65018.1 | *Hxt13pS. cerevisiae* | 564 |
| **3** | CAA97640.1 | *GAL2 S. cerevisiae* | 574 |
| **4** | CAA89511.1 | *HXT8 S. cerevisiae* | 569 |
| **5** | KZV12581.1 | *HXT7 S. cerevisiae* | 570 |
| **6** | KZV12582.1 | *HXT6 S. cerevisiae* | 570 |
| **7** | KZV10828.1 | *HXT4 S. cerevisiae* | 576 |
| **8** | KZV10830.1 | *HXT1 S. cerevisiae* | 563 |
| **9** | KZV08856.1 | *HXT2 S. cerevisiae* | 541 |
| **10** | KZV12584.1 | *HXT3 S. cerevisiae* | 567 |
| **11** | ONH76975.1 | *HXT13 S. cerevisiae* | 318 |
| **12** | KZV10443.1 | *HXT12 S. cerevisiae* | 457 |
| **13** | CAA99178.1 | *HXT11 S. cerevisiae* | 567 |
| **14** | KZV11590.1 | *HXT10 S. cerevisiae* | 546 |
| **15** | CAA89516.1 | *HXT9 S. cerevisiae* | 567 |
| **16** | KZV10831.1 | *HXT5 S cerevisiae* | 592 |
| **17** | CAA96250.1 | *HXT14 S. cerevisiae* | 540 |
| **18** | KZV10448.1 | *SUC2 S. cerevisiae* | 532 |
| **19** | AAL89826.1 | *A. Niger* | 514 |
| **20** | XP 748226.1 | *A. fumigatus* | 559 |
| **21** | XP 746835.1 | *A. fumigatus* | 545 |
| **22** | AAL89824.1 | *A. niger* | 583 |
| **23** | XP 663464.1 | *A. nidulans* | 562 |
| **24** | XP 755484.1 | *A. fumigatus* | 560 |
| **25** | XP 001838247.1 | *C. cinereaokayama* | 610 |
| **26** | NP 033994.1 | *M. musculus* | 884 |
| **27** | XP 012051260.1 | *C. neoformans* | 520 |

| | | | |
|---|---|---|---|
| **28** | AJV31630.1 | *S. cerevisiae* | 570 |
| **29** | CAC41332.1 | *U. viciae-fabae* | 522 |
| **30** | XP 012052306.1 | *C. neoformans* | 428 |
| **31** | XP 018212773.1 | *O. Polymorpha* | 471 |
| **32** | XP 755791.1 | *A. fumigatus* | 530 |
| **33** | XP 746376.1 | *A. fumigatus* | 486 |
| **34** | OAP09858.1 | *A. thaliana* | 173 |
| **35** | CAA98771.1 | *S. cerevisiae* | 884 |
| **36** | CAA65621.1 | *RGT2 S. cerevisiae* | 763 |
| **37** | FR93278.2 | *AC. neoformans* | 551 |
| **38** | AAL89822.1 | *A. niger* | 530 |
| **39** | AFR96248.1 | *C. neoformans* | 563 |
| **40** | AFR96943.1 | *C. neoformans* | 565 |
| **41** | AFR98493.1 | *C. neoformans* | 550 |
| **42** | AFR96303.1 | *C. neoformans* | 538 |
| **43** | AFR96311.2 | *C. neoformans* | 543 |
| **44** | APT31665.1 | *M. phyllosphaerae* | 324 |
| **45** | EON89330.1 | *P. shigelloides* | 198 |
| **46** | APT30767.1 | *M. phyllosphaerae* | 408 |
| **47** | XP 012048363.1 | *C. neoformans* | 555 |
| **48** | EON89123.1 | *P. shigelloides* | 198 |
| **49** | XP 751523.1 | *A. fumigatus* | 548 |
| **50** | APT30947.1 | *M. phyllosphaerae* | 369 |
| **51** | XP 748233.1 | *A. fumigatus* | 498 |
| **52** | AFR97888.2 | *C. neoformans* | 600 |
| **53** | CAA59996.1 | *S. pombe* | 433 |
| **54** | AAL89825.1 | *A. niger* | 527 |
| **55** | XP 753575.1 | *A. fumigatus* | 532 |
| **56** | AFR95279.1 | *C. neoformans* | 539 |
| **57** | XP 012051959.1 | *C. neoformans* | 515 |
| **58** | XP 012051958.1 | *C. neoformans* | 512 |

| | | | |
|---|---|---|---|
| **59** | XP 749346.1 | *A. fumigatus* | 559 |
| **60** | XP 746324.1 | *A. fumigatus* | 531 |
| **61** | EKV51641.1 | *A. bisporus* | 1021 |
| **62** | XP 012052468.1 | *C. neoformans* | 595 |
| **63** | XP 750516.1 | *A. fumigatus* | 580 |
| **64** | AIE67034.1 | *K. michiganensis* | 192 |
| **65** | XP 001835254.1 | *C. cinereaokayama* | 532 |
| **66** | AY585934.1 | *P. involutus* | 1817 |
| **67** | XP 753203.1 | *A. fumigatus* | 531 |
| **68** | 755147.1 | *XP A. fumigatus* | 501 |
| **69** | P 749649.1 | *XA. fumigatus* | 557 |
| **70** | XP 012053734.1 | *C. neoformans* | 558 |
| **71** | XP 012053541.1 | *C. neoformans* | 540 |
| **72** | AFR98131.2 | *C. neoformans* | 545 |
| **73** | XP 012050040.1 | *C. neoformans* | 545 |
| **74** | XP 012047633.1 | *C. neoformans* | 535 |
| **75** | EON90036.1 | *P. shigelloides* | 185 |
| **76** | AFR99098.2 | *C. neoformans* | 462 |
| **77** | AFR98808.1 | *C. neoformans* | 560 |
| **78** | AFR93524.2 | *C. neoformans* | 459 |
| **79** | AFR99093.1 | *C. neoformans* | 631 |
| **80** | AFR93410.1 | *C. neoformans* | 678 |
| **81** | NP 001269793.1 | *H. sapiens* | 535 |
| **82** | CDI57695.1 | *L. helveticus* | 525 |
| **83** | NP 008862.1 | *H. sapiens* | 496 |
| **84** | XP 011536149.1 | *H. sapiens* | 606 |
| **85** | NP 703150.1 | *H.sapiens* | 520 |
| **86** | XP 016874330.1 | *H. sapiens* | 521 |
| **87** | NP 001273166.1 | *H. sapiens* | 535 |
| **88** | NP 001273163.1 | *H. sapiens* | 497 |
| **89** | AD92224.1H | *B.sapiens* | 517 |

| | | | |
|---|---|---|---|
| **90** | NP 006507.2 | *H. sapiens* | 492 |
| **91** | NP 001273165.1 | *H. sapiens* | 411 |
| **92** | AAI17118.1 | *H. sapiens* | 629 |
| **93** | CAB89809.1 | *H. sapiens* | 477 |
| **94** | CAB75702.1 | *H. sapiens* | 477 |
| **95** | EAW92683.1 | *H. sapiens* | 511 |
| **96** | AAH18897.1 | *H. sapiens* | 511 |
| **97** | AAA52569.1 | *H. sapiens* | 505 |
| **98** | NP 060055.2 | *H. sapiens* | 507 |
| **99** | CAB96996.1 | *H. sapiens* | 507 |
| **100** | NP 001258641.1 | *H. sapiens* | 314 |
| **101** | NP 001138571.1 | *H. sapiens* | 445 |
| **101** | EAW57811.1 | *H. sapiens* | 551 |
| **102** | NP 997303.2 | *H. sapiens* | 512 |
| **103** | AAH94735.1 | *H. sapiens* | 503 |
| **104** | NP 660159.1 | *H. sapiens* | 617 |
| **105** | EAW71608.1 | *H. sapiens* | 524 |
| **106** | EAW59618.1 | *H. sapiens* | 503 |
| **107** | NP 001315550.1 | *H. sapiens* | 354 |
| **108** | NP 060890.2 | *H. sapiens* | 547 |
| **109** | NP 001275979.1 | *H. sapiens* | 302 |
| **110** | ALQ34158.1 | *H. sapiens* | 435 |
| **111** | EFI28665.1 | *C. cinereaokayama* | 495 |
| **112** | EAU90547.2 | *C. cinereaokayama* | 502 |
| **113** | EAU86601.1 | *C. cinereaokayama* | 532 |
| **114** | EAU81684.2 | *C. cinereaokayama* | 537 |
| **115** | EAU90475.1 | *C. cinereaokayama* | 570 |
| **116** | EAU83615.1 | *C. cinereaokayama* | 553 |
| **117** | EAU81720.2 | *C. cinereaokayama* | 526 |
| **118** | EAU82583.1 | *C. cinereaokayama* | 530 |
| **119** | EAU81614.1 | *C. cinereaokayama* | 533 |

| | | | |
|---|---|---|---|
| **120** | EAU85390.1 | *C. cinereaokayama* | 610 |
| **121** | EFI27205.1 | *C. cinereaokayama* | 525 |
| **122** | XP 011386723.1 | *U. maydis* | 522 |
| **123** | XP 011391901.1 | *U. maydis* | 572 |
| **125** | XP 011387640.1 | *U. maydis* | 526 |
| **126** | XP 011391314.1 | *U. maydis* | 624 |
| **127** | XP 011390556.1 | *U. maydis* | 532 |
| **128** | XP 011388425.1 | *U. maydis* | 615 |
| **129** | XP 011388799.1 | *U. maydis* | 645 |
| **130** | P 011392103.1 | *XU. maydis* | 537 |
| **131** | XP 011391149.1 | *U. maydis* | 570 |
| **132** | XP 011392285.1 | *U. maydis* | 534 |
| **133** | XP 011392296.1 | *U. maydis* | 545 |
| **134** | CCA66651.1 | *S. indica* | 539 |
| **135** | CCA66652.1 | *S. indica* | 554 |
| **136** | CCA66654.1 | *S. indica* | 543 |
| **137** | CCA67037.1 | *S. indica* | 567 |
| **138** | CCA67897.1 | *S. indica* | 580 |
| **139** | CCA68264.1 | *S. indica* | 573 |
| **140** | CCA68995.1 | *S. indica* | 560 |
| **141** | CCA69467.1 | *S. indica* | 608 |
| **142** | CCA69468.1 | *S. indica* | 512 |
| **143** | CCA69469.1 | *S. indica* | 530 |
| **144** | CCA69766.1 | *S. indica* | 573 |
| **145** | CCA70504.1 | *S. indica* | 354 |
| **146** | CCA70963.1 | *S. indica* | 518 |
| **147** | CCA71201.1 | *S. indica* | 525 |
| **148** | CCA74136.1 | *S. indica* | 568 |
| **149** | CCA67930.1 | *S. indica* | 518 |
| **150** | CCA73450.1 | *S. indica* | 509 |
| **151** | CCA70422.1 | *S. indica* | 608 |

| | | | |
|---|---|---|---|
| **152** | XP 001885761.1 | *L. bicolor* | 489 |
| **153** | XP 001873946.1 | *L. bicolor* | 521 |
| **154** | XP 001885131.1 | *L. bicolor* | 517 |
| **155** | XP 001890363.1 | *L. bicolor* | 489 |
| **156** | XP 001885951.1 | *L. bicolor* | 511 |
| **157** | XP 001874568.1 | *L. bicolor* | 527 |
| **158** | XP 001888732.1 | *L. bicolor* | 529 |
| **159** | XP 001874382.1 | *L. bicolor* | 529 |
| **160** | XP 001873734.1 | *L. bicolor* | 554 |
| **161** | XP 001882059.1 | *L. bicolor* | 517 |
| **162** | XP 001877943.1 | *L. bicolor* | 526 |
| **163** | XP 001881032.1 | *L. bicolor* | 519 |
| **164** | XP 001880629.1 | *L. bicolor* | 553 |
| **165** | XP 001879389.1 | *L. bicolor* | 620 |
| **166** | XP 001878308.1 | *L. bicolor* | 455 |

**Table S2: Primers for expression analysis of Hexose Transporters from *P. indica***

| **Name** | **Sequence** | **Length** | **Tm** | **Length** |
|---|---|---|---|---|
| ***PiHXT1*** | For 5'ACAACTCGATGGTCCCAACCA3'<br>Rev 5'AGGACGATCGGGTGAGTCGTT3' | 21<br>21 | 58<br>59 | 134 bp |

| | | | | |
|---|---|---|---|---|
| ***PiHXT2*** | For 5’CGGAAAGCTCCTCGACGGTAT3’<br>Rev5`CAGCGGTCGAGATGACGAAGA 3’ | 21<br>21 | 59<br>60 | 155 bp |
| ***PiHXT3*** | For 5’GAGACTGCAGCCCTGTTCGAT 3’<br>Rev 5’ TCGCGGTGAGAGTGCTCAAT 3’ | 21<br>20 | 58<br>58 | 146 bp |
| ***PiHXT4*** | For 5’ ATCTCCGAGGTTGCCTACCCA 3’<br>Rev 5’ACGCCAGGACCAGTCACTGTT 3’ | 21<br>21 | 59<br>58 | 135 bp |
| ***PiHXT5*** | For 5’ATGGCCAGCTTCTTCAACCAA 3’<br>Rev 5’ CGCCAGCTCTTCCAGTGTTCT 3’ | 21<br>21 | 58<br>58 | 156 bp |
| ***PiHXT6*** | For 5’ GCATGGAGCTCGCAATGTCA 3’<br>Rev5’TGAAAGCAGAGGACAAGGCGT 3’ | 20<br>21 | 59<br>58 | 166 bp |
| ***PiHXT7*** | For 5’GGAGACCCGAGCAAAGGGTAT 3’<br>Rev 5’GATAACGTCCCATCCGACGAA 3’ | 21<br>21 | 58<br>58 | 139 bp |
| ***PiHXT8*** | For 5’ATGCCTGGTGGTGGTGCAGT 3’<br>Rev 5’ TTAGACCTTCTCTTGTGCACT 3’ | 20<br>21 | 59<br>60 | 155 bp |
| ***PiHXT9*** | For 5’ TCTCCGGTGTCAAGGAAATGA3’<br>Rev 5’AGAAGATGAGGCAAGCACCGA3’ | 21<br>21 | 60<br>60 | 160 bp |
| ***PiTef*** | For 5`-TCGTCGCTGTCAACAAGATG-3`<br>Rev 5`-GAGGGCTCGAGCATGTTGT-3’ | 20<br>19 | 59<br>59 | 135 bp |

**Table S3: Primers used in cloning of *PiST12* and its sequencing**

| **Sr. No.** | **Primer Name** | **Primer Sequence** | **Purpose** |
|---|---|---|---|
| **1.** | PiHXT3-F | 5’ CAAGCTTGGATGGCTCCCAACACTGG TGT 3’ | For Cloning PiHXT3 |
| **2.** | PiHXT3-R | 5’-TCAGGCAACCTCGCGGTGAGAGT-3’ | |
| **3.** | PiHXT3-Fi | 5’-ACCTTTGCCCAGATGGCTTCT-3’ | Internal Primer for PiHXT3 cloning |
| **4.** | PiHXT3-Ri | 5’-AAGATGAAGACGAATGCGAGGA-3’ | |
| **10.** | HygroR | 5’-AAGATGTTGGCGACCTCGTATTG-3’ | |
| **11.** | M13For | 5’-CGCCAGGGTTTTCCCAGTCACGAC-3’ | Sequencing and colony PCR |
| **12.** | M13Rev | 5’-AGCGGATAACAATTTCACACAGGA-3’ | |
| **13.** | T7 Rev | 5’-TAATACGACTCACTATAGGG-3’ | Sequencing and colony PCR |
| **14.** | SP6 | 5’-ATTTAGGTGACACTATAG-3’ | |

**Table S4: Conserve domain analysis of putative PiST12. Amino acid sequence of putative PiST12 was analyzed by NCBI's Conserved Domain Database (CDD)**

| Name | Accession No. | Description | Interval | E-value |
|---|---|---|---|---|
| **Sugar-tr** | Pfam00083 | Sugar transporter | 44-485 | 1.09e-79 |
| **SP** | TIGR00879 | MFS transporter, sugar porter (SP) family; This model represents the sugar porter subfamily. | 77-481 | 1.66e-70 |
| **xylE** | PRK10077 | D-xylose transporter XylE | 49-484 | 1.74e-30 |
| **MFS** | cd06174 | The Major Facilatator Superfamily (MFS) is a large and diverse group of secondary transporters | 47-470 | 1.37e-17 |
| **FucP** | COG0738 | Fucose Permease (Fructose transport and metabolism) | 63-194 | 8.34e-04 |
